# Pseudovibriamides from *Pseudovibrio* marine sponge bacteria promote swarming motility via transcriptional modulation

**DOI:** 10.1101/2024.04.03.587961

**Authors:** Yitao Dai, Vitor Lourenzon, Laura P. Ióca, Dua Al-Smadi, Lydia Arnold, Ian McIntire, Roberto G. S. Berlinck, Alessandra S. Eustáquio

## Abstract

*Pseudovibrio* α-Proteobacteria have been repeatedly isolated from marine sponges and proposed to be beneficial to the host. Bacterial motility is known to contribute to host colonization. We have previously identified pseudovibriamides A and B, produced in culture by *Pseudovibrio brasiliensis* Ab134, and shown that pseudovibriamide A promotes flagellar motility. Pseudovibriamides are encoded in a hybrid nonribosomal peptide synthetase-polyketide synthase gene cluster that also includes several accessory genes. Pseudovibriamide A is a linear heptapeptide and pseudovibriamide B is a nonadepsipeptide derived from pseudovibriamide A. Here we define the borders of the pseudovibriamides gene cluster, assign function to biosynthetic genes using reverse genetics and test the hypothesis that pseudovibriamides impact motility by modulating gene transcription. RNA-seq transcriptomic analyses of strains having different compositions of pseudovibriamides suggested that both pseudovibriamides A and B affect genes potentially involved in motility, and that a compensatory mechanism is at play in mutants that produce only pseudovibriamide A, resulting in comparable swarming motility as the wild type. The data gathered suggest that pseudovibriamides A and B have opposite roles in modulating a subset of genes, with pseudovibriamide B having a primary effect in gene activation, and pseudovibriamide A on inhibition. Finally, we observed many differentially expressed genes (up to 29% of the total gene number) indicating that pseudovibriamides have a global effect on transcription that goes beyond motility.

**Importance:** Marine sponges are found throughout the oceans from tropical coral reefs to polar sea floors, playing crucial roles in marine ecosystems. *Pseudovibrio* bacteria have been proposed to contribute to sponge health. We have previously shown that pseudovibriamides produced by *Pseudovibrio brasiliensis* promote bacterial motility, a behavior that is beneficial to bacterial survival and to host colonization. The gene cluster that encodes pseudovibriamide biosynthesis is found in two thirds of *Pseudovibrio* genomes. This gene cluster is also present in *Pseudomonas* bacteria that interact with terrestrial plants and animals. Here we first assign function to pseudovibriamide biosynthetic genes using reverse genetics. We then show that pseudovibriamides play a major role in transcriptional regulation, affecting up to 29% of *P. brasiliensis* genes, including motility genes. Thus, this work gives insights into pseudovibriamide biosynthesis and provides evidence that they are signaling molecules relevant to bacterial motility and to other yet to be identified phenotypes.

## Introduction

Marine sponges are among the oldest animals on Earth (1). Their filter feeding capacity contributes to biogeochemical cycling and they are also involved in habitat formation, properties that are critical to marine ecology (2). In a bulk nutrient-depleted environment like the open ocean, colonization on marine sponges provides microbes greater access to nutritional resources and environmental stability (3). Conversely, sponge-associated microbes have nutritional and protective roles, contributing to the animal’s health (4).

Marine sponges develop symbiotic relationships with numerous bacterial species (5, 6). Among these, *Pseudovibrio* spp. are Gram-negative α-Proteobacteria with high frequency of association with marine sponges and that have been proposed to contribute to sponge health (7). The presence of *Pseudovibrio* was also confirmed in larvae of a marine sponge suggesting this genus could be a vertically transmitted symbiont (8).

Metabolites produced by microbes are important in the establishment and maintenance of host-microbe associations and in holobiont homeostasis. *Pseudovibrio* bacteria, for example, produce antimicrobials which could ward off pathogens (7). Moreover, swarming motility – the collective motion of bacterial cells across a solid surface powered by rotating flagella – has been shown to be important for host colonization (9). Bacterial metabolites such as surfactants are known to facilitate swarming motility (9).

We have previously reported a link between a biosynthetic gene cluster (BGC) in *Pseudovibrio brasiliensis* Ab134 and its swarming motility (10). Strain Ab134 was isolated from marine sponge *Arenosclera brasiliensis* (11). The BGC, which we termed *<u>P</u>seudovibrio* and *<u>P</u>seudomonas* non-ribosomal <u>p</u>eptide (*ppp*), is present in 2/3 of sequenced *Pseudovibrio* genomes, and sporadically found in *Pseudomonas* γ-Proteobacteria known to interact with terrestrial plants and insects (10). Moreover, a *ppp*-like BGC was recently reported from *Microbulbifer* γ-Proteobacteria likewise isolated from marine sponges (12).

The *ppp* BGC consists of genes *pppABC* encoding nonribosomal peptide synthetases (NRPS), *pppD* encoding a hybrid NRPS-polyketide synthase (NRPS-PKS), and multiple flanking genes *pppE to pppP* (**Fig. 1**; **Table S1**). NRPS *pppA::neo* mutants helped us identify the products of the *ppp* BGC, which we termed pseudovibriamides A, and B (PA and PB) (10). *P. brasiliensis* Ab134 also accumulates a third product, pseudovibriamide C (PC), detected here by mass spectrometry analysis. Impaired swarming motility was reported for *pppA::neo* mutants, but not for *pppD::neo* mutants, indicating that only PA is required for wild-type level motility (10).

**Figure 1.**
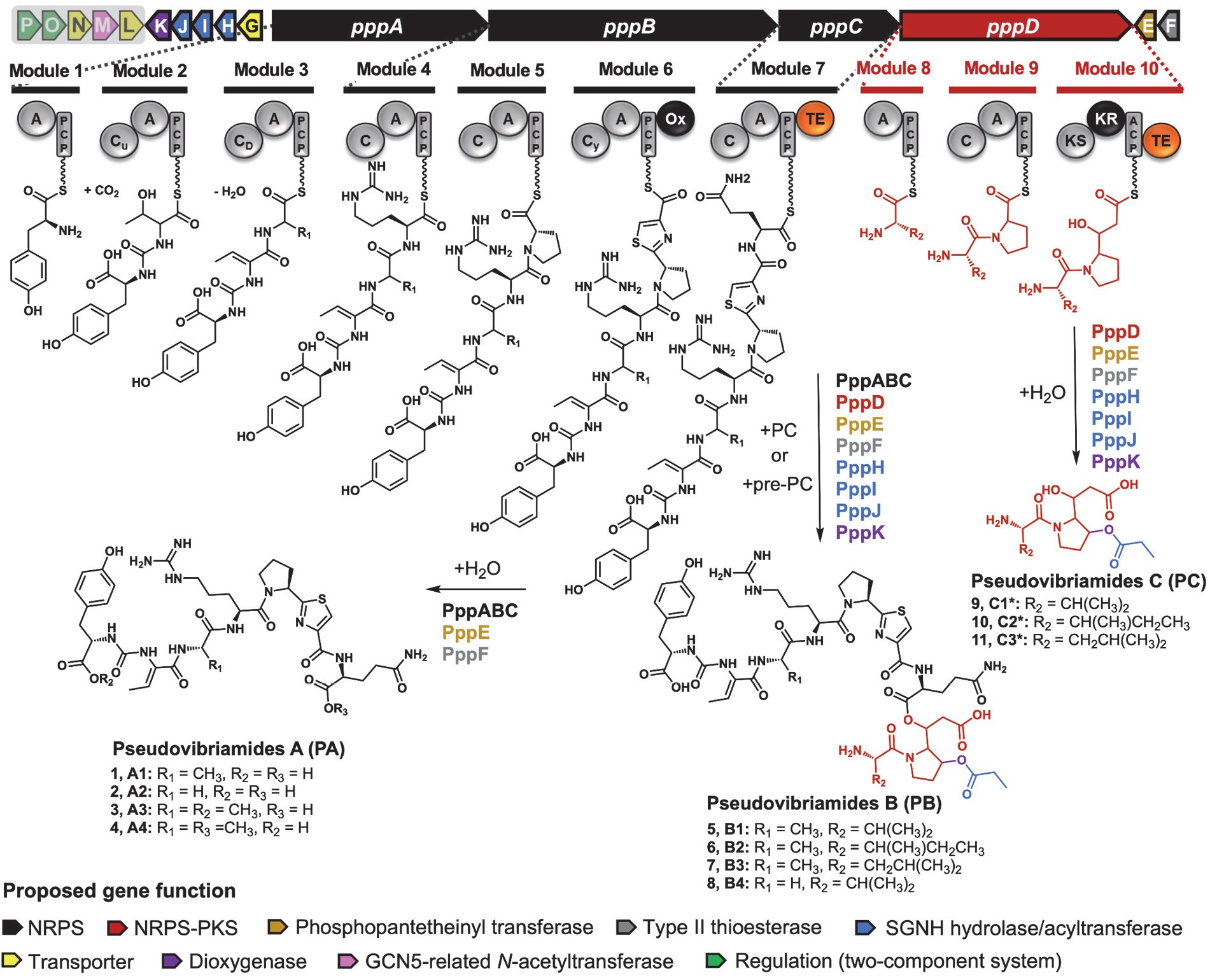
Organization of the *ppp* gene cluster from *Pseudovibrio brasiliensis* Ab134, and proposed biosynthesis of pseudovibriamides. **A, B, and C.** Biosynthetic proposal based on gene knockout results reported in this study. Structural components are color-coded according to genes that encode them. Enzymes proposed to be involved in the biosynthesis of each pseudovibriamide are listed next to the arrows. NRPS, nonribosomal peptide synthetase. PKS, polyketide synthase. See **Table S1** for further details on the Ppp proteins. Domain key: A, adenylation; ACP/PCP, acyl/peptidyl carrier protein; C, condensation; Cu, ureido-linkage formation condensation domain; CD, dehydration condensation domain; Cy, condensation and heterocyclization; KR, ketoreductase; KS, ketosynthase; Ox, oxidase; TE, thioesterase. Figure adapted from ref. (10). The structures of PA and PB have been previously reported (10). PC structures (*) are proposed based on mass spectrometry analyses performed in this study. Genes proposed to be outside the *ppp* gene cluster are shaded.

However, an understanding of how pseudovibriamides affect swarming motility is lacking. Swarming is a complex behavior that involves many factors, including signal transduction cascades like chemotaxis and quorum sensing, flagella which are driven by the proton motive force, and metabolites like surfactants (9). Most nonribosomal peptides that are known to mediate swarming are lipopeptide surfactants (13). Surfactants facilitate swarming by lowering the surface tension and easing cell spreading. Pseudovibriamides have no surfactant activity, as expected from their chemical structures (10). Alternatively, nonribosomal peptides, including those with a surfactant nature, can function as signaling molecules, exerting an effect on transcription and regulating various phenotypes (14–17).

Herein we report and discuss the results of a study aiming to gain insight into pseudovibriamide biosynthesis using reverse genetics, while obtaining mutants with different compositions of pseudovibriamides to ultimately test the hypothesis that pseudovibriamides act as signaling molecules, regulating genes affecting swarming motlity.

## Results

### Biosynthetic insights from mutagenesis studies and access to strains with different compositions of pseudovibriamides

Having mutants that produce only one of the pseudovibriamide congeners would facilitate probing their individual roles. According to the scheme presented in **Fig. 1**, PB is the full-length product. PA and PC could conceivably either represent hydrolysis products of PB catalyzed by an accessory hydrolase or be directly released from the assembly lines using water as the nucleophile in reactions catalyzed by thioesterase (TE) domains present in modules 7 and 10, respectively. To probe these two hypotheses and to assign functions to accessory genes, we employed reverse genetics. The only genes that had been previously inactivated were *pppA* and *pppD* (10). However, mutants had been generated by replacement with a selectable marker. To avoid polar effects, we generated markerless, in-frame deletion mutants of *pppA* and *pppD*, in addition to each of nine predicted accessory genes, i.e., *pppE* to *pppM* (**Fig. 1**). Prediction of the borders of the BGC was based on MultiGeneBlast results from our previous investigation (10), that showed *pppA*-*pppD* and *pppG*-*pppK* to be conserved in *Pseudovibrio* and *Pseudomonas*, *pppEF* to be conserved in *Pseudovibrio* and *pppL*-*pppP* to be present in some *Pseudovibrio* strains.

All *P. brasiliensis* mutants were generated using homologous recombination and confirmed by PCR (**Figs. S1-S2**). Different congeners of PA, PB and PC are produced by *P. brasiliensis* (**Fig. 1**). Here we will refer only to the major congeners PA1, PB1/2/3 (**Fig. S3**) and PC1/2/3 (**Figs. S4-S5**) as identified by Matrix-Assisted Laser Desorption/Ionization Time of Flight (MALDI-ToF) mass spectrometry (MS) and liquid chromatography MS (LC-MS), respectively, since minor congeners are not always detected.

Deletion of flanking genes *pppL* and *pppM* showed no effect on the pseudovibriamides produced by *P. brasiliensis* (**Figs. S6-S8**). Genes *pppLMNOP* appear to be part of the same operon (**Fig. 1**), and they are not conserved in all *Pseudovibrio* strains (10) which agrees with a non-requirement for pseudovibriamide biosynthesis. They were thus assigned as not part of the *ppp* BGC and we did not generate deletion mutants of *pppN*, *pppO*, and *pppP*. In contrast, we observed changes in pseudovibriamide production for most other *P. brasiliensis* mutants (**Fig. 2**) as described below.

**Figure 2.**
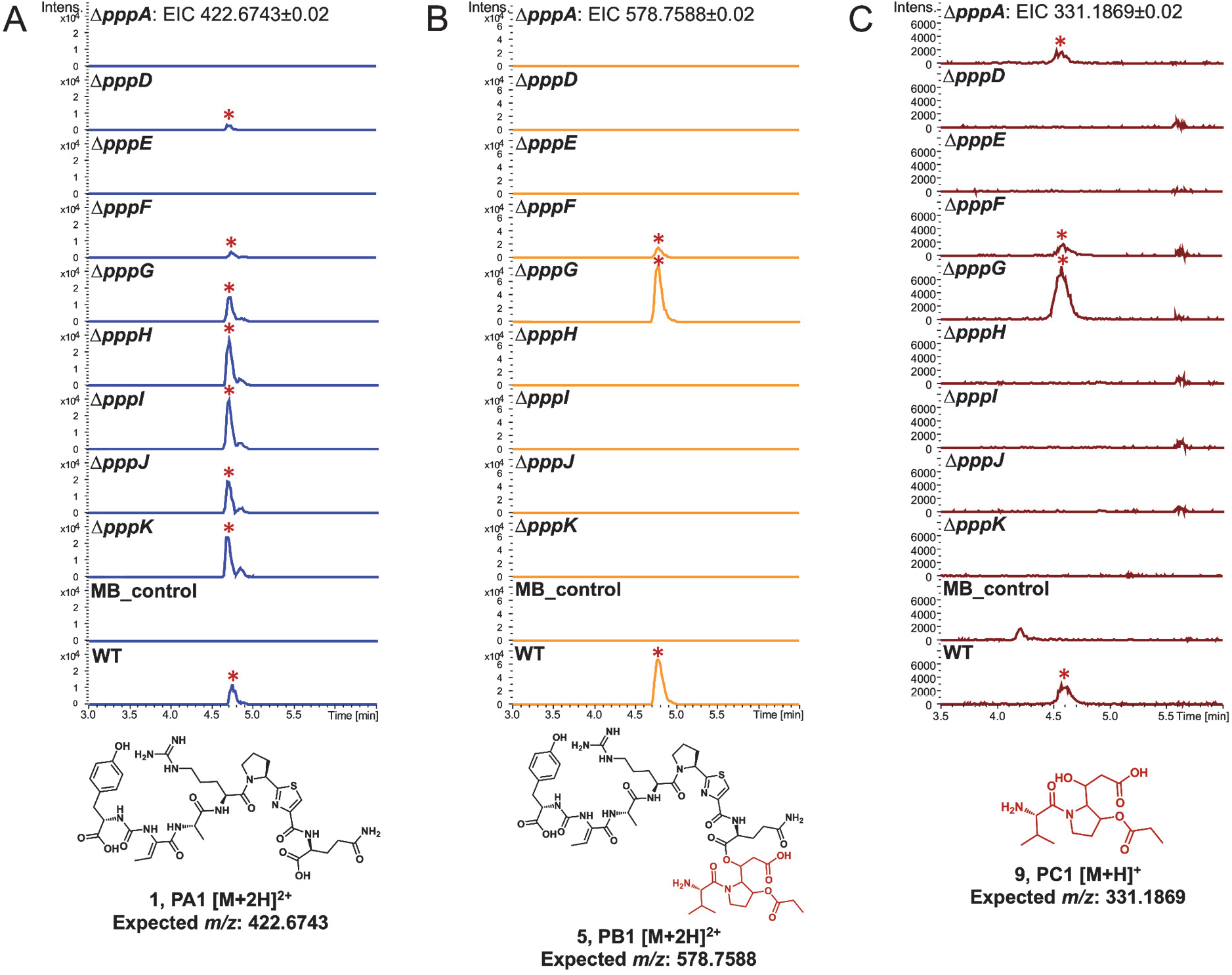
Comparison of the production of PA, PB, and PC representatives between the wild type and mutants. Extracted Ion Chromatograms (EIC) from LC-MS analyses of wild type (WT) and mutants. (**A**) PA1. (**B**) PB1. (**C**) PC1. Pseudovibriamides-related EIC peaks are marked with red asterisks. MB control is the extract of marine broth as negative control. The same mass filter (the expected *m/z* ± 0.02) was applied to all extracts. The same intensity scale was applied in-between strains for each pseudovibriamide. All analyses were performed in at least triplicates.

The Δ*pppA* mutant did not produce PA or PB, matching the result of our previous study using *pppA::neo* mutants (**Figs. 2AB, S9-S11**) (10). However, we observed the production of a metabolite we tentatively identify here as PC by MS analyses, consistent with PppD being still intact in the Δ*pppA* mutant (**Figs. 2C, S4-S5**). In contrast, the Δ*pppD* mutant produced only PA in agreement with previous results with *pppD::neo* (**Figs. 2, S9-S11**) (10).

The Δ*pppE* mutant produced no pseudovibriamides consistent with PppE’s crucial role as a 4’-phosphopantetheinyl transferase to activate NRPS and NRPS-PKS enzymes (**Figs. 2, S4-S5, S9-S11**) (18). Genetic complementation using pYDcompE (**Table S2**) successfully recovered the production of pseudovibriamides (**Fig. S12**). The Δ*pppF* mutant showed a reduction in overall pseudovibriamide abundance, whereas genetic complementation using pYDcompF improved pseudovibriamide production (**Fig. S13**), consistent with a role for PppF as a type II, proofreading thioesterase that regenerates mis-acylated ACPs (19, 20).

Mutants Δ*pppH, ΔpppI,* and Δ*pppJ* produced only PA (**Fig. 2**). Blast analyses revealed sequence similarity to unknown proteins, except for PppH which showed sequence similarity to SGNH hydrolases (**Table S1**). Additionally, protein structure prediction using Phyre2 (21) suggested PppH, PppI and PppJ belong to hydrolase or acyltransferase families of proteins whereas analysis using CLEAN (22) predicted all three to be transferases. To test for polar effects, we performed genetic complementation using plasmids pDS00H, pDS00I and pDS00J, respectively (**Table S2**). The production of all pseudovibriamides was recovered in each of the *P. brasiliensis* mutants, ruling out polar effects (**Figs. S14-S18**). We hypothesize that PppHIJ are involved in propionylation of the hydroxyproline residue and/or function as a *trans*-acyltransferase to load the PKS module of PppD. If PppHIJ are involved in propionylation, we expected to observe a pseudovibriamide analog lacking this modification, however, none was detected. The data suggests that PppHIJ may act together as a complex and that they are each crucial for PB and PC biosynthesis. Another hypothesis we investigated was that PppM, predicted to be a GCN5-related *N*-acetyltransferase, could catalyze propionyl transfer or function as the acyltransferase that is missing in module 10. However, as stated above, Δ*pppM* mutants had the same metabolite profile as the wild type (**Figs. S7-S8**), indicating that *pppM* is not required for pseudovibriamide biosynthesis.

Thus, no PB hydrolase was identified. Instead, it is conceivable that the TE in module 7 can accept either water as the nucleophile leading to PA or products of the PppD enzyme (PC or pre-PC) leading to PB.

### PppK is a hydroxylase

PppK shows sequence similarity to Fe(II)/α-ketoglutarate dependent dioxygenases (23) and we predicted it would be responsible for the hydroxyl group in the proline residue of PB and PC. Accordingly, the Δ*pppK* mutant produced PA as in the wild type (**Fig. 2**) but PB and PC analogs that showed a Δ72 Da mass loss, indicating that they lacked the hydroxyl group on the proline and consequently also lacked propionylation (**Figs. 3, S19-S20**). Fragmentation patterns from MS^2^ spectra further verified the assignment (**Figs. S21-S22**). Genetic complementation using plasmid pVL00K recovered the production of PB and PC (**Fig. S23**). The timing of hydroxylation remains unknown. Three scenarios are conceivable. PppK could be either a proline hydroxylase acting on free proline, or it could act on-line while the proline-containing substrate is attached to PppD, or after pre-pseudovibriamides are released from PppD, either on pre-PC or pre-PB.

**Figure 3.**
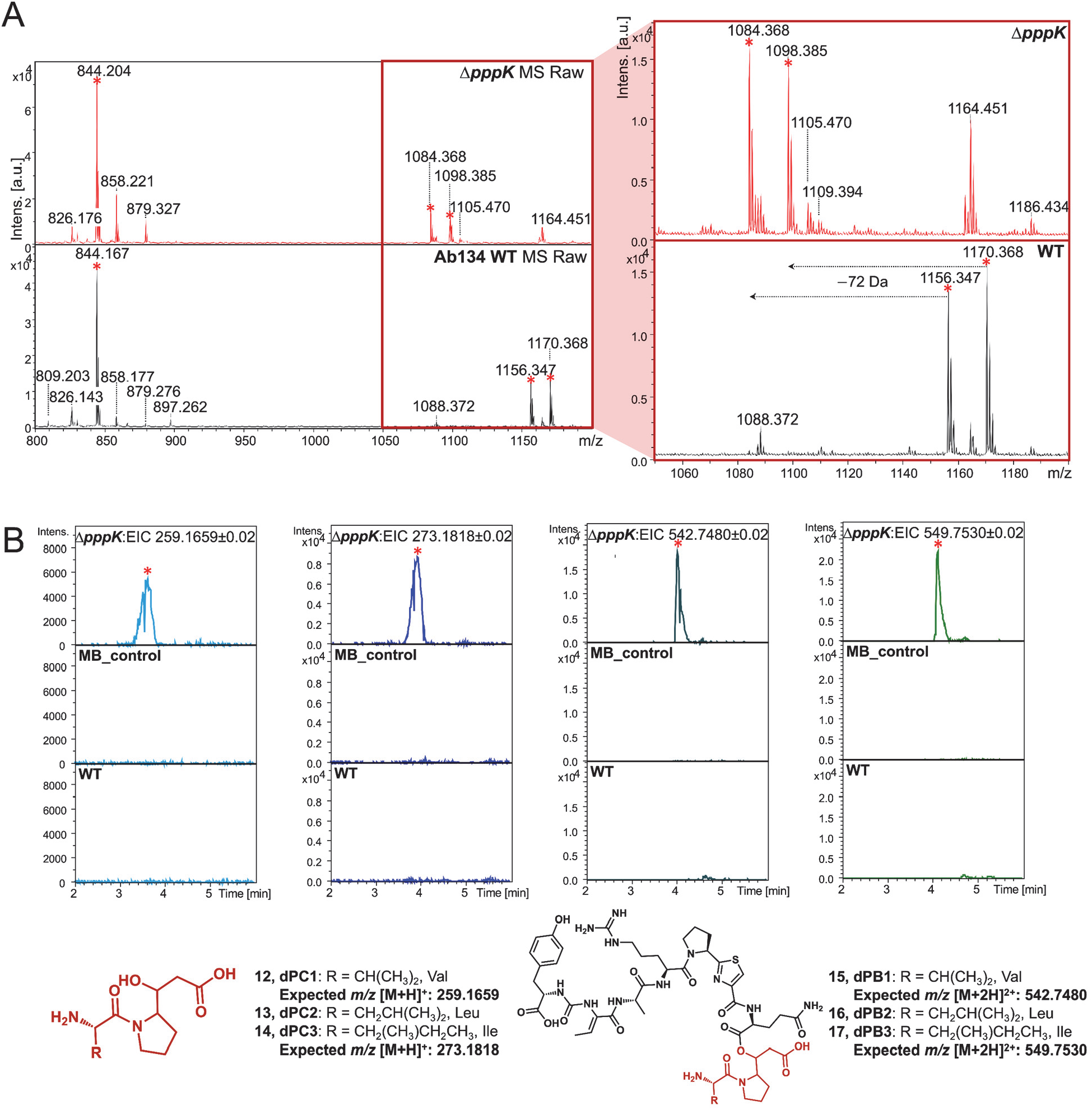
Comparison of PB and PC production between the wild type and the Δ*pppK* mutant. (**A**) MALDI-ToF MS analyses. Molecular features for pseudovibriamides are indicated with red asterisks. The peak at *m/z* 844.2 represents PA1 ([M+H]^+^); *m/z* 1156.3, PB1 (Val, [M+H]^+^); *m/z* 1170.4, PB2 or PB3 (Ile or Leu, [M+H]^+^); *m/z* 1084.4, depropionylated PB1 (dPB1 Val, [M+H]^+^); and *m/z* 1098.4, depropionylated PB2 or PB3 (dPB2, Leu, [M+H]^+^ or dPB3, Ile, [M+H]^+^). The range from *m/z* 1050 to 1200 was zoomed in (red box) to show the *m/z* change of PBs. (**B**) LC-MS analyses. dPC, depropionylated PC; dPB, depropionylated PB. Extracted Ion Chromatogram (EIC) from left to right: dPCs (Val, [M+H]^+^, *m/z* 259.1659; and Leu or Ile, [M+H]^+^, *m/z* 273.1818), dPBs (Val, [M+2H]^2+^, *m/z* 542.748; and Leu or Ile, [M+2H]^2+^, *m/z* 549.753). The same mass filter (the expected *m/z* ± 0.02) was applied to all samples. The predicted structure is listed below each chromatogram. All analyses were performed in at least triplicates.

### The amount of pseudovibriamides exported is not significantly affected in Δ*pppG* and Δ*pppL* mutants

According to previous imaging mass spectrometry studies, PA and PB are excreted (10). Both *pppG* and *pppL* genes are predicted to encode transporters that could be responsible for pseudovibriamide export (**Table S1**). Both Δ*pppG* and Δ*pppL* mutants produce all pseudovibriamides (**Figs. 2**, **S4-S6**, **S9-S11**). To check for export, we extracted pseudovibriamides from cell pellets and supernatant separately (**Fig. S24**). The export ratio – defined as the amount of pseudovibriamides in the supernatant by the amount in the cell pellet – of PB1, PB2, and PB3 was slightly higher for the wild type and genetically complemented Δ*pppG* strain using pYDcompG compared to Δ*pppG* cultures, but not statistically significant (**Fig. S25A**). Likewise, no statistically significant difference was observed between the wild type and the Δ*pppL* mutants (**Figs. S25B, S26-S38**). Thus, the protein involved in pseudovibriamide export remains to be identified.

### Effect of gene deletion on swarming motility

We performed swarming assays for the 11 in-frame deletion mutants generated here. Both Δ*pppA* and Δ*pppD* mutants showed consistent swarming phenotypes as reported previously (**Figs. 4**, **S39-S40**) (10). Moreover, wild-type level swarming motility was observed for Δ*pppH, ΔpppI,* and Δ*pppJ* mutants, which have the same metabolite profile as the Δ*pppD* mutant, that is, production of PA only (**Fig. S41**). In contrast, the Δ*pppE* mutant that produces no pseudovibriamides showed decreased swarming motility like the Δ*pppA* mutant that produces only PC (**Figs. 4, S42**). The *pppA::neo* mutant had been previously genetically complemented showing restoration of swarming proficiency (10). Likewise, the swarming motility of the Δ*pppE* mutant was successfully recovered by genetic complementation (**Fig. S43**).

**Figure 4.**
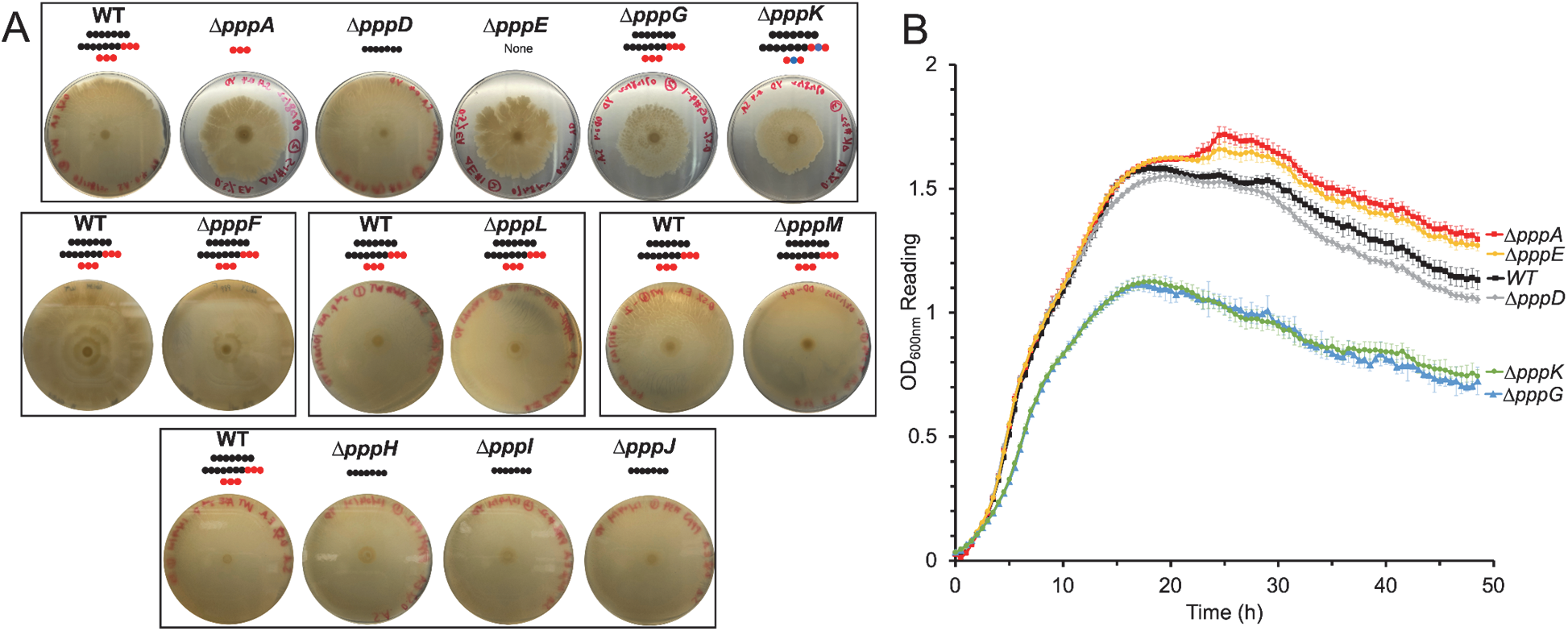
Effect of *ppp* gene inactivation on flagellar motility. (**A**) Swarming assays performed on marine broth with 0.5 % Eiken agar. Pictures shown were taken 72 hours after inoculation. The assay was performed multiple times, each time in at least triplicates with the wild type (WT) as the control, and similar results were obtained each time (see **Figs. S39-S47**). Representative results are shown. Plates are grouped and boxed based on assays that were run together the same day. Pseudovibriamides are represented by beads: PA, seven black beads; PB, seven black beads and three red beads; PC, three red beads; depropionylated PB, seven black beads, two red beads, and one blue bead; depropionylated PC, two red beads and one blue bead. (**B**) Growth of strains from top box in panel A as measured by OD600. *N*=6. Error bars indicate standard deviation. Note that the apparent reduced swarming of Δ*pppG* and Δ*pppK* mutants is in fact due to reduced growth.

The combined results of Δ*pppD, ΔpppH, ΔpppI,* and Δ*pppJ* mutants suggest that only PA is required for wild-type level swarming motility. However, both Δ*pppK* and Δ*pppG* mutants showed impaired swarming, even though PA was produced in these mutants (**Fig. 4A**). Moreover, pseudovibriamide production was restored by genetic complementation using pVL00K and pYDcompG, respectively (**Fig. S23**); however, swarming was not, suggesting factors other than pseudovibriamides were involved in the phenotype of Δ*pppK* and Δ*pppG* mutants (**Fig. S44**). It turns out both Δ*pppG* and Δ*pppK* mutants showed decreased growth, suggesting the cause of apparently reduced swarming motility is in fact related to reduced growth (**Fig. 4B**). Whole genome sequencing of both Δ*pppG* and Δ*pppK* mutants showed several potential mutations (**Table S3-S4**). It remains to be shown which mutations are responsible for the reduced growth phenotype.

Other mutants (Δ*pppF*, Δ*pppH*, Δ*pppI*, Δ*pppJ*, Δ*pppL*, and Δ*pppM*) showed no observable defect in swarming motility (**Figs. 4A, S41, S45-S47**).

### Transmission electron microscopy identifies no apparent changes in flagella

We used transmission electron microscopy (TEM) to visualize cells of strains with different compositions of pseudovibriamides, that is, Δ*pppA*, Δ*pppD*, Δ*pppE* mutants and the wild type. No apparent differences in flagella were observed between the strains, although there seemed to be more cell aggregation for Δ*pppA* and Δ*pppE* mutants which is consistent with decreased swarming motility (**Fig. S48**).

### PA and PB modulate gene transcription whereas PC does not

To test the hypothesis that pseudovibriamides affect motility by modulating gene transcription, RNA-seq data sets were obtained for the wild type and Δ*pppA*, Δ*pppD*, and Δ*pppE* mutants having different compositions of pseudovibriamides (**Fig. 5**). Triplicate samples of each strain are similar in terms of gene expression pattern, which is more apparent in the principal component analysis (PCA) plot where triplicate samples from each strain cluster together, supporting the quality of the data (**Fig. 5B**). The highly similar transcriptomic profiles of Δ*pppA* and Δ*pppE* mutants indicates that PC has none or only minimal effect on gene transcription (**Fig. 5B**). In contrast, the large differences in principal components 1 and 2 observed between the wild-type and either the Δ*pppD* mutant or the Δ*pppA/ΔpppE* mutants indicates that PA and PB have major effects on gene transcription (**Fig. 5B**).

**Figure 5.**
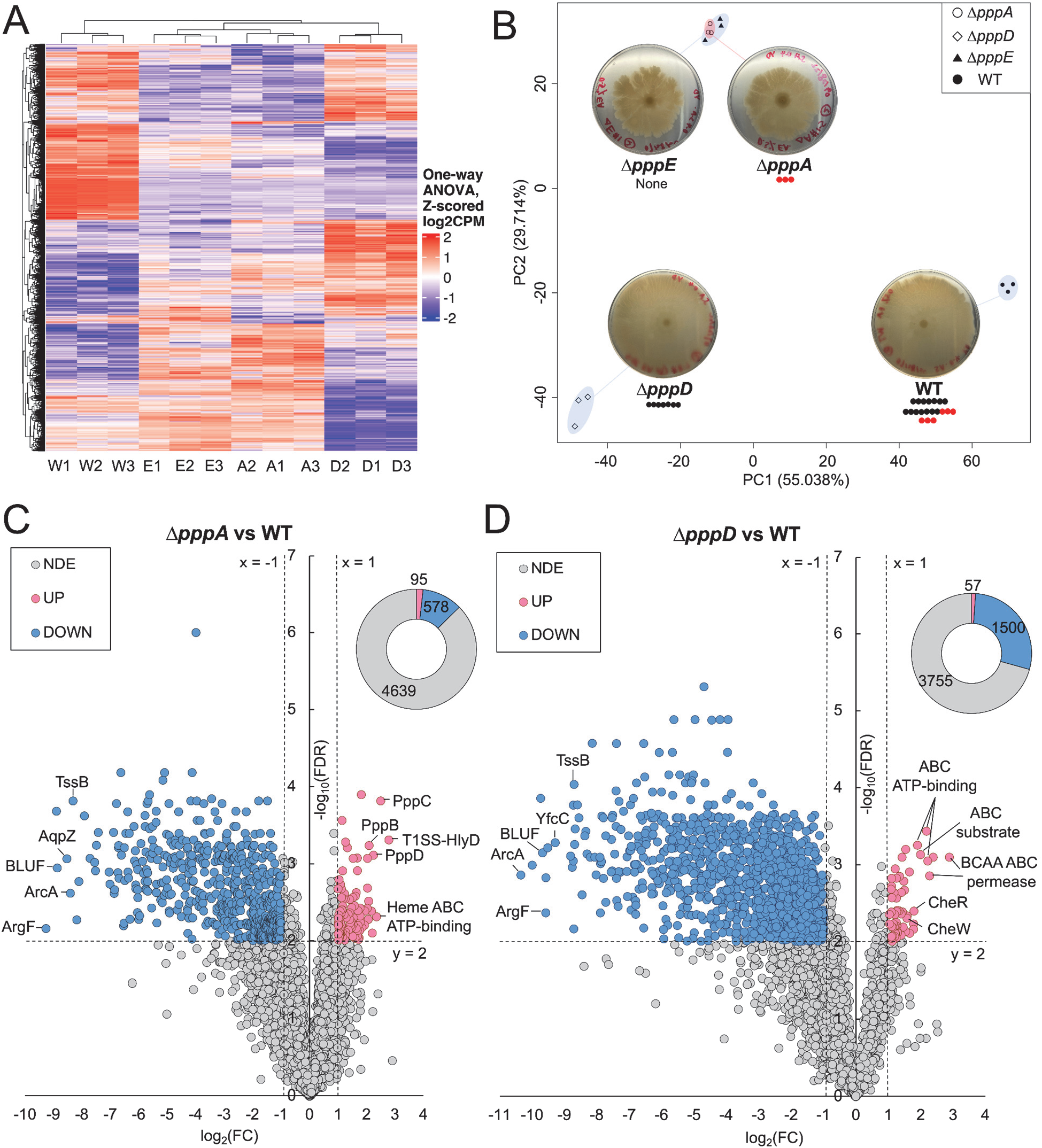
Overview of RNA-Seq results. (**A**) Heatmap of gene expression based on Z-scored counts per million (CPM) of the wild type (W1-3), Δ*pppA* (A1-3), Δ*pppD* (D1-3), and Δ*pppE* mutant (E1-3). Each column represents one of three replicates. Each row represents one of 5,312 genes. One-way analysis of variance (ANOVA) was used to compare whether four samples’ means are significantly different or not. Values in each row were scaled to CPM mean of the row by using Z-score normalization. (**B**) Principal component analysis (PCA) of gene expression based on normalized CPM of triplicate samples. Corresponding swarming images at 72 h are shown and pseudovibriamide composition is indicated for each strain. Seven black beads represent PA; Seven black beads plus three red beads represent PB; three red beads represent PC. (**C**) and (**D**) Volcano plots of differentially expressed (DE) genes identified between the wild type and Δ*pppA* and Δ*pppD* mutants, respectively, using transcripts per million (TPM). FDR, false discovery rate or *q*-value; FC, fold-change; NDE, non-differential expressed; UP, upregulated; DOWN, downregulated; Blue dots or blue donut portion, downregulated genes; Pink dots or pink donut portion, upregulated genes; and grey dots or grey donut portion, non-differentially expressed genes. The same threshold (dotted lines) was applied to all differential expression analyses, that is, FC ≤ −2 (x = −1) or FC ≥ 2 (x = 1), and FDR ≤ 0.01 (y = 2). The total number of UP, DOWN, and NDE genes are listed in the donut chart. Labeled genes are TssB, type VI secretion system contractile sheath small subunit; YfcC, Arginine/ornithine antiporter; AqpZ, aquaporin Z; ArcA, arginine deiminase; ArgF, ornithine carbamoyltransferase; BLUF, blue light using flavin domain; T1SS-HlyD, HlyD family type I secretion periplasmic adaptor subunit; BCAA ABC permease, branched-chain amino acid ATP-binding cassette transporter permease; ABC ATP-binding, ATP-binding cassette transporter ATP-binding protein; ABC substrate, ATP-binding cassette transporter substrate binding protein; Heme ABC ATP-binding, heme ATP-binding cassette transporter ATP-binding protein; CheW, chemotaxis protein; and CheR, protein-glutamate O-methyltransferase. Some outstanding dots left unlabeled are hypothetical proteins.

### Pairwise differential expression analyses methods and parameters

To ensure the identified differentially expressed (DE) genes would have physiological relevance, we started by testing distinct analysis methods and performing follow up experiments for validation. We first used DESeq2, which is based on negative binomial distribution (**Fig. S49**). An upregulation of *ppp* core biosynthetic genes was observed in both Δ*pppA* and Δ*pppD* mutants in which *pppBCD* and *pppABC* appeared upregulated compared to the wild type, respectively (**Fig. S49**). According to the RNA-seq data, a promoter is located upstream of the *pppA* gene (P*_pppA_*) (**Fig. S50**). Upregulation of *ppp* genes in the mutants is suggestive of negative autoregulation by pseudovibriamides as reported for other products (24). To test this hypothesis, we cloned P*_pppA_* directly upstream of *GFP* in the promoterless pSEVA227M-based vector (**Table S2**). In the case of negative autoregulation, we expected to observe increased GFP production in the mutants. However, this was not the case (**Fig. S51**). Instead, the promoter probe studies suggested that DESeq2 resulted in the identification of false positives.

We next tested transcripts per million (TPM) for data normalization and DE calls (**Figs. 5C-5D**). Genes *pppBCD* are still upregulated but only in the Δ*pppA* mutant, which is likely an artifact of bringing P*_pppA_* closer to the other genes in the operon by deleting ∼8,000 bp of the *pppA* gene. Thus, the results indicate that TPM is more accurate for comparing gene expression levels across our samples. Therefore, TPM values were chosen for further analyses. The total number of DE genes was reduced from 1,298 to 673 in the Δ*pppA* mutant, and from 1,616 to 1,557 in the Δ*pppD* mutant when using TPM (**Figs. 5C-5D, S49**). There was only one gene, *pppC*, upregulated, and no downregulated gene in the Δ*pppA* mutant when compared to Δ*pppE* mutant using FDR ≤ 0.01 as the cutoff. Because the PCA plots overlap, we considered Δ*pppA* and Δ*pppE* mutants to be indistinguishable, and the FDR cutoff of ≤ 0.01 as appropriate for further analyses.

### Global effects of PA and PB

The large number of DE genes identified (**Figs. 5C-5D**) suggests that PA and PB have a global effect on transcription that goes beyond motility. To obtain a broader view of the role of PB, DE genes of Δ*pppD* mutant (missing PB but having PA) when compared to the wild type were classified based on clusters of orthologous genes (COG) (**Fig. 6A, Supplemental file 1**). Except for the poorly characterized (PC) group, the metabolism (M) group is the largest, followed by the cell processes and signaling (CPS) group (**Fig. 6A**). Regarding specific categories within these groups, cell wall biogenesis [M], inorganic ion transport and metabolism [P], amino acid transport and metabolism [E], and transcription [K] dominate (**Fig. 6A**, **Supplemental file 1**). Strikingly, downregulated genes outnumber upregulated genes by roughly 30 to 1, indicating that PB may have a primary effect on gene activation.

**Figure 6.**
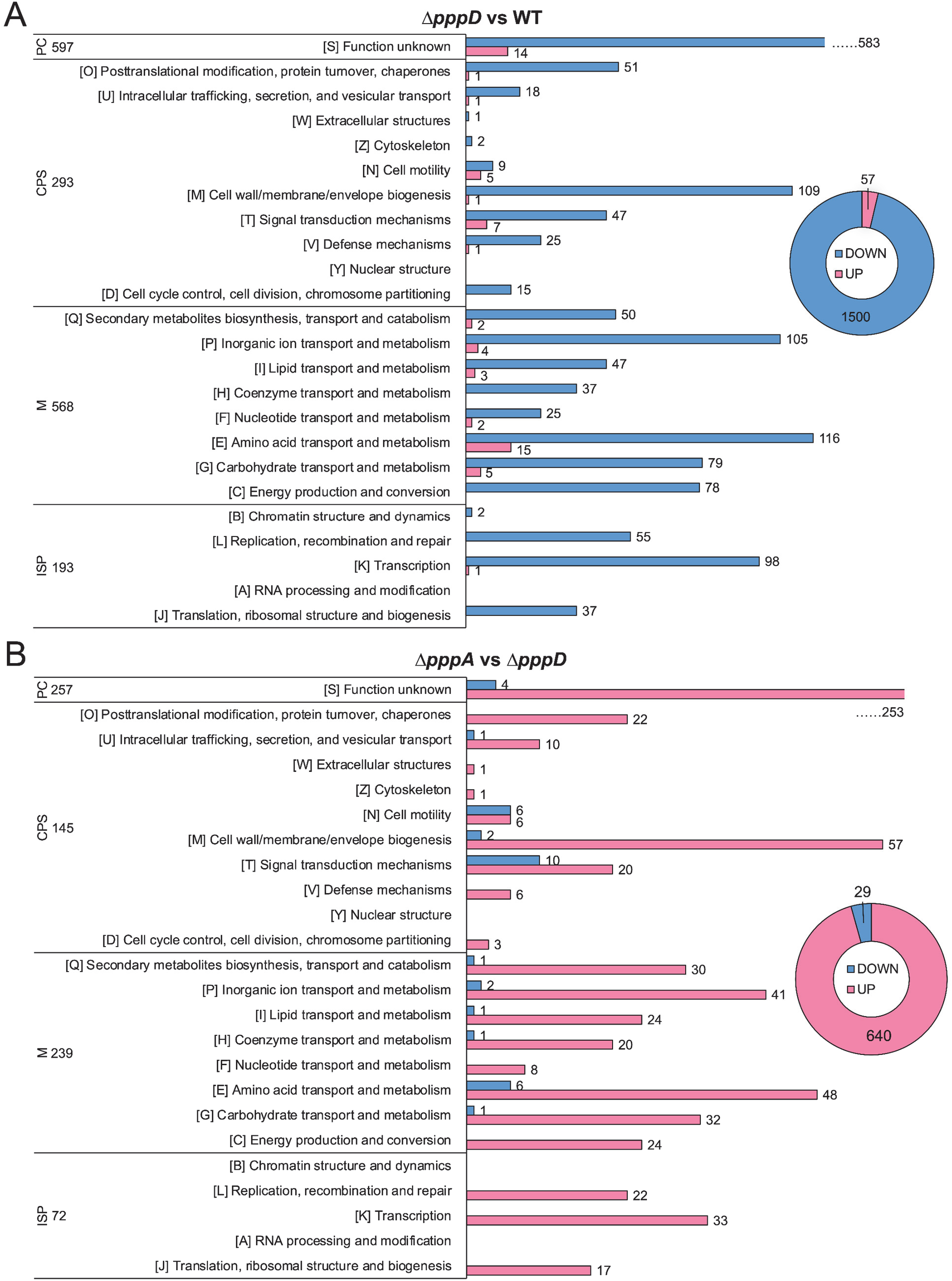
COG classification of DE genes potentially affected by PA and PB. (**A**) DE genes of Δ*pppD* mutant compared to the wild type. (**B**) DE genes of Δ*pppA* mutant compared to the Δ*pppD mutant*. Blue, downregulated; Pink, upregulated. PC, poorly characterized; CPS, cellular processes and signaling; M, metabolism; and ISP, information storage and processing. The total number of genes in each category is listed.

Regarding genes potentially affected by PA, the Δ*pppA* mutant (missing PA) was compared to the Δ*pppD* mutant (produces only PA) (**Fig. 6B, Supplemental file 1**). Contrary to what was observed in the previous comparison, upregulated genes outnumber downregulated genes by roughly 20 to 1 in the Δ*pppA* mutant, indicating that PA may have an inhibitory effect on gene transcription. Metabolism is still the largest group followed by cell processes and signaling. The categories most affected are also the same (**Fig. 6B**). Thus, the results suggest that PB and PA may have opposite roles in modulating gene transcription. Indeed, 446 DE genes overlap between the two pairwise analyses and are inversely regulated (**Supplemental file 1**).

### Identifying DE genes potentially involved with reduced swarming motility

We considered two assumptions for narrowing down potential candidate genes related to swarming motility. On one hand, the Δ*pppD* mutant may possess the same set of genes unaffected compared to the wild type, which are DE in Δ*pppA* and Δ*pppE* mutants, resulting in reduced swarming motility. On the other hand, the Δ*pppD* mutant might harness different pathways than the wild type for promoting swarming motility, resulting in the same observable phenotype.

Assuming the first scenario, genes were compiled that were DE in both Δ*pppA* and Δ*pppE* mutants but also non-differentially expressed (NDE) in the Δ*pppD* mutant, each compared to the wild type (**Fig. 7A**). As a result, 12 upregulated genes and 5 downregulated genes in Δ*pppA/E* mutants were identified (**Figs. 7A**, **S52A; Table S5**). According to COG, most upregulated genes, except those encoding hypothetical proteins, are ATP-binding cassette (ABC) transporters belonging to the metabolism category. This pattern generally matches observations from a study in *Pseudomonas aeruginosa* where the authors found that genes related to the transport of small molecules were upregulated in non-swarming cells (25).

**Figure 7.**
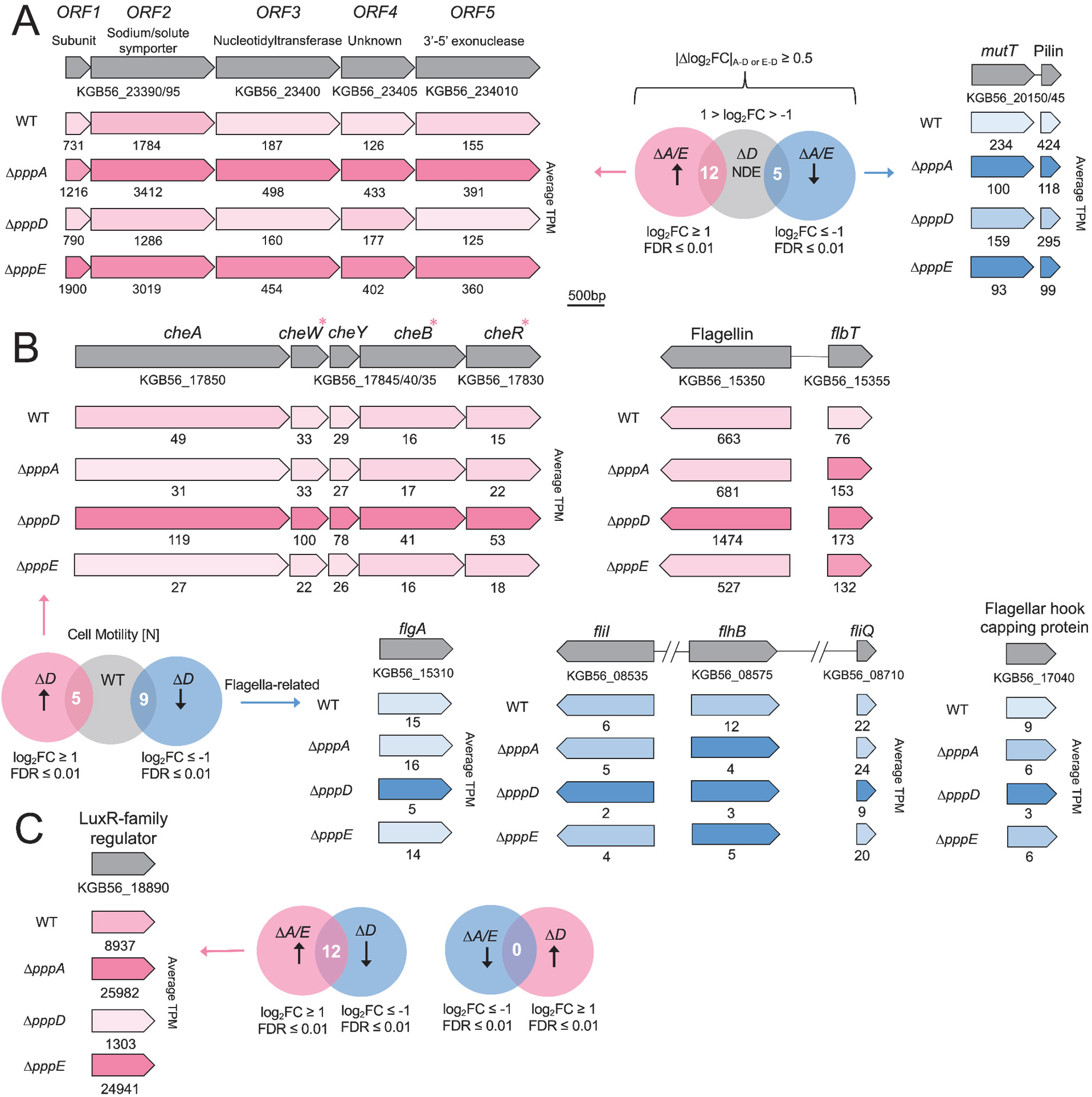
DE genes potentially involved in differential swarming motility. (**A**) Assumption 1: the Δ*pppD* mutant may possess the same set of genes unaffected compared to the wild type, which are DE in Δ*pppA*/Δp*ppE* mutants compared to the wild type. Filters used for selecting such genes. |Δlog2FC|, the absolute difference of log2FC between Δ*pppA,* or Δ*pppE* and Δ*pppD* mutants was set to be larger than or equal to 0.5 to exclude genes with minor variation in differential expression. See **Fig. S52A** for COG categories. Examples of DE operons are shown, that is, a five-gene operon that is upregulated in the Δ*pppA/E* mutants (subunit, the small subunit of the sodium/solute symporter), and a two-gene operon that is downregulated in the Δ*pppA/E* mutants including MutT CDS (KGB56_20150) and Flp family type IVb pilin CDS (KGB56_20145). Pink, upregulated; blue, downregulated. The darker the pink, the higher the relative expression level. The darker the blue, the lower the relative expression level. TPM values are indicated below genes. (**B**) Assumption 2: the Δ*pppD* harnesses different pathways than the wild type for promoting swarming motility, resulting in the same observable phenotype. Selected motility genes that are up or downregulated in the Δ*pppD* mutant are shown. Upregulated chemotaxis genes (FDR ≤ 0.01) are denoted with pink asterisks. *cheA*, chemotaxis histidine protein kinase; *cheW*, linker protein; *cheY*, chemotaxis response regulator; *cheB*, chemotaxis response regulator protein-glutamate methylesterase; *cheR*, chemotaxis glutamate *O*-methyltransferase; *flbT*, flagellar biosynthesis repressor; *flgA*, flagellar basal body P-ring formation protein; *fliI*, flagellar biosynthesis type III secretory pathway ATPase; *flhB* and *fliQ,* flagellar biosynthesis protein. (**C**) Filters used for selecting DE genes in Δ*pppA*/Δp*ppE* mutants that are reversely regulated in the Δ*pppD* mutant compared to the wild type. See **Fig. S53** for COG categories. The expression levels (in TPM) of a LuxR family transcriptional regulator CDS (KGB56_18890) are indicated for mutants and the wild type. The darker the pink, the higher the expression level.

There are no genes in the cell motility [N] category. We next investigated signal transduction [T], which could include genes regulating the swarming motility phenotype directly or indirectly. Of the two genes upregulated in the T category (**Table S5, Supplementary Results**), one is predicted to encode a nucleotidyltransferase and is located within a five-gene operon (**Fig. 7A**). The other four genes were not hits due to the stringent cut off we set, but they still show considerable log_2_FC of 0.73 - 1.78. ORF2 encodes a sodium/solute symporter, and ORF1 encodes its small subunit. These genes are highly expressed in the Δ*pppA* mutant, e.g., the TPM value of 3411 for ORF2 ranks as the 25^th^ most highly transcribed gene. An overexpression of this symporter (26) will not only increase the uptake of nutrients, but also of sodium. Sodium influx is used to power flagellar rotation (27, 28). It is possible that an inbalance in the sodium gradient in Δ*pppA/E* mutants slows down the flagellar motor, leading to reduced but not abolished motility. The remaining one gene upregulated in the signal transduction [T] category is described under **Supplementary Results**.

There are only five downregulated genes in Δ*pppA/E* mutants but NDE in the Δ*pppD* mutant (**Figs. 7A**, **S52A, Table S5**). One of them is predicted to encode a Nudix hyrdolase (29), (30) (**Fig. S52B**). Inactivation of Nudix hydrolase genes *mutT* from *E. coli* and *PA4400* from *P. aeruginosa* results in a higher mutation rate, and *PA4400* can complement an *E. coli mutT* deficient strain (31, 32). A recent study showed that *P. aeruginosa* Δ*PA4400* possessed severely impaired swarming motility (33). Thus, the downregulation of this gene could help explain the reduced swarming motility of Δ*pppA/E* mutants. In addition, the gene downstream of the *mutT* homolog is predicted to encode a type IV pilin subunit (34). A previous study showed that a *P. aeruginosa* type IV pilin mutant was unable to swarm (35). Thus, downregulation of the type IV pilin gene in Δ*pppA/E* mutants may contribute to the attenuated swarming motility as well.

To probe the second scenario that the Δ*pppD* mutant might harness different pathways than the wild type for promoting swarming motility, the direct pairwise differential expression analysis between Δ*pppD* mutant and the wild type was analyzed (**Fig. 6A**). In the most obvious category, cell motility [N], there are five genes upregulated in the Δ*pppD* mutant (encoding one flagellin protein, one flagellar biosynthesis repressor FlbT, and three chemotaxis proteins *cheW*, *cheB* and *cheR* that are part of a five-gene operon), and nine genes downregulated (encoding five flagella-related proteins; three L,D-transpeptidases, and one type IV secretion system protein) (**Figs. 6A, 7B, Supplemental file 1**). The downregulation of genes encoding components of flagella and the upregulation of a flagella biosynthesis repressor gene suggests the Δ*pppD* mutant should have reduced swarming motility unless there was a compensatory mechanism, which seems to be the case with the upregulation of chemotaxis genes. Downregulation of chemotaxis genes was observed in a swarming-deficient mutant of *Vibrio parahaemolyticus* (36); conversely, upregulation might promote swarming motility in the Δ*pppD* mutant as a compensatory mechanism.

To identify genes that may be implicated in the opposing effects of PA and PB, we searched for genes that were DE in opposite directions in the Δ*pppD* and Δ*pppA/E* mutants when compared to the wild type, respectively, and identified 12 genes (**Figs. 7C, S53, Table S6**). One gene caught our attention because it is predicted to encode a LuxR-type transcriptional regulator (**Fig. 7C, Supplemental file 1**). The original LuxR is a well known cell-density-dependent transcriptional regulator involved in quorum sensing in *Vibrio fischeri* (37). Many LuxR variants exist, including QscR and VjbR (38, 39) which show sequence indentity to the DE LuxR we identified albeit low (26%). Analogously to what we observed with Δ*pppA/E* mutants, upregulation of QscR from *P. aeruginosa* PAO1 resulted in reduced swarming motility (38). Conversely, downregulation in the Δ*pppD* mutant may improve motility potentially as a compensatory mechanism. Another gene in the signal transduction [T] category is described under **Supplementary Results**.

Taken together, compensatory mechanisms seem to be at play, resulting in the same observable swarming phenotype between the wild type and the Δ*pppD* mutant. Moreover, LuxR family regulators can influence many genes, such as 14% of *Brucella melitensis* genes by VjbR (39). Thus, the identified LuxR family regulator is a candidate gene that could explain the large differences in transcription between mutants and the wild type (**Fig. 5**), but future studies are necessary to test this hypothesis.

## Discussion

The term ‘holobiont’ has been coined to express the crucial relationship between plants and animals and their associated microbes (4). Microbial metabolites are important in establishing and maintaining microbe-host associations. For instance, motility, mediated by bacterial metabolites, is known to be important for host colonization (9). We previously identified a link between swarming motility and pseudovibriamides (10). The *ppp* gene cluster that encodes pseudovibriamides (**Fig. 1**) is found not only in bacteria that interact with marine sponges but also in terrestrial bacteria that interact with plants and animals (10).

The main goal of the present work was to reveal how pseudovibriamides affect swarming motility. We started by obtaining mutant strains with different compositions of pseudovibriamides while gaining insight into pseudovibriamide biosynthesis. We considered two hypotheses to explain the presence of PA, PB and PC in *P. brasiliensis*. Either PA and PC represent hydrolysis products of PB catalyzed by an accessory hydrolase, or PA is directly released from PppC catalyzed by the thioesterase domain using water as the nucleophile (**Fig. 1**). We considered PppH as a PB hydrolase because it shows sequence similarity to the hydrolase family of enzymes. If true, we would expect the hydrolase-inactive mutant to produce only PB. However, the Δ*pppH* mutant produced only PA (**Fig. 2**). None of the accessory genes was identified as encoding a PB hydrolase. Instead, the second hypothesis seems plausible that the TE domain in PppC can accept either water as nucleophile resulting in PA or it can accept PC (or pre-PC) resulting in PB. *In vitro* studies are necessary to further test this hypothesis.

Swarming motility assays suggested that only PA is required for wild-type level swarming motility. Mutants that produced only PA (Δ*pppD, ΔpppH, ΔpppI,* and Δ*pppJ*) displayed swarming motility comparable to the wild type (**Figs. 4, S41**), whereas mutants that produced only PC (Δ*pppA*) or no pseudovibriamides (Δ*pppE*) showed reduced swarming motility (**Fig. 4**).

We next performed transcriptomic studies of the wild-type and of mutant strains that produce either only PA (Δ*pppD*), only PC (Δ*pppA*) or no pseudovibriamides (Δ*pppE*) to test whether pseudovibriamides influence gene transcription. Since several RNA-seq normalization methods are available and errors in normalization can result in false positives (41), we first compared normalization methods and performed follow up experiments for validation using a GFP reporter assay to ensure the identified DE genes would have physiological relevance. From the two methods tested – DESeq2 and TPM, representing normalization by distribution and library size, respectively (41) – TPM seemed to eliminate false positives (**Figs. 5CD, S49**). A recent study to evaluate RNA-seq normalization methods also concluded that TPM performed best in preserving biological signal (42).

From the transcriptomic data, we concluded that PC plays no role in modulating transcription since the transcriptomes of Δ*pppA* and Δ*pppE* mutants were equivalent (**Fig. 5B**). In contrast, PA and PB play major roles as 13% of the total number of genes are differentially expressed when PA and PB are missing (Δ*pppA* vs. wild type, **Fig. 5C**) and 29% of the total number of genes are differentially expressed when PB is missing but PA is present (Δ*pppD* vs. wild type, **Fig. 5D**). The results also suggested that PA and PB have opposite effects on a subset of 446 DE genes, with PB having a primary role in gene upregulation and PA in downregulation (**Fig. 6, Supplemental file 1**).

Accordingly, a compensatory mechanism appears to be at play in the Δ*pppD* mutant that results in the same observable swarming phenotype as the wild type. This is plausible based on the upregulation of chemotaxis genes while flagella component genes are downregulated. In addition, 12 genes are inversely regulated between the Δ*pppD* mutant and Δ*pppA*/Δ*pppE* mutants when compared to the wild type. For instance, the drastic difference in the transcription level of a LuxR-type regulator may help explain not only the swarming motility phenotypes but also the large differences in transcription between mutants and the wild type (**Fig. 7C**).

In conclusion, motility enables bacteria to reach new habitats and to colonize host tissue. Many questions remain to be answered regarding bacterial motility, including identifying which chemical signals mediate motility and how they do so. Here we showed that pseudovibriamides affect motility by modulating transcription, ultimately revealing new signaling molecules. Importantly, the effects of pseudovibriamides appear to extend beyond motility to affect yet-to-be identified phenotypes. Future studies should focus on more detailed biosynthetic investigations such as the timing of hydroxylation by PppK, and the joint role of PppHIJ in propionylation – literature precedence for multi-protein complexes catalyzing acylation does exist (40). Future studies should also elucidate the exact mechanisms by which pseudovibriamides modulate gene transcription.

## Materials and Methods

### General cultivation conditions

*P. brasiliensis* Ab134 was cultivated at 30°C on BD Difco™ Marine Agar 2216 (MA) or in BD Difco™ Marine Broth 2216 (MB) for 18-20 hours unless otherwise noted. Chloramphenicol (8 µg/mL) and kanamycin (200 µg/mL) were used for mutant selection as appropriate. *E. coli* strains were cultured in BD Difco™ Luria Broth (LB) or on LB agar for 18-20 hours. Chloramphenicol (25 µg/mL) and kanamycin (50 µg/mL) were used for mutant selection as appropriate. *E. coli* DH5α(λpir) was used for propagation of pDS132-based vectors and *E. coli* SM10(λpir) for conjugation with Ab134. *E. coli* DH5α was used for propagation of pSEVA227M-based or pAM4891-based vectors and *E. coli* S17-1 for conjugation. All strains were cryo-preserved in 20% glycerol [*v/v*] at −80 °C.

### Plasmid construction

Plasmids used in this study are summarized in **Table S2**. Oligonucleotide primers (**Table S7**) were synthesized by Sigma-Aldrich. Vector pDS132 was used to construct plasmids for in-frame deletion, pSEVA227M for promoter probe studies, and pAM4891 for genetic complementation (43–45). See Supplementary Information for details.

### In-frame deletion and complementation

Mutants were generated by in-frame deletion via homologous recombination (**Fig. S1**). pDS132 or pYD004-based suicide vectors were first transformed into *E. coli* SM10(λpir), which was used as the conjugation donor for transferring vectors into Ab134. The detailed conjugation protocol can be found under Supplementary Information. Obtained clones were analyzed by two parallel PCRs to identify single crossover (SCO) colonies (**Fig. S1**). Confirmed SCO clones were streaked onto non-selective MA plates and incubated overnight at 30°C. MA containing 5% sucrose was used for counterselection of the vector. Chloramphenicol-sensitive clones were analyzed by PCR to confirm the gene replacement (**Fig. S2**). All pAM4891-based complementation vectors were first electroporated into *E. coli* S17-1, which was used as the conjugation donor for introducing vectors into Ab134 mutants. Kanamycin (200 μg/mL) was used to select for the incoming plasmid and exconjugants were confirmed by plasmid extraction and restriction digest.

### Swarming assays

The protocol used was as we previously reported (10) with only minor modifications as detailed under Supplementary Information.

### Pseudovibriamide extraction and analysis

Pseudovibriamides were extracted in two ways based on culture conditions. A) Swarming agar cultures were extracted with one volume (20 mL) of methanol by sonicating for 1 h, after which the extract was filtered through filter paper. B) Liquid cultures were extracted by first capturing metabolites using XAD-7HP resin (Sigma-Aldrich), and then extracting the resin with methanol. Methanol extracts were dried under reduced pressure and stored at −20°C until analysis. See Supplementary Information for further details and for pseudovibriamide extraction from cell pellet and supernatant. Extracts were analyzed by either dried droplet MALDI-ToF MS or by UPLC-QToF-MS/MS as described under Supplementary Information.

### RNA extraction and transcriptomics analyses

The overall scheme is summarized in **Fig. S54**. RNA was isolated from 24-h triplicate cultures using the RiboPure™-Bacteria kit (Thermo Fisher Scientific) following the manufacturer’s instructions. Purified RNA (**Table S8**) with RNA integrity number > 8.0 were sent to SeqCenter (Pittsburgh, MA) for bacterial rRNA depletion RNA sequencing using a NextSeq2000 sequencer giving 2×51bp reads. Demultiplexing, quality control, and adapter trimming was performed with bcl-convert (v3.9.3). Raw Illumina reads of each sample were mapped to the Ab134 whole genome (NCBI accession number GCA_018282095.1) using Geneious. Expression levels (FPKM, RPKM, and TPM) were calculated (all contigs at once) with the option that ambiguously mapped reads were counted as partial matches. Pairwise differential expression analysis was performed by using DESeq2 package in Geneious or using TPM values to calculate FC directly (FDR was calculated using Benjamini-Hochberg function in R studio). The eggnog-mapper V2.0 was used to perform functional annotation based on Clusters of Orthologous Genes (COGs) (46). Further analysis of DE gene function was performed using Basic Local Alignment Search Tool (BLAST) (47–50), Phyre2 protein fold recognition (21) and contrastive learning-enabled enzyme annotation (CLEAN) (22). See Supplementary Information for further details.

### Data availability

RNA-seq data was deposited at NCBI (accession code XXXXXXX – pending assignment).

## Acknowledgments

We thank Dr. D. Schneider (Grenoble Alpes University) for vector pDS132, Dr. V. de Lorenzo (SEVA collection) for pSEVA227M, Dr. S. Golden (Addgene plasmid #120080) for pAM4891, and Dr. F. Thompson (Federal University of Rio de Janeiro) for the *P. brasiliensis* Ab134 strain. Financial support for this work was provided by the Division of Integrative Organismal Systems of the National Science Foundation under grant 1917492 (to ASE), and by FAPESP scholarships 2016/05133-7 and 2018/10742-8 (to LPI).

## Author contributions

Y.D. generated in-frame deletion mutants of *pppADGKLM*, with help from I.M. and L.A. for three of the mutants. V.L. generated in-frame deletion mutants of *pppEF* and L.P.I. generated in-frame deletion mutants of *pppHIJ*. Y.D. performed genetic complementation of Δ*pppEFGHIJ* mutants, with the help from D.A. who constructed plasmids for Δ*pppHIJ*. V.L. performed genetic complementation of the Δ*pppK* mutant. Y.D. and V.L. performed LC-MS analyses. Y.D. performed swarming assays, with the help from V.L. for the Δ*pppF* mutant. Y.D. extracted RNA and performed transcriptomics analyses. A.S.E and Y.D. designed experiments. A.S.E. obtained funding for the project and advised Y.D., V.L., L.P.I, D.A.S., I.M., and L.A. R.G.S.B. advised L.P.I. Y.D. advised V.L., D.A.S. I.M., and L.A. with vector construction and/or mutant generation. Y.D. wrote the initial paper draft. A.S.E. wrote the final paper. All authors commented on and approved the manuscript.

## Notes

### Competing Interest Statement

The authors have declared no competing interest.

