## Supplementary Information for "Pseudovibriamides from *Pseudovibrio* marine sponge bacteria promote swarming motility via transcriptional modulation"

### Supplementary Results

**Identifying DE genes potentially involved with reduced swarming motility.** Considering scenario 1 (the Δ*pppD* mutant possess the same set of genes unaffected compared to the wild type and regulated inversely in Δ*pppA* and Δ*pppE* mutants compared to the wild type), two of the three genes identified in the signal transduction [T] category not described in the main text are described here. One is predicted to encode YscD (locus tag: KGB56_13360), a type III secretion system (T3SS) inner membrane ring subunit (1). However, it is not near any genes that encode the remaining components of T3SS, and there is no direct relationship reported in the literature between YscD and swarming motility that we could find. The second gene (locus tag: KGB56_13020) upregulated in [T] is predicted to encode an amino acid periplasmic substrate binding protein of an ABC transporter. The substrate could be arginine, lysine, or histidine based on BLAST results. As described in the main text, upregulation of ABC transporters has been observed in non-swarming cells (2).

### Materials and Methods

**Chemicals and general experimental procedures.** Chemicals and enzymes were acquired from Sigma-Aldrich, VWR, BD, and Thermo Fisher Scientific, unless otherwise noted. Restriction enzymes were purchased from New England Biolabs (NEB). Phusion™ high-fidelity DNA polymerase (Thermo Fisher Scientific, referred as PHF polymerase later) and DreamTaq Master Mix 2× (Thermo Fisher Scientific) were used for PCR amplification. T4 DNA ligase (Thermo Fisher Scientific) was used for cloning. Oligonucleotide primers were designed in Geneious and synthesized by Sigma-Aldrich (see **Table S7** for detailed sequences). A Bio-Rad T100 thermal cycler was used for PCR reactions. The general PCR parameters for PHF polymerase reactions were: 60 s at 98°C; 30 cycles of 98°C for 10 s, Ta for 30 s, and 72°C for 30 s/kb; final extension for 10 mins at 72°C; and an indefinite hold at 10°C, and for DreamTaq polymerase reactions were: 3 mins at 95°C; 30 cycles of 95°C for 30 s, Ta for 30 s, and 72°C for 1 min/kb; final extension for 10 mins at 72°C; and an indefinite hold at 10°C. Genomic DNA was isolated using GenElute™ Bacterial Genomic DNA Kit (Sigma-Aldrich). ZymoPURE Plasmid Miniprep kits were used to extract plasmid DNA. The RiboPure™-Bacteria kit (Thermo Fisher Scientific) was used to extract RNA. Zymo Research Clean and Concentrator kit was used for DNA cleanup. To confirm the accuracy of plasmid constructs, Sanger sequencing was performed by UIC's genome research core and whole plasmid sequencing by Primordium labs.

**Plasmid construction.**

*General.* Ab134 genomic DNA was used as template for amplification of homology arm pairs (construction of pYD00A/D/G/K, pVL00E/F, pLI04/5/7, pYDORF1 and pLA002), a fragment containing predicted promoter of *pppA* (pYDproA), and fragments containing *pppG/H/I/J/K/E/F* genes respectively (pDS00H/I/J, pYDcompE/F/G, pVL00K). Phusion™ high-fidelity DNA polymerase (Thermo Scientific, referred as PHF polymerase later) was used for PCR amplification using primers listed in **Table S7** at 500 nM each unless otherwise noted. After restriction digests with the appropriate enzymes (**Table S7**), T4 DNA Ligase (Thermo Scientific) was used to ligate inserts and vector backbones. NEBuilder® HiFi DNA Assembly Master Mix was used for building pYDcompE/F/G.

*Construction of pYD004 to be used as vector to construct deletion plasmids.* PHF polymerase and the primer pair lacZ_F / lacZ_R (Ta = 63°C, 30 cycles) were used to amplify the *lacZ*α gene from pSET152 (Bierman, 1992). PCR products were digested with SpeI-HF on both ends and ligated into pDS132 digested with XbaI. LB agar (20 mL) supplemented with chloramphenicol (25 μg/mL) was treated with 5-bromo-4-chloro-3-indolyl-β-D-galactopyranoside (X-Gal, 0.8 mg/mL) and used to select *E. coli* DH5α (λpir) clones (blue colonies) that were further confirmed by using restriction digestion (EcoRV) and Sanger sequencing.

*Construction of pYD00A to generate ΔpppA strain.* One 895-bp upstream homology arm and one 888-bp downstream homology arm were amplified using PHF DNA polymerase and primer pairs pppA_Up_F / pppA_Up_R (Ta = 62°C, 30 cycles) and pppA_Down_F / pppA_Down_R (Ta = 62°C, 30 cycles) respectively. Both homology arms were assembled using splicing by overlap extension-PCR (SOE-PCR) with the following reaction (50 µL): purified homology arms (~100 ng), 0.2 mM dNTP mix (Thermo Scientific), 3% DMSO, 0.02 U/µL PHF polymerase with 1× PHF buffer, and molecular biology grade water. Extension reactions were performed using following parameters: 60 s at 98°C; 5 cycles of 98°C for 10 s, 60°C for 30 s, and 72°C for 90 s; and a final hold at 10°C. The primer pair pppA_Up_F / pppA_Down_R (Ta = 62°C) was added to the extension reaction before starting amplification: 60 s at 98°C; 30 cycles of 98°C for 10 s, 62°C for 30 s, and 72°C for 90 s; and a final extension for 10 mins at 72°C. The PCR product was digested with XbaI and SalI-HF and ligated into pYD004 digested with the same enzymes (located inside *lacZ*α gene). LB agar (20 mL) supplemented with chloramphenicol (25 µg/mL) was treated with X-Gal (0.8 mg/mL) to select *E. coli* DH5α(λpir) clones (white colonies as *lacZ*α gene was interrupted) that were further confirmed by using restriction digestion (SmaI) and Sanger sequencing.

*Construction of pYD00D to generate ΔpppD strain.* One 801-bp upstream homology arm and one 799-bp downstream homology arm were amplified using PHF DNA polymerase and primer pairs pppD_Up_F / pppD_Up_R (Ta = 61°C, 30 cycles) and pppD_Down_F_2 / pppD_Down_R (Ta = 61°C, 30 cycles) respectively. Both homology arms were assembled using SOE-PCR as described for pYD00A above. The same parameters of pYD00A were used for the first extension step except for the annealing temperature at 58°C for 30 s. The same parameters were also used for the second, amplification step except that the primer pair added was pppD_Up_F / pppD_Down_R (Ta = 61°C). The same restriction digestion, ligation, screening, and verification procedures of pYD00A were used.

*Construction of pVL00E to generate ΔpppE strain.* One 740-bp upstream homology arm and one 730-bp downstream homology arm were amplified using PHF DNA polymerase and primer pairs pppE_Up_F / pppE_Up_R (Ta = 62°C, 30 cycles) and pppE_Down_F / pppE_Down_R (Ta = 61°C, 30 cycles) respectively. Both homology arms were assembled using SOE-PCR as described for pYD00A above. The same parameters of pYD00A were used for the first thermal cycler extension. The same parameters were also used for the second amplification step except that the primer pair added was pppE_Up_R / pppE_Down_F (Ta = 61°C). Same restriction digestion, ligation, screening, and verification procedures of pYD00A were used.

*Construction of pVL00F to generate ΔpppF strain.* One 873-bp upstream homology arm and one 870-bp downstream homology arm were amplified using PHF DNA polymerase and primer pairs pppF_Up_F / pppF_Up_R (Ta = 63°C, 30 cycles) and pppF_Down_F / pppF_Down_R (Ta = 63°C, 30 cycles) respectively. Both homology arms were assembled using SOE-PCR with the reaction mentioned in pYD00A construction. The same parameters of pYD00A were used for the first thermal cycler extension except for the annealing temperature at 61°C for 30 s. The same parameters were also used for the second amplification step except for the primer pair added was pppF_Up_R / pppF_Down_F (Ta = 61°C). The PCR product was digested with XbaI and XhoI (compatible with SalI-HF) due to the presence of a SalI restriction site in the upstream homology arm. The same ligation, screening, and verification procedures of pYD00A were used.

*Construction of pYD00G to generate ΔpppG strain.* One 892-bp upstream homology arm and one 925-bp downstream homology arm were amplified using PHF DNA polymerase and primer pairs pppG_Up_F / pppG_Up_R (Ta = 61°C, 30 cycles) and pppG_Down_F / pppG_Down_R (Ta = 62°C, 30 cycles) respectively. Both homology arms were assembled using SOE-PCR with the reaction mentioned for pYD00A. The same parameters of pYD00A were used for the first thermal cycler extension except for the annealing temperature at 59°C for 30 s. The same parameters were also used for the second amplification step except for the primer pair added was pppG_Up_R / pppG_Down_F (Ta = 61°C). The same restriction digestion, ligation, and screening procedures of pYD00A were used. Positive clones were further confirmed by using restriction digestion (NcoI) and Sanger sequencing.

*Construction of pYD00K to generate ΔpppK strain.* One 977-bp upstream homology arm and one 921-bp downstream homology arm were amplified using PHF DNA polymerase and primer pairs pppK_Up_F / pppK_Up_R (Ta = 64°C, 30 cycles) and pppK_Down_F / pppK_Down_R (Ta = 66°C, 30 cycles) respectively. Both homology arms were assembled using SOE-PCR with the reaction mentioned for pYD00A. The same parameters of pYD00A were used for the first thermal cycler extension except that the annealing temperature was 64°C for 30 s. The same parameters were also used for the second amplification step except for the primer pair added was pppK_Up_R / pppK_Down_F (Ta = 66°C). The same ligation, screening, and verification procedures of pYD00G were used.

*Construction of pYDORF1 to generate ΔpppL(=ORF1) strain.* One 752-bp upstream homology arm and one 765-bp downstream homology arm were amplified using PHF DNA polymerase and primer pairs ORF1_Up_F_2 / ORF1_Up_R (Ta = 61°C, 30 cycles) and ORF1_Down_F_2 / ORF1_Down_R (Ta = 61°C, 30 cycles) respectively. Both homology arms were assembled using SOE-PCR with the reaction mentioned for pYD00A. The same parameters of pYD00A were used for the first thermal cycler extension except that the annealing temperature at 58°C for 30 s. The same parameters were also used for the second amplification step except for the primer pair added was ORF1_Up_F_2 / ORF1_Down_R (Ta = 61°C). Same ligation, and screening procedures of pYD00A were used. Positive clones were further confirmed by using double restriction digestion (XbaI and SalI-HF) and Sanger sequencing.

*Construction of pLA002 to generate ΔpppM (=ORF2) strain.* One 919-bp upstream homology arm and one 980-bp downstream homology arm were amplified using PHF DNA polymerase and primer pairs ORF2_Up_F / ORF2_Up_R (Ta = 61°C, 30 cycles) and ORF2_Down_F / ORF2_Down_R (Ta = 61°C, 30 cycles) respectively. Both homology arms were assembled using SOE-PCR with the reaction mentioned for pYD00A. The same parameters of pYD00A were used for the first thermal cycler extension except for the annealing temperature at 61°C for 30 s. The same parameters were also used for the second amplification step except for the primer pair added was ORF2_Up_F / ORF2_Down_R (Ta = 61°C). Same restriction digestion, ligation, screening, and verification procedures of pYDORF1 were used.

*Construction of pLI04 to generate ΔpppH strain.* One 951-bp upstream homology arm and one 905-bp downstream homology arm were amplified using PHF DNA polymerase and primer pairs oLI32 / oLI33 (Ta = 57°C, 30 cycles) and oLI34 / oLI35 (Ta = 60°C, 30 cycles) respectively. Both homology arms were assembled using SOE-PCR with the reaction mentioned for pYD00A. The same parameters of pYD00A were used for the first thermal cycler extension except for the annealing temperature at 62°C for 30 s. The same parameters were also used for the second amplification except for the primer pair added was oLI33 / oLI34 (Ta = 59°C). The PCR product was digested with XbaI and ligated into pDS132 digested with XbaI. LB agar (20 mL) supplemented with chloramphenicol (25 μg/mL) was used to select *E. coli* DH5α(λpir) clones that were further confirmed by using double restriction digestion (NcoI and BamHI) and Sanger sequencing.

*Construction of pLI05 to generate ΔpppI strain.* One 979-bp upstream homology arm and one 980-bp downstream homology arm were amplified using PHF DNA polymerase and primer pairs oLI36 / oLI37 (Ta = 61°C, 30 cycles) and oLI38 / oLI39 (Ta = 61°C, 30 cycles) respectively. Both homology arms were assembled using SOE-PCR with the reaction mentioned for pYD00A. The same parameters of pLI04 were used for the first thermal cycler extension. The same parameters were also used for the second amplification step except that the primer pair added was oLI37 / oLI38 (Ta = 61°C). The same restriction digestion, ligation, screening, and verification procedures of pLI04 were used. Clones were further confirmed by using restriction digestion (EcoRV-HF) and Sanger sequencing.

*Construction of pLI07 to generate ΔpppJ strain.* One 1014-bp upstream homology arm and one 994-bp downstream homology arm were amplified using PHF DNA polymerase and primer pairs oLI40 / oLI41 (Ta = 61°C, 30 cycles) and oLI46 / oLI43 (Ta = 61°C, 30 cycles) respectively. Both homology arms were assembled using SOE-PCR with the reaction mentioned for pYD00A. The same parameters of pLI04 were used for the first thermal cycler extension. Theame parameters were also used for the second amplification step except that the primer pair added was oLI41 / oLI46 (Ta = 61°C). The PCR product was digested with XbaI and SacI and ligated into pDS132 also digested with XbaI and SacI. LB agar (20 mL) supplemented with chloramphenicol (25 µg/mL) was used to select *E. coli* DH5α(λpir) clones that were further confirmed by using double restriction digestion (NcoI and BamHI) and Sanger sequencing. The same ligation and screening of pLI04 were used.

*Construction of pYDproA for the autoregulation test.* The promoterless pSEVA227M plasmid vector was used as the backbone with a monomeric superfolder green fluorescent protein (msfGFP) inserted as a reporter. One 216-bp upstream fragment of *pppA* gene containing a predicted promoter and a ribosome binding site was amplified using PHF DNA polymerase and the primer pair Pro_pppA_F / Pro_pppA_R (Ta = 57°C, 30 cycles). Both PCR product and pSEVA227M were digested with XbaI and HindIII, and the promoter fragment was inserted into a position 14-bp upstream of the *msfGFP* gene. After ligation, the mixture was chemically transformed into *E. coli* DH5α, and positive clones were selected by using LB agar (20 mL) supplemented with kanamycin (50 µg/mL) and further confirmed by using restriction digestion (AseI) and Sanger sequencing.

*Construction of pDS00H for pppH trans-complementation.* The 1141-bp fragment containing the *pppH* gene and ribosomal binding site (RBS) was amplified using PHF DNA polymerase and the primer pair pppH_comp_F / pppH_comp_R (Ta = 60°C, 30 cycles). The PCR product was digested with NdeI and BamHI (compatible with BglII) and ligated into pAM4891 digested with NdeI and BglII (located inside *the GFP* gene; 68% of the *GFP* gene was retained downstream of the insertion site). LB agar (20 mL) supplemented with kanamycin (50 µg/mL) was used to select

*E. coli* DH5α clones with lower intensity of green fluorescence and were further confirmed by using restriction digestion (SmaI) and Sanger sequencing.

*Construction of pDS00I for pppI trans-complementation.* The 831-bp fragment containing the *pppI* gene and RBS was amplified using PHF DNA polymerase and the primer pair pppI_comp_F / pppI_comp_R (Ta = 59°C, 30 cycles). The PCR product was digested with NdeI and BglII and ligated into the same sites of pAM4891. The same screening and verification procedures of pDS00H were used.

*Construction of pDS00J for pppJ trans-complementation.* The 723-bp fragment containing the *pppJ* gene and RBS was amplified using PHF DNA polymerase and the primer pair pppJ_comp_F / pppJ_comp_R (Ta = 60°C, 30 cycles). The same restriction digestion, ligation, screening, and verification procedures of pDS00I were used.

*Construction of pVL00K for pppK trans-complementation.* The 1017-bp fragment containing the *pppK* gene and RBS was amplified using PHF DNA polymerase and the primer pair pppK_F / pppK_R (Ta = 64°C, 30 cycles). The same restriction digestion, ligation, and screening procedures of pDS00I were used. Whole plasmid sequencing was used for verification.

*Construction of pYDcompE for pppE trans-complementation.* The 831-bp fragment containing the *pppE* gene and RBS was amplified using Q5 High-fidelity DNA polymerase and primer pairs pppE_comp_F / pppE_comp_R (Ta = 69°C, 30 cycles) following the manufacturer's instructions. A 7057-bp fragment excluding the *GFP* gene was amplified from pAM4891 using PHF DNA polymerase and the primer pair pAM4891_GA_F / pAM4891_GA_R (Ta = 61°C, 30 cycles). Gibson assembly was used for cloning (NEBuilder® HiFi DNA Assembly Master Mix) (Gibson, 2009). LB agar (20 mL) supplemented with kanamycin (50 µg/mL) was used to select *E. coli* DH5α clones with no green fluorescence that were further confirmed by using restriction digestion (XhoI and EcoRV) and whole plasmid sequencing.

*Construction of pYDcompF for pppF trans-complementation.* The 834-bp fragment containing the *pppF* gene and RBS was amplified using Q5 High-fidelity DNA polymerase and primer pairs pppF_comp_F / pppF_comp_R (Ta = 63°C, 30 cycles). The same 7057-bp fragment from pAM4891 as described above for pYDcompE was used for cloning. The same screening procedure was used to pick positive clones, which were further confirmed by using restriction digestion (BamHI and EcoRV) and whole plasmid sequencing.

*Construction of pYDcompG for pppG trans-complementation.* The 1280-bp fragment containing the *pppG* gene and RBS was amplified using Q5 High-fidelity DNA polymerase and primer pairs pppG_comp_F / pppG_comp_R (Ta = 65°C, 30 cycles). The same 7057-bp fragment from pAM4891 as described above for pYDcompE was used for cloning. The same screening procedure was used to pick positive clones, which were further confirmed by using restriction digestion (SmaI) and whole plasmid sequencing.

**Scarless in-frame deletion in Ab134.** Before conjugation, all pDS132 or pYD004 based suicide vectors were chemically transformed into *E. coli* SM10(λpir) using heat shock at 42°C for 45 s. For conjugation, fresh cultures of Ab134 or *E. coli* SM10(λpir) with suicide vectors were started by inoculating 400 μL of overnight seed culture into 20 mL of MB or 20 mL of LB containing chloramphenicol (25 μg/mL), respectively. The cultures were incubated at 30°C, 200 rpm, until an Optical Density at 600nm ($OD_{600nm}$) between 0.4 and 0.6 was reached. Cells were harvested by centrifugation at 4°C, 4000 rpm. Ab134 cells were resuspended into 2 mL of MB. *E. coli* SM10(λpir) cells were washed twice with 20 mL of LB to remove antibiotics prior to resuspension in 2 mL of LB. Cell suspensions of Ab134 and *E. coli* SM10(λpir) (500 μL each) were mixed in a 1.5-mL Eppendorf tube, and then plated evenly onto two MA plates. Negative control plates were prepared by plating only *E. coli* SM10(λpir) (250 μL) or only Ab134 (250 μL) onto MA plates. Plates were incubated at 30°C for 16-20 hours. A loopfull (10 μL loop) of cells were then streaked onto MA (20 mL) containing chloramphenicol (8 μg/mL) to select for the incoming plasmid and carbenicillin (50 μg/mL) to selectively kill *E. coli*. The wild type Ab134 strain was used as parent to generate mutants, except for Δ*pppD* only, for which fresh culture of Ab134 *pppD::neo* was used as the conjugation recipient (MB containing kanamycin 200 µg/mL). A loopfull of cells were then streaked onto MA (20 mL) containing kanamycin (200 µg/mL), chloramphenicol (8 µg/mL) to select for the incoming plasmid, and carbenicillin (50 µg/mL) to selectively kill *E. coli*. Positive clones (sticky single colonies) were analyzed by two parallel PCRs to identify single crossover (SCO) colonies using two primer pairs P1 / P3 and P4 / P2 (**Fig. S1**). The primer pairs used for each mutant were P1: TP_pppA_F / P3: pppA_Down_R (Ta = 50°C) and P4: pppA_Up_F / P2: TP_pppA_R (Ta = 51°C) (*pppA* SCO); TP_pppD_F_2 / pppD_Down_R (50°C) and pppD_Up_F / TP_pppD_R_4 (49°C) (*pppD* SCO); E_TP1_F / E_TP1_R (52°C) and E_TP2_F / E_TP2_R (52°C) (*pppE* SCO); F_TP1_F / F_TP1_R (52°C) and F_TP2_F / F_TP2_R (52°C) (*pppF* SCO); TP_pppG_F / pppG_Up_R (48°C) and pppG_Down_F / TP_pppG_R (49°C) (*pppG* SCO); TP_pppK_F / pppK_Up_R (53°C) and pppK_Down_F / TP_pppK_R (55°C) (*pppK* SCO); TP_ORF1_F_2 / ORF1_Down_R (49°C) and ORF1_Up_F_2 / TP_ORF1_R (51°C) (*pppL* SCO); and TP_ORF2_F / ORF2_Down_R (47°C) and ORF2_Up_F / TP_ORF2_R (50°C) (*pppM* SCO). For testing *pppH/I/J* SCO, P3 (pDS132_R) and P4 (pDS132_F) were located on pDS132. P1 and P2 were: oLI38 and JD_P2 (*pppH SCO*); oLI45 and oLI33 (*pppI SCO*); and oLI45 and oLI37 (*pppJ SCO*). All SCO PCR reactions were performed using the reaction mixture (20 µL): genomic DNA (~100 ng), 10 µL DreamTaq PCR Master Mix 2× (Thermo Scientific), both primers (250 nM each), and molecular biology grade water. Reactions were performed using following parameters: 60 s at 95°C; 30 cycles of 95°C for 30 s, Ta °C for 30s, and 72°C for 1min/kb; and a final extension for 10 mins at 72°C.

Once SCO colonies were confirmed, they were streaked evenly onto non-selective MA plates to allow double crossover (DCO) events to happen. Plates were incubated overnight at 30°C. A loopful of cells were then streaked for single colonies onto MA containing 5% sucrose (pDS132-based vectors have a *sacB* counterselection marker) and incubated for one to two days at 30°C. Obtained single colonies were replica plated onto MA or MA containing chloramphenicol

(8 µg/mL) except for potential Δ*pppD* colonies, which were replica plated onto MA or MA containing kanamycin (200 µg/mL), or MA containing chloramphenicol (8 µg/mL).

Chloramphenicol-sensitive clones (or kanamycin / chloramphenicol-sensitive clones for Δ*pppD*) were analyzed by colony PCR or PCR using genomic DNA (in case of poor colony PCR result) to confirm the gene replacement using the primer pair P1 / P2 (20 µL reaction, DreamTaq polymerase) for SCO verification. Thermocycling conditions were the same as those of SCO except that for the colony PCR the initial denaturation was extended from 60 s to 5 min at 95°C.

**Trans-complementation.** All pAM4891-based complementation vectors were first electroporated into *E. coli* S17-1, which was then used as the conjugation donor for introducing vectors into Ab134. The same conjugation protocol as for gene deletions was used, except the recipient strain is not Ab134 but the corresponding mutants. Selection of exconjugants was carried out on MA (20 mL) containing kanamycin (200 µg/mL) to select for the incoming plasmid and carbenicillin (50 µg/mL) to selectively kill *E. coli*. Positive clones (sticky single colonies) were inoculated into MB (5 mL) containing kanamycin (200 µg/mL) for plasmid extraction after overnight growth at 30°C. Plasmids from complemented strains were confirmed using the same restriction digestion protocol used in the plasmid construction step.

**Swarming assays.** The protocol used was as we previously reported (3) with slight modifications (the $OD_{600}$ of cultures used was 0.4 instead of 1.0). For convenience, we provide details here. A swarming agar with lower surface tension was prepared by mixing MB and 0.5% (*w/v*) of Eiken agar (Eiken Chemical CO., Japan). The autoclaved agar medium was equilibrated at 50°C for 2 h and aliquoted onto Petri dishes (exactly 20 mL), which were air dried for 20 mins in a laminar flow hood. Fresh cultures of Ab134 and mutants with $OD_{600nm}$ of 0.4-0.6 were normalized to $OD_{600nm}$ of 0.4. To perform the swarming assays, 5 µL of normalized cultures were carefully dropped on the center of the plate. Plates were air dried again for 10 min and incubated at 30°C (a water dish was placed in the incubator to maintain humidity. Pictures were taken daily for 4 days (when the swarming front had reached the plate edge). The swarming phenotype was apparent after two days.

**Pseudovibriamides extraction.** Pseudovibriamides were extracted in two ways based on culture conditions. (A) swarming agar plates were broken into pieces and extracted with one volume (20 mL) of methanol by sonicating for 1 h, after which the extract was filtered through filter paper. These extracts from WT, Δ*pppK*, and Δ*pppM* were analyzed with MALDI-ToF MS. (B) For all mutants and complemented Δ*pppH/I/J* strains, 1 mL of normalized seed culture based on $OD_{600nm}$ readings was added to 250 mL Erlenmeyer flasks containing 25 mL of MB in triplicates. After a 24-hour incubation at 30˚C, 200 rpm, exactly 10 mL of liquid cultures were extracted by first adding 5% w/v XAD-7HP resins (Sigma-Aldrich) directly to the culture, and shaking for 1 h at 500 rpm. The resin was collected by filtration and mixed with one volume methanol (10 mL), after which is was shaken for 1 h at 500 rpm. Methanol extracts were dried by the SpeedVac concentrator (Thermo Scientific) and stored at -20°C.

*Pseudovibriamides extraction from cell supernatant and pellet.* In addition to the whole culture extraction of Δ*pppG* and Δ*pppL* mutants, which were predicted to encode membrane transport proteins, supernatant and cell pellet were separated for extraction as shown in **SI Figure S24**. For the comparison between Δ*pppG* and WT, 1 mL of normalized seed culture based on $OD_{600nm}$ readings was added to 250 mL Erlenmeyer flasks with 50 mL of MB in triplicates and grown for 48 hours, 200 rpm. For the comparison between Δ*pppL* and WT, the same inoculation procedure was used but in quadruplicates. The same method as extracting liquid culture was used to extract exactly 30 mL of the supernatant portion, and the same method as extracting swarming agar plates was used to extract the corresponding cell pellet portion.

**MALDI-ToF MS sample preparation and analysis.** An Autoflex Speed LRF mass spectrometer (Bruker Daltonics) equipped with a SmartBeamTM-II laser (355 nm) was used for analysis. Dried-droplet analyses were performed in positive reflectron mode (Laser intensity: 70%-80%; Reflector: 15-18×; RepRate: 2000 Hz; ion source 1 voltage: 19 kV; ion source 2 voltage: 16.55 kV; lens voltage: 8.3 kV; mass range: 700 to 1500 Da). Samples (dissolved in methanol) and the calibrant were mixed in 1:1 ratio with universal matrix and 1 µL of the mixture was applied onto a designated spot on a MALDI ground-steel target plate (Bruker Daltonics). The universal matrix was prepared by mixing 1:1 of α- cyano-4-hydroxycinnamic acid (CHCA) and 2,5-dihydroxybenzoic acid (2,5-DHB) (Sigma-Aldrich) solubilized in 78:22 MeCN in water with 0.1% TFA. External calibration was made with a Bruker Daltonics peptide calibration standard using “PeptideCalibStandard mono” as the mass control list. Data was acquired by flexControl software v. 3.4.135.0 and analyzed using flexAnalysis software v. 3.4.

**UPLC-QToF-MS/MS sample preparation and analysis.** UPLC-MS/MS analyses were performed on a Bruker Elute LC system coupled to a Bruker Compact QToF mass spectrometer with electrospray ionization (ESI), using positive mode with following parameters: capillary voltage of 4500 V, nebulizer of 4.0 Bar, dry gas ($N_2$) flow of 12.0 L/min, source temperature of 225°C, scanning range from 50 to 3000 Da, and collision energy of 10 eV. Internal calibration was conducted with 5 mM sodium formate solution (Sigma-Aldrich) charged by the lock-mass probe at a flow rate of 40 µL/min. UPLC was performed at 0.5 mL/min through an Agilent Poroshell 120 EC-C18 UPLC column (2.1 × 50 mm × 1.9 µm) at 40°C. A solvent phase of aqueous 10% acetonitrile for 1 min (equilibration), 10% to 100% acetonitrile for 6 min (gradient), 100% acetonitrile for 2 min (wash), and 100% to 10% acetonitrile (re-equilibration) for 1 min was used. Both acetonitrile and water were LC-MS grade supplemented with 0.1% formic acid (v/v). Crude extracts were resuspended in LC-MS grade methanol and diluted to a final concentration of 1 mg/mL. Diluted samples were filtered using 4mm PTFE syringe filter 0.2 µm (Thermo Scientific) and injected into the system as 2 µL. Data was acquired and analyzed using Bruker Compass DataAnalysis software v. 5.1. For analyzing supernatant and cell pellet extracts, the area under curve (AUC) of each pseudovibriamide was obtained by using the extracted ion chromatogram (EIC) function. The ratio of AUC of pseudovibriamides in supernatant to AUC of pseudovibriamides in pellet was compared between Ab134 and Δ*pppG/L* mutants. The equation used for calculating export ratio is shown below:

$$Export\ ratio_{PA/B/C} = \frac{AUC_{supernatant}}{AUC_{pellet}}$$

**RNA extraction and transcriptomics.** The overall scheme is summarized in **Fig. S55**. Triplicate cultures of Ab134 and Δ*pppA/D/E* mutants were started by adding 1 mL of standardized overnight seed cultures ($OD_{600nm}$ ~ 1.0) to 50 mL of MB in 250 mL Erlenmeyer flasks and incubating at 30°C, 200 rpm. After 24 h or 48 h, 5 mL of culture of each triplicate was taken for metabolite analysis (extracted with 5% XAD-7HP resin [w/v]). Meanwhile, another 0.25 mL of culture, standardized based on $OD_{600nm}$ between samples so that an equivalent number of cells was collected, was centrifuged at 16,000 ×g for 2 min. Cell pellets were mixed with 0.25 mL of RNA-later® (Sigma-Aldrich) and stored at -20°C. RNA was isolated using the RiboPure™-Bacteria kit (Thermo) following manufacturer's instructions. The purified RNA was pre-screened by using a NanoDrop® ND-1000 UV-Vis Spectrophotometer. Although both 24 h and 48 h samples had all pseudovibriamides produced based on LC-MS/MS chromatograms, the RNA of 24 h samples had better quality than those of 48 h samples based on Nanodrop readings (**SI Table S8**). Thus, 24 h RNA samples were submitted to UIC's Genome Research Core for further quality control using Qubit® dsDNA HS (High Sensitivity) and Qubit® RNA BR (Broad-Range) assay kits (Thermo Fisher Scientific), and TapeStation 4200 (Agilent) (**SI Table S8**). Qualified samples (RIN, RNA integrity number > 8.0) were sent to SeqCenter (Pittsburgh, MA) for bacterial rRNA depletion RNA Sequencing. At SeqCenter, samples were DNAse treated with Invitrogen DNAse kit (Thermo

Fisher Scientific). Library preparation was performed using Illumina's Stranded Total RNA Prep Ligation with Ribo-Zero Plus kit and 10 bp IDT for Illumina indices. Sequencing was done on a NextSeq2000 instrument giving 2×51bp reads. Demultiplexing, quality control, and adapter trimming was performed with bcl-convert (v3.9.3). Raw Illumina reads of each sample were mapped to the Ab134 whole genome (NCBI accession number GCA_018282095.1) by using Geneious. Expression levels (FPKM, RPKM, and TPM) were calculated (all contigs at once) by using Geneious with the option that ambiguously mapped reads were count as partial matches. Pairwise differential expression analysis was performed by using DESeq2 package in Geneious or using TPM values to calculate FC directly (FDR was calculated using Benjamini-Hochberg function in R studio). The principal component analysis (PCA) plot, volcano plot, and heatmap were generated by using ggplot2 and ComplexHeatmap functions in R studio.

**Bioinformatics.** To check Δ*pppK* and Δ*pppG* strains for mutations, whole genome sequencing data (Illumina reads) were trimmed by BBDuk Trimmer and mapped to reference in Geneious Prime. Low coverage regions were anotated using Geneious Prime. Single nucleotide polymorphisms (SNPs) are found using Geneious Prime with the cutoff that the minimum vairant frequency is 0.90. SNPs that fell into low coverage regions were excluded.

**Visualization of Ab134 using transmission electron microscopy (TEM).** TEM sample preparation was modified from a previous study (Grossart, 2000). Seed cultures were prepared by growing 50 µL of cryo-preserved culture in 5 mL of MB at 30°C overnight (18-20 h). Cultures were preserved by adding 4% paraformaldehyde to a final concentration of 2%. 300-mesh copper TEM grids with Carbon-stabilized Formvar supports (FCF-300CU, Electron Microscopy Science) were used. Bacteria cells were adsorbed to TEM grids by floating the grids Formvar side down on an undiluted drop of culture (30 µL) for 5 min. Grids loaded with bacteria were stained by submerging into 1% phosphotungstic acid (30 µL) for 30 s and then were submerged in aliquots of Milli-Q water for rinsing three times (10 s each). Extra liquid of each step was removed gently by using filter paper. The grids were examined with a JEOL JEM-1400 Flash TEM (accelerating voltage of 80kV) at UIC's Electron Microscopy Core the same day of preparation.

**Growth curves.** The growth of each strain was monitored for 48 h using a microtiter plate method. Seed cultures were prepared by growing 50 µL of cryo-preserved culture in 5 mL of MB at 30°C overnight (18-20 h). Seed cultures were diluted in MB to $OD_{600nm}$ of 0.05. Standardized cultures (200 µL each well, 12 replicates for each strain) were aliquoted into BIOFLOAT® 96-well plates (faCellitate, U-bottom, inert, cell-repelling surface, sterile) using a multi-channel pipettor. Border wells were filled with MB (blank). Mutants were divided into two plates due to limited wells (Ab134 was in both plates as the control). BioTek Synergy HTX multimode reader (Agilent) was used to incubate plates at 30°C and shake the plate orbitally at speed of 425 cycle per minute (3 mm). $OD_{600nm}$ reading of each well was taken every 30 min (98 reads in total over 48 h). Half of the replicates without outlier readings were used to calculate average values and standard deviations for plotting the final growth curve.

**Pseudovibriamide autoregulation assay.** All strains (8 in total) were generated by introducing pYDproA (pSEVA227M based) or pSEVA227M (negative control) vectors into wild type/Δ*pppA*/Δ*pppD*/Δ*pppE* via using *E. coli* S17-1 as the conjugation donor. Before conjugation, both vectors were respectively transformed into *E. coli* S17-1 using electroporation (100 ng plasmid, 2.5kV, time constant 5 ms, 200Ω). For conjugation, the same protocol as conjugation of in-frame deletion vectors was used. Kanamycin (200 µg/mL) was used to select for the incoming plasmid and carbenicillin (50 µg/mL) to selectively kill *E. coli*. Plasmid DNA from kanamycin-resistant, single colonies were isolated and confirmed by using restriction digestion (AseI). Cultures of the eight strains were started by adding enough of overnight seed cultures (based on

$OD_{600nm}$ readings) to 50 mL of MB with kanamycin (200 µg/mL) in 250 mL Erlenmeyer flasks so that the starting $OD_{600nm}$ was standardized to 0.05. Cultures were incubated at 30°C, 200 rpm. Starting from T0, 300 µL of each culture was diluted into 2700 µL of salted PBS buffer (extra 8 g NaCl per liter added), and aliquoted into Corning Costar® 96-well plate (black plate, clear bottom with lid, 200 µL of diluted culture per well, 12 wells for each strain). Measurements of both $OD_{600nm}$ and GFP fluorescence were taken every 2 hours in a Tecan infinite® 200Pro plate reader (488 nm excitation and 520 nm emission). After cultures reached $OD_{600nm}$ 0.5 of (~6 hours), measurements were taken every one hour until 14 hours. The dilution ratio was raised from 1:10 to 1:50 (60 µL of each culture added to 3940 µL salted PBS buffer) for measurements taken at 24 hours, 26 hours, 28 hours, 38 hours, and 48 hours due to overly high fluorescence levels. All flasks were kept shaking for 48 hours except when samples were taken. The data was plotted using Excel.

### Tables

**Table S1. Putative functions of proteins encoded in the *ppp* BGC.**

| Gene | Protein size (a. a.) | BLAST analysis | Phyre2 analysis | Notes |
|---|---|---|---|---|
| *pppA* | 2766 | Non-ribosomal peptide synthetase | | Biosynthesis of peptides as modular enzymes. (4) |
| *pppB* | 3468 | | | |
| *pppC* | 1373 | | | |
| *pppD* | 2993 | Polyketide synthase/ non-ribosomal peptide synthetase hybrid | | Biosynthesis of hybrid polyketide-peptides as modular enzymes. (5) |
| *pppE* | 256 | 4'-phosphopantetheinyl transferase | | Transfer of phosphopantetheine from coenzyme A to ACP/PCP. (6) |
| *pppF* | 258 | Type II thioesterase | | Cleavage of aberrant intermediates or regenerating mis-acylated thiol groups. (7, 8) |
| *pppG* | 414 | Major Facilitator Superfamily (MFS) transporter; PTR2[a] | | Translocation of small solutes (di/tripeptides)[a] across the membrane. (9) |
| *pppH* | 349 | SGNH hydrolase superfamily | Acetyltransferase or acyltransferase | The SGNH hydrolase family includes esterases and acetyltransferases. (10) |
| *pppI* | 238 | Hypothetical protein; SGNH domain containing protein[a] | *O*-acetyltransferase AlgX/J (11) | |
| *pppJ* | 211 | Hypothetical protein | Hydrolase (low confidence) | |
| *pppK* | 304 | Fe(II)-oxoglutarate dependent dioxygenase | | Reactions include hydroxylation. (12) |
| *pppL* | 411 | MFS transporter; drug/proton antiporter[a] | | Same family as PppG; efflux of a variety of drugs and toxic compounds using proton gradient. (13) |
| *pppM* | 158 | GCN5-related *N*-acetyltransferase | | Acetyl transfer from acetyl-CoA to a substrate. (14) |
| *pppN* | 346 | ABC transporter periplasmic substrate-binding protein; AfuA[a] | | Recognition and delivery of a specific substrate for a specific ABC transporter. (15) |
| *pppO* | 462 | Sensor histidine kinase | | Part of a two-component system (TCS); |

| | | | |
|---|---|---|---|
| | | | Phosphorylation of response regulator transcription factor in response to signals. (16) |
| *pppP* | 222 | DNA-binding response regulator | Part of TCS. DNA binding and transcriptional control upon phosphorylation. (17) |

[a]Predicted by Conserved Domains in BLAST: PTR2, di/tripeptide permease; and AfuA, periplasmic component of $Fe^{3+}$ transport system. Low confidence, <20%; MFS, major facilitator superfamily; ABC, ATP-binding cassette; AlgJ/X, alginate *O*-acetyltransferase; and ACP/PCP, acyl/peptidyl carrier protein.

**Table S2. Summary of plasmids used in this study.**

| Plasmid | Description of features | Application | Reference |
|---|---|---|---|
| pDS132 | *ori* R6K (replication only in *E. coli* λpir); RP4 (conjugation system); $Cm^R$; *sacB* (a counter selectivion marker utilizing sucrose); MCS | Vector used to construct pYD004. | (18) |
| pYD004 | Based on pDS132*; ori* R6K; RP4; $Cm^R$; *lacZ*α (blue-white screening); *sacB*; MCS | pYD004 was constructed by introducing *lacZ*α from pSET152 into pDS132. Vector used to construct in-frame deletion plasmids. | pSET152 (19) pYD004 (this study) |
| pYD00A pYD00D pYD00G pYD00K | Based on pYD004; upstream and downstream homology arms of *pppA/D/G/K* cloned inside *lacZ*α | Delivery of replacement alleles for in-frame deletion into Ab134 cells. | This study |
| pVL00E pVL00F | Based on pYD004; upstream and downstream homology arms of *pppE/F* cloned inside *lacZ*α | Delivery of replacement alleles for in-frame deletion into Ab134 cells. | This study |
| pLI04 pLI05 pLI07 | Based on pDS132; upstream and downstream homology arms of *pppH/I/J* ligated in the MCS | Delivery of replacement alleles for in-frame deletion into Ab134 cells. | This study |
| pYDORF1 | Based on pYD004; upstream and downstream homology arms of *pppL* cloned inside *lacZ*α | Delivery of replacement alleles for in-frame deletion into Ab134 cells. | This study |
| pLA002 | Based on pYD004; upstream and downstream homology arms of *pppM* cloned inside *lacZ*α | Delivery of replacement alleles for in-frame deletion into Ab134 cells. | This study |
| pSEVA227M | *oriT* RK2 (*oriV, trfA;* broad host range); $Km^R$; *msfGFP* (promoterless) | Source of the GFP reporter gene and negative control in promoter-probe studies. | (20) |
| pYDproA | Based on pSEVA227M but with *msfGFP* under the control of the *pppA* promoter) | Test pseudovibriamide autoregulation of the $P_{pppA}$ promoter. | This study |
| pAM4891 | *Ori* RSF1010 (*oriV/T*, *repA/B,* broad host range); $Km^R$; *gfp; tac* promoter | Source of the backbone in construction of conjugatable, genetic complementation plasmids. | (21) |
| pVL00K | Based on pAM4891 but with *pppK* (native RBS included) inserted into *gfp* | Genetic complementation of the Δ*pppK* mutant. | This study |
| pYDcompE pYDcompF pYDcompG | Based on pAM4891; *gfp* replaced with *pppE/F/G* (native RBS included) | Genetic complementation of Δ*pppE/F/G* mutants, respectively. | This study |
| pDS00H pDS00I pDS00J | Based on pAM4891 but with *pppH/I/J* (native RBS included) inserted into *gfp* | Genetic complementation of Δ*pppH/I/J* mutants, respectively. | This study |

**ori*, origin of replication; *rep*, replication gene; R, resistance; Cm, chloramphenicol; Km, kanamycin; *msfGFP*, monomeric superfolder green fluorescent protein gene; and RBS, ribosome binding site.

**Table S3. Potential mutations identified in the Δ*pppG* mutant compared to the wild type.**

| Start (bp) | End (bp) | Nucleotide Change | Polymorphism Type | Variant % | Amino Acid Change | Locus tag | Protein | Protein Effect |
|---|---|---|---|---|---|---|---|---|
| **Δ*pppG* chromosome** | | | | | | | | |
| 3018287 | 3018286 | +C | Insertion | 100.0% | | | | |
| 3018285 | 3018285 | A -> G | SNP (transition) | 100.0% | | | | |
| 3018282 | 3018282 | A -> T | SNP (transversion) | 100.0% | | | | |
| 3018280 | 3018280 | A -> C | SNP (transversion) | 100.0% | | | | |
| 3018275 | 3018278 | AAAC -> TCTG | Substitution | 100.0% | | | | |
| 3018272 | 3018273 | AA -> GC | Substitution | 100.0% | | | | |
| 3018261 | 3018261 | A -> T | SNP (transversion) | 100.0% | | | | |
| 3018251 | 3018254 | AATA -> GCAT | Substitution | 100.0% | | | | |
| 3018247 | 3018247 | A -> T | SNP (transversion) | 100.0% | | | | |
| 3018244 | 3018245 | AA -> GC | Substitution | 100.0% | | | | |
| 3018209 | 3018209 | T -> G | SNP (transversion) | 100.0% | | | | |
| 550563 | 550563 | T -> C | SNP (transition) | 99.6% | T -> A | KGB56 _02520 | Hypothetical protein | Substitution |
| 3018270 | 3018270 | G -> T | SNP (transversion) | 99.5% | | | | |
| 3018256 | 3018259 | ACAA -> TGGC | Substitution | 99.5% | | | | |
| 3018249 | 3018249 | A -> T | SNP (transversion) | 99.5% | | | | |
| 3018288 | 3018290 | AAA -> TCT | Substitution | 99.4% | | | | |
| 3960649 | 3960649 | G -> T | SNP (transversion) | 99.3% | T -> K | KGB56 _17835 | CheB | Substitution |
| 3018263 | 3018263 | A -> T | SNP (transversion) | 99.0% | | | | |
| 3018265 | 3018268 | TATC -> GCAT | Substitution | 97.3% | | | | |
| **Plasmid #4** | | | | | | | | |
| 3951 | 3951 | A -> G | SNP (transition) | 100.0% | K -> R | KGB56 _26455 | PAPS | Substitution |
| 54820 | 54819 | +GCTTGGG | Insertion | 98.6% | | KGB56 _26670 | Hypothetical protein | Frame Shift |

*SNP, single nucleotide polymorphism; Variant %, frequency of variant; only SNPs with frequency of variant ≥ 90% are included; CheB, chemotaxis response regulator protein-glutamate methylesterase; PAPS, phosphoadenosine phosphosulphate reductase; SNPs without protein effect are in intergenic spaces; T, threonine; A, alanine; K, lysine; R, arginine.

**Table S4. Potential mutations identified in the Δ*pppK* mutant compared to the wild type.**

| Start (bp) | End (bp) | Nucleotide Change | Polymorphism Type | Variant % | Amino Acid Change | Locus tag | Protein | Protein Effect |
|---|---|---|---|---|---|---|---|---|
| **Δ*pppK* chromosome** | | | | | | | | |
| 3018287 | 3018286 | +C | Insertion | 100.0% | | | | |
| 3018285 | 3018285 | A -> G | SNP (transition) | 100.0% | | | | |
| 3018280 | 3018280 | A -> C | SNP (transversion) | 100.0% | | | | |
| 3018263 | 3018263 | A -> T | SNP (transversion) | 100.0% | | | | |
| 3018251 | 3018254 | AATA -> GCAT | Substitution | 100.0% | | | | |
| 2746853 | 2746864 | GTGGTCGTGGTC -> none | Deletion (tandem repeat) | 100.0% | HDHD -> none | KGB56_12280 | CobW | Deletion |
| 3018247 | 3018247 | A -> T | SNP (transversion) | 99.7% | | | | |
| 3018244 | 3018245 | AA -> GC | Substitution | 99.7% | | | | |
| 550563 | 550563 | T -> C | SNP (transition) | 99.7% | T -> A | KGB56_02520 | Hypothetical protein | Substitution |
| 3018270 | 3018270 | G -> T | SNP (transversion) | 99.6% | | | | |
| 3018265 | 3018268 | TATC -> GCAT | Substitution | 99.6% | | | | |
| 3018261 | 3018261 | A -> T | SNP (transversion) | 99.6% | | | | |
| 3018249 | 3018249 | A -> T | SNP (transversion) | 99.6% | | | | |
| 3018209 | 3018209 | T -> G | SNP (transversion) | 99.5% | | | | |
| 3960649 | 3960649 | G -> T | SNP (transversion) | 99.4% | T -> K | KGB56_17835 | CheB | Substitution |
| 3018275 | 3018278 | AAAC -> TCTG | Substitution | 99.2% | | | | |
| 3018272 | 3018273 | AA -> GC | Substitution | 99.2% | | | | |
| 3018288 | 3018290 | AAA -> TCT | Substitution | 99.1% | | | | |
| 3018282 | 3018282 | A -> T | SNP (transversion) | 99.1% | | | | |
| 3018256 | 3018259 | ACAA -> TGGC | Substitution | 98.9% | | | | |

*SNP, single nucleotide polymorphism; Variant %, frequency of variant; only SNPs with frequency of variant ≥ 90% are included; CobW, cobalamin biosynthesis protein; CheB, chemotaxis response regulator protein-glutamate methylesterase; SNPs without protein effect are in intergenic spaces; H, histidine; D, aspartic acid; T, threonine; A, alanine; K, lysine.

**Table S5. List of DE genes in Δ*pppA/E* mutants but NDE in Δ*pppD* mutant compared to the wild type.**

| Description | Locus tag | A $\log_2$FC | A FDR | E $\log_2$FC | E FDR | D $\log_2$FC | $\lvert\Delta\log_2FC\rvert$ A_D | $\lvert\Delta\log_2FC\rvert$ E_D | COG |
|---|---|---|---|---|---|---|---|---|---|
| **Upregulation** | | | | | | | | | |
| hypothetical protein CDS | KGB56_ 25290 | 2.06 | 0.0053 | 1.57 | 0.0039 | 0.90 | 1.16 | 0.66 | S |
| VOC family protein CDS | KGB56_ 25440 | 1.93 | 0.0062 | 1.89 | 0.0012 | -0.55 | 2.48 | 2.44 | E |
| FAD-dependent oxidoreductase CDS | KGB56_ 25450 | 1.78 | 0.0051 | 1.70 | 0.0034 | -0.61 | 2.39 | 2.31 | CH |
| TRAP transporter substrate-binding protein CDS | KGB56_ 25435 | 1.78 | 0.0057 | 1.59 | 0.0056 | -0.91 | 2.69 | 2.50 | G |
| cupin domain-containing protein CDS | KGB56_ 07995 | 1.57 | 0.0035 | 2.15 | 0.0017 | -0.32 | 1.90 | 2.47 | S |
| ABC transporter substrate-binding protein CDS | KGB56_ 22975 | 1.53 | 0.0061 | 1.16 | 0.0073 | 0.64 | 0.89 | 0.52 | P |
| cyclic nucleotide-binding/CBS domain-containing protein CDS | KGB56_ 23400 | 1.41 | 0.0053 | 1.28 | 0.0067 | -0.22 | 1.63 | 1.50 | T |
| amino acid ABC transporter ATP-binding protein CDS | KGB56_ 12375 | 1.27 | 0.0077 | 1.16 | 0.0070 | 0.55 | 0.72 | 0.61 | E |
| SIS domain-containing protein CDS | KGB56_ 04810 | 1.25 | 0.0029 | 1.27 | 0.0055 | 0.34 | 0.91 | 0.92 | M |
| hypothetical protein CDS | KGB56_ 25995 | 1.19 | 0.0028 | 1.13 | 0.0040 | 0.39 | 0.80 | 0.74 | S |
| cadherin-like domain-containing protein CDS | KGB56_ 13365 | 1.07 | 0.0041 | 1.28 | 0.0037 | 0.49 | 0.58 | 0.79 | Q |
| type III export protein YscD CDS | KGB56_ 13360 | 1.02 | 0.0063 | 1.21 | 0.0023 | 0.22 | 0.80 | 0.99 | T |
| **Downregulation** | | | | | | | | | |
| transglutaminase-like cysteine peptidase CDS | KGB56_ 14760 | -3.04 | 0.0044 | -2.93 | 0.0039 | -0.49 | 2.55 | 2.43 | S |
| peptidoglycan DD-metalloendopeptidase family protein CDS | KGB56_ 11125 | -2.12 | 0.0009 | -2.13 | 0.0009 | -0.51 | 1.61 | 1.61 | M |
| methyltransferase domain-containing protein CDS | KGB56_ 20140 | -2.11 | 0.0010 | -1.85 | 0.0010 | -0.96 | 1.15 | 0.89 | Q |
| argininosuccinate synthase CDS | KGB56_ 20790 | -1.93 | 0.0012 | -1.12 | 0.0094 | -0.43 | 1.50 | 0.69 | E |
| mutT CDS | KGB56_ 20150 | -1.35 | 0.0087 | -1.33 | 0.0054 | -0.56 | 0.79 | 0.77 | L |

* A, Δ*pppA* mutant; E, Δ*pppE* mutant; D, Δ*pppD* mutant; FC, fold change when compared to wild type; $\lvert\Delta\log_2FC\rvert$, absolute values obtained from $\log_2$FC of Δ*pppA/E* mutants minus $\log_2$FC of Δ*pppD* mutant; COG, cluster of orthologous groups.

**Table S6. List of specified DE genes in Δ*pppA/E* mutants but inversely regulated in Δ*pppD* mutant compared to wild type.**

| Description | Locus tag | A $\log_2$FC | A FDR | E $\log_2$FC | E FDR | D $\log_2$FC | COG |
|---|---|---|---|---|---|---|---|
| **Upregulation** | | | | | | | |
| hypothetical protein CDS | KGB56_00075 | 2.20 | 0.0079 | 2.19 | 0.0029 | -3.90 | S |
| HlyD family type I secretion periplasmic adaptorsubunit CDS | KGB56_04835 | 2.08 | 0.0030 | 2.33 | 0.0046 | -2.39 | M |
| basic amino acid ABC transporter substrate-binding protein CDS | KGB56_13020 | 1.95 | 0.0041 | 1.24 | 0.0021 | -1.31 | ET |
| NUDIX hydrolase CDS | KGB56_03310 | 1.68 | 0.0073 | 1.10 | 0.0023 | -1.17 | L |
| type I secretion system permease/ATPase CDS | KGB56_04840 | 1.68 | 0.0008 | 2.26 | 0.0048 | -2.67 | V |
| LuxR family transcriptional regulator CDS | KGB56_18890 | 1.54 | 0.0050 | 1.48 | 0.0006 | -2.78 | K |
| YifB family Mg chelatase-like AAA ATPase CDS | KGB56_21835 | 1.48 | 0.0037 | 1.85 | 0.0006 | -3.28 | O |
| hypothetical protein CDS | KGB56_24120 | 1.48 | 0.0067 | 2.04 | 0.0021 | -2.30 | S |
| lipid-A-disaccharide synthase N-terminal domain-containing protein CDS | KGB56_12395 | 1.25 | 0.0084 | 1.47 | 0.0031 | -3.88 | S |
| PaaI family thioesterase CDS | KGB56_22365 | 1.13 | 0.0027 | 1.51 | 0.0073 | -3.84 | Q |
| type III polyketide synthase CDS | KGB56_24430 | 1.04 | 0.0070 | 1.57 | 0.0096 | -4.24 | Q |
| MoxR family ATPase CDS | KGB56_16960 | 1.02 | 0.0061 | 1.12 | 0.0028 | -2.24 | S |

* A, Δ*pppA* mutant; E, Δ*pppE* mutant; D, Δ*pppD* mutant; FC, fold change when compared to wild type; COG, cluster of orthologous groups.

**Table S7. Oligonucleotide primers used in this study.**

| Primer name | Sequence from 5' to 3' | Additional Info |
|---|---|---|
| *lacZ* gene amplification from pSET152 | | |
| lacZ_F | *GG*ACTAGTTCTCCGACCTGATGCAGC | SpeI |
| lacZ_R | *GG*ACTAGTGCCTTTGAGTGAGCTGATACC | SpeI |
| *pppA* gene in-frame deletion and verification | | |
| pppA_Up_F | *ACGC*GTCGACAGCTACCTTCAACAACCTCG | SalI |
| pppA_Up_R | TACGTTGGAAAGACTAAACTCACC | |
| pppA_Down_F | GGTGAGTTTAGTCTTTCCAACGTAGCCGAAGAATTTTCCTGACG | |
| pppA_Down_R | *GC*TCTAGACTCACATGCAAACTCTCTTACC | XbaI |
| TP_pppA_F | AGGAAGAACGTGAGAGGC | |
| TP_pppA_R | ATCTCTTCCAGCTCTTTACGG | |
| *pppD* gene in-frame deletion and verification | | |
| pppD_Up_F | *ACGC*GTCGACAGATGCACAAGACAACCTG | SalI |
| pppD_Up_R | ATGTGAGATAGTGCTGTTCATG | |
| pppD_Down_F_2 | CATGAACAGCACTATCTCACATGTGCTCAGTGTCATGGAC | |
| pppD_Down_R | *GC*TCTAGAGATGTCTCAACCCGCTAC | XbaI |
| TP_pppD_F_2 | TAAACCCAACAATCGAAGCG | |
| TP_pppD_R_4 | AGCAAAGTGGAAGAATGGAC | |
| *pppE* gene in-frame deletion and verification | | |
| pppE_Up_F | CCAGCGAAGAGGAACATCAAAATGTTGCCACTTCCCTCTC | |
| pppE_Up_R | *GC*TCTAGATTGGTGATCAGGTGAACGTG | XbaI |
| pppE_Down_F | *ACGC*GTCGACTGACCCTCACCACAGTAAC | SalI |
| pppE_Down_R | TTTGATGTTCCTCTTCGCTGG | |
| E_TP1_F | TGCCGTAACTGTACATGAGTG | |
| E_TP1_R | TTGGTGATCAGGTGAACGTG | |
| E_TP2_F | TGACCCTCACCACAGTAAC | |
| E_TP2_R | TGAGTTTTCCGATGCCTTCC | |
| *pppF* gene in-frame deletion and verification | | |
| pppF_Up_F | CGTCCATTCTTCCACTTTGCTTGGCATCACGTTCACCTG | |
| pppF_Up_R | *GC*TCTAGAAGCAAAGCCCGTCTCAAC | XbaI |
| pppF_Down_F | *CCG*CTCGAGAGCTGTAACACCTGGATCAAC | XhoI |
| pppF_Down_R | AGCAAAGTGGAAGAATGGACG | |
| F_TP1_F | AACTGGCTGGGAGTTCTTC | |
| F_TP1_R | AGCAAAGCCCGTCTCAAC | |
| F_TP2_F | AGCTGTAACACCTGGATCAAC | |
| F_TP2_R | ACGGCACTGTTCTGGTTG | |
| *pppG* gene in-frame deletion and verification | | |
| pppG_Up_F | GGTATACACAGCATCGCAGAAAGCATCCGGAATTGACG | |
| pppG_Up_R | *GC*TCTAGACAGTACTCGTGCATTCTCG | XbaI |
| pppG_Down_F | *ACGC*GTCGACTGCTTCCAACTGACCAGC | SalI |
| pppG_Down_R | TCTGCGATGCTGTGTATACC | |
| TP_pppG_F | AGGATGGAAAGTACTCAACG | |
| TP_pppG_R | TTCAGATATGGATGCGTCG | |
| *pppH* gene in-frame deletion and verification | | |
| oLI32 | GTCTTCTTCAGCAGTTTTTTGTTGGTATGGGTTCATTTTACACC | |
| oLI33 | *AACTG*TCTAGACAGTCATCATCGTGATGTC | XbaI; Pairs with oLI32 |
| oLI34 | *AACTG*TCTAGACATCTGAGCGGCTTCTAT | XbaI |

| | | |
|---|---|---|
| oLI35 | CAACAAAAAACTGCTGAAGAGA | Pairs with oLI34 |
| pDS132_F | AGTGAACGGCAGGTATATGTG | |
| pDS132_R | TGTGGAATTGTGAGCGGATAAC | |
| JD_P2 | CAACTTGCTCTATTCGGGTC | Pairs with oLI38 below |
| *pppI* gene in-frame deletion and verification | | |
| oLI36 | CTGACCAAACGCTACGTTCCCATGGGCATGAGAATCTCCAAGCA | |
| oLI37 | *AACTG*TCTAGAGATCTCCAAAGATGCTACGATC | XbaI; Pairs with oLI36 |
| oLI38 | *AACTG*TCTAGACGTAGCTCTCCAAACTTTCG | XbaI |
| oLI39 | CATGGGAACGTAGCGTTTG | Pairs with oLI38 |
| oLI45 | GCAGATTGCGAGTGTAAGGAAAG | Pairs with oLI33 (*pppI*) or oLI37 (*pppJ*) |
| *pppJ* gene in-frame deletion | | |
| oLI40 | AAGGCTGAGTAAAGCAACCTCGTCGGTTTCAAGCTGAGGTCG | |
| oLI41 | *AACTG*TCTAGAGACAGCCTCTACTTACACCA | XbaI; Pairs with oLI40 |
| oLI46 | *AACTG*GAGCTCTGGTCATCAACCTAGAACTCC | SacI |
| oLI43 | GACGAGGTTGCTTTACTCAG | Pairs with oLI46 |
| *pppK* gene in-frame deletion and verification | | |
| pppK_Up_F | GGATGCGGCAGTTCAGAGCAAGATCTACACCACAAATTTCGGC | |
| pppK_Up_R | *GC*TCTAGATCGTATGGGCTTGAAAACTCTGG | XbaI |
| pppK_Down_F | *ACGC*GTCGACAGCCAACAAAGGAAGCGTGG | SalI |
| pppK_Down_R | GCTCTGAACTGCCGCATCC | |
| TP_pppK_F | TGATCGTCTCTATGGGTATCAAAGC | |
| TP_pppK_R | GCTGTTTCACGACCTTATCTGCG | |
| *pppL* gene in-frame deletion and verification | | |
| ORF1_Up_F_2 | *ACGC*GTCGACCAACGGAACAGCTCAACG | SalI |
| ORF1_Up_R | AGAATAGCGGTCAATCAACG | |
| ORF1_Down_F_2 | CGTTGATTGACCGCTATTCTGTGCTCCAGAAACTCGTC | |
| ORF1_Down_R | *GC*TCTAGACTATGAGATGTTATCCCCTGC | XbaI |
| TP_ORF1_F_2 | GCTAGAAATCTCCGATGCC | |
| TP_ORF1_R | AAGTCTCGTTTTGTAGCCG | |
| *pppM* gene in-frame deletion and verification | | |
| ORF2_Up_F | *ACGC*GTCGACAGGTGAAACTGGCAAACG | SalI |
| ORF2_Up_R | GTAGTAATCCTTCGCACCG | |
| ORF2_Down_F | CGGTGCGAAGGATTACTACTCAGACATCAGCCTTGCC | |
| ORF2_Down_R | *GC*TCTAGAGATGCCCAGTTTCAAAATGC | XbaI |
| TP_ORF2_F | GTTTCCATCGATTACACCG | |
| TP_ORF2_R | CAGGCAAAACAGGTGAGC | |
| Native promoter of *pppA* gene amplification | | |
| Pro_pppA_F | *GC*TCTAGAATGAACAAAACAACCTGAG | XbaI |
| Pro_pppA_R | *CCC*AAGCTTACAACTACCGATGATTGAG | HindIII |
| *pppH* trans-complementation | | |
| pppH_comp_F | *GGAATTC*CATATGTCTCCAAGCAGCAAGAAC | NdeI |
| pppH_comp_R | *CG*GGATCCGAGGAGGTCGCATGATATC | BamHI |
| *pppI* trans-complementation | | |
| pppI_comp_F | *GGAATTC*CATATGGTCATACATGCCGCTAAG | NdeI |
| pppI_comp_R | *GA*AGATCTAAACTGCTGAAGAAGACAC | BglII |
| *pppJ* trans-complementation | | |
| pppJ_comp_F | *GGAATTC*CATATGATGCATGACGTTTCCTCC | NdeI |
| pppJ_comp_R | *GA*AGATCTCTCGATGCATTCACCAAC | BglII |
| *pppK* trans-complementation | | |
| pppK_F | *GGAATTC*CATATGTCAACCTAGAACTCCCACC | NdeI |

| | | |
|---|---|---|
| pppK_R | *CGG*GGATCCAGCAGCTCACTTGTACATCAG | BamHI |
| pAM4891 backbone (Gibson assembly) | | |
| pAM4891_GA_F | AGAAGGCCATCCTGACGG | |
| pAM4891_GA_R | AGATCTGGGTACCATTATACGAG | |
| *pppG* trans-complementation (Gibson assembly) | | |
| pppG_comp_F | ATAATGGTACCCAGATCTCTCAGGTTGTTTTGTTCATGTCTG | |
| pppG_comp_R | CCGTCAGGATGGCCTTCTTCACATGACGCGAACTAGATATC | |
| *pppE* trans-complementation (Gibson assembly) | | |
| pppE_comp_F | ATAATGGTACCCAGATCTTGGCGTAAGACACAATGCCG | |
| pppE_comp_R | CCGTCAGGATGGCCTTCTGAGGGGTGAGGCGCTAGAT | |
| *pppF* trans-complementation (Gibson assembly) | | |
| pppF_comp_F | ATAATGGTACCCAGATCTTTTTAAATGAGTTTTCCGATGCCT | |
| pppF_comp_R | CCGTCAGGATGGCCTTCTTGTGTCTTACGCCAGCTG | |

*SOE-PCR or Gibson assembly overhangs are shown in blue, restriction sites in red and underlined. Random bases inserted to allow restriction of PCR fragments are in *italics*.

**Table S8. Summary of RNA sample quality used in this study.**

| Sample | Nanodrop conc. (ng/μL) | 260nm /280nm | 260nm /230nm | Qubit conc. (ng/μL) | dsDNA conc. (ng/μL) | %dsDNA | RIN |
|---|---|---|---|---|---|---|---|
| **24 h samples** | | | | | | | |
| W1 | 54.8 | 2.15 | 2.28 | 176 | 3.48 | 1.98% | 8.3 |
| W2 | 50.9 | 2.06 | 2.44 | 159 | 3.04 | 1.91% | 8.5 |
| W3 | 48.8 | 2.07 | 2.35 | 154 | 2.26 | 1.47% | 8.1 |
| A1 | 68.7 | 2.35 | 3.45 | 228 | 6.34 | 2.78% | 8.4 |
| A2 | 47.2 | 2.33 | 1.22 | 180 | 3.2 | 1.78% | 8.3 |
| A3 | 43.2 | 2.53 | 0.4 | 190 | 3.14 | 1.65% | 8.6 |
| D1 | 68.3 | 2.29 | 3.09 | 226 | 1.21 | 0.54% | 8.5 |
| D2 | 29.8 | 2.6 | 2.67 | 126 | 1.41 | 1.12% | 8.6 |
| D3 | 46.5 | 2.43 | 4.01 | 173 | 3.2 | 1.85% | 8.6 |
| E1 | 40.7 | 2.54 | 5.41 | 168 | 2.12 | 1.26% | 8.6 |
| E2 | 37.5 | 2.46 | 0.93 | 149 | 1.92 | 1.29% | 8.8 |
| E3 | 78.4 | 2.28 | 3.28 | 268 | 7.22 | 2.69% | 8.8 |
| **48 h samples (Nanodrop readings only)** | | | | | | | |
| W1 | 42.2 | 2.23 | 2.14 | | | | |
| W2 | 17.1 | 2.38 | 0.17 | | | | |
| W3 | 9.6 | 1.99 | 1.39 | | | N/A | |
| A1 | 35.2 | 2.20 | 2.20 | | | | |
| D1 | 22.1 | 2.14 | 1.84 | | | | |
| E1 | 14.0 | 2.23 | 1.15 | | | | |

dsDNA: double-stranded DNA; RIN: RNA integrity number.

### Figures

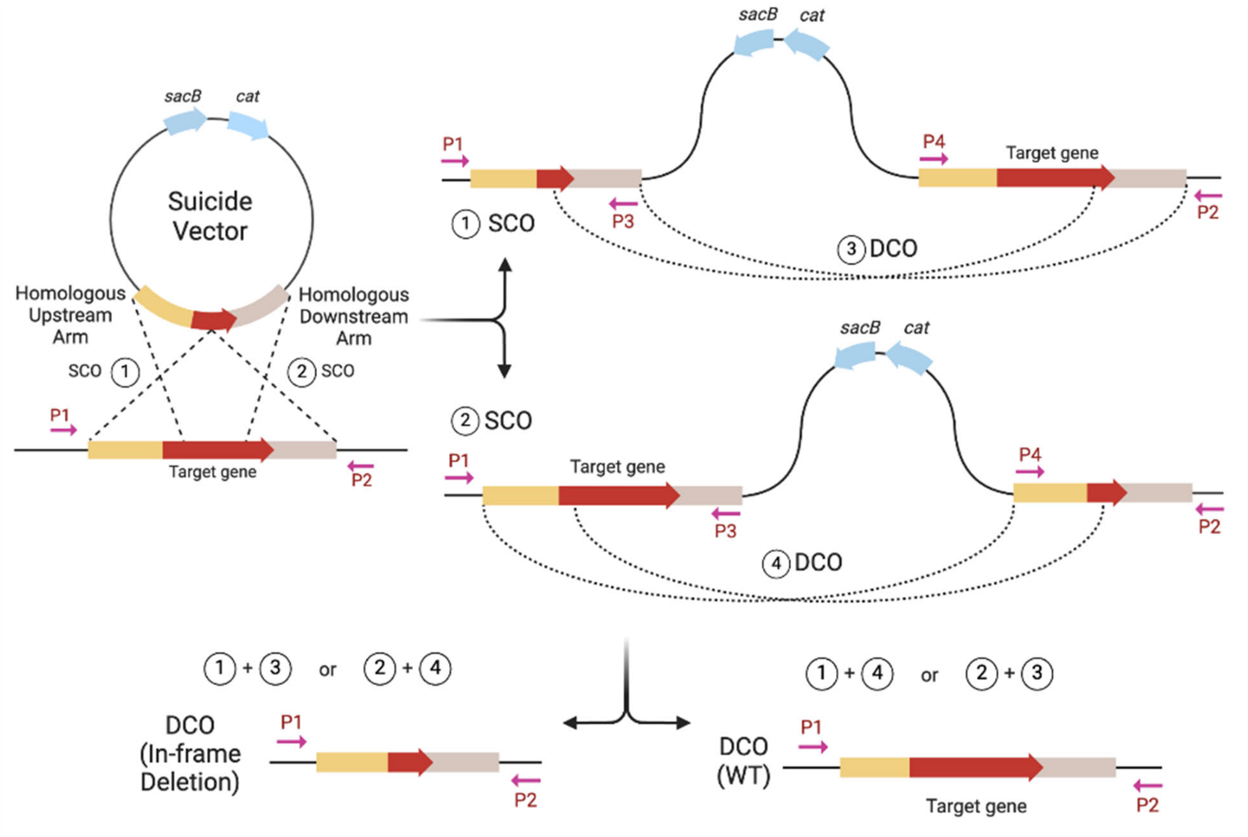

**Figure S1. Scheme of in-frame deletion via homologous recombination.** The first crossover event can happen at each of the two homologous sequences as indicated (single crossover, SCO 1 and 2). SCO mutants are obtained by selection on chloramphenicol and confirmed by PCR (not shown). The obtained SCO mutants are then grown on agar media without antibiotics but containing sucrose for counterselection of the vector backbone. The second crossover event (double crossover, DCO) can again happen at either arm resulting either in the desired in-frame deletion or in reversion to the wild type as indicated. P1, P2, P3, and P4 are primers used to verify SCO and DCO mutants by PCR. *sacB*, gene encoding levansucrase; *cat*, chloramphenicol acetyltransferase resistance gene.

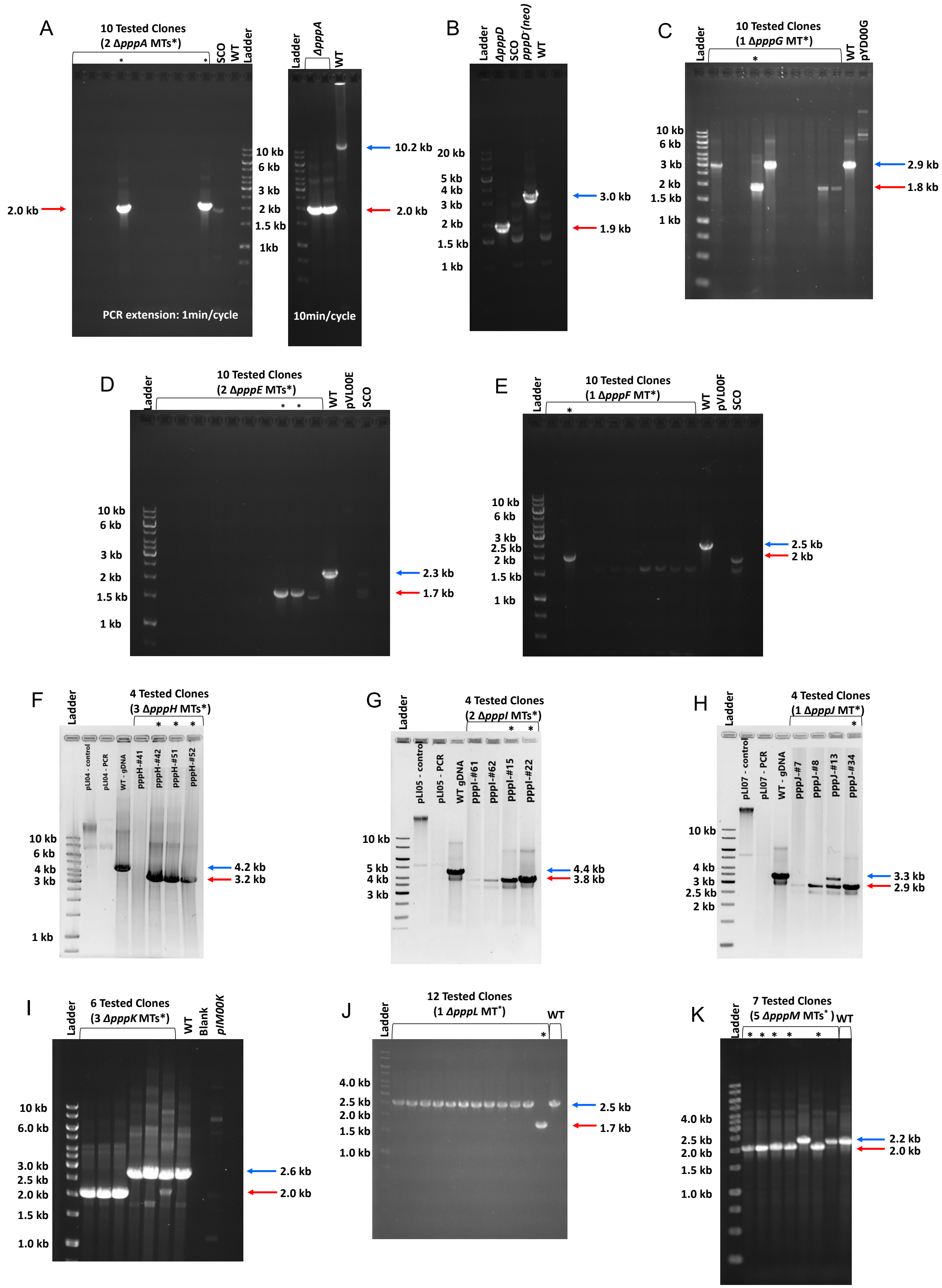
A
10 Tested Clones
(2 ΔpppA MTs*)
SCO
WT
Ladder
2.0 kb
10 kb
6 kb
3 kb
2 kb
1.5 kb
1kb
PCR extension: 1min/cycle
Ladder
ΔpppA
WT
10.2 kb
2.0 kb
10min/cycle
B
Ladder
ΔpppD
SCO
pppD'(neo)
WT
20 kb
5 kb
4 kb
3 kb
2 kb
1.5 kb
1 kb
3.0 kb
1.9 kb
C
Ladder
10 Tested Clones
(1 ΔpppG MT*)
WT
pYD00G
10 kb
6 kb
3 kb
2 kb
1.5 kb
1 kb
2.9 kb
1.8 kb
D
Ladder
10 Tested Clones
(2 ΔpppE MTs*)
WT
pVL00E
SCO
10 kb
6 kb
3 kb
2 kb
1.5 kb
1 kb
2.3 kb
1.7 kb
E
Ladder
10 Tested Clones
(1 ΔpppF MT*)
WT
pVL00F
SCO
10 kb
6 kb
3 kb
2.5 kb
2 kb
1.5 kb
1 kb
2.5 kb
2 kb
F
Ladder
4 Tested Clones
(3 ΔpppH MTs*)
pLI04 - control
pLI04 - PCR
WT - gDNA
pppH-#41
pppH-#42
pppH-#51
pppH-#52
10 kb
6 kb
4 kb
3 kb
1 kb
4.2 kb
3.2 kb
G
Ladder
4 Tested Clones
(2 ΔpppI MTs*)
pLI05 - control
pLI05 - PCR
WT gDNA
pppI-#61
pppI-#62
pppI-#15
pppI-#22
10 kb
5 kb
4 kb
3 kb
4.4 kb
3.8 kb
H
Ladder
4 Tested Clones
(1 ΔpppJ MT*)
pLI07 - control
pLI07 - PCR
WT - gDNA
pppJ-#7
pppJ-#8
pppJ-#13
pppJ-#34
10 kb
4 kb
3 kb
2.5 kb
2 kb
3.3 kb
2.9 kb
I
Ladder
6 Tested Clones
(3 ΔpppK MTs*)
WT
Blank
pIM00K
10 kb
6.0 kb
3.0 kb
2.5 kb
2.0 kb
1.5 kb
1.0 kb
2.6 kb
2.0 kb
J
Ladder
12 Tested Clones
(1 ΔpppL MT*)
WT
4.0 kb
2.5 kb
2.0 kb
1.5 kb
1.0 kb
2.5 kb
1.7 kb
K
Ladder
7 Tested Clones
(5 ΔpppM MTs*)
WT
4.0 kb
2.5 kb
2.0 kb
1.5 kb
1.0 kb
2.2 kb
2.0 kb

**Figure S2. Gel electrophoresis images of scarless in-frame deletion mutants verified by PCR.** Comparison of PCR amplification products between in-frame deletion mutants (MTs) and the wild type (WT) (**A**) Δ*pppA*. Here two PCRs were performed, one with a shorter extension (left) and another with a longer extension time required for amplifying the WT *pppA* (right). (**B**) Δ*pppD* . The Δ*pppD* mutant was generated from the *pppD::neo* mutant as parent. No band would show up for the wild type due to the short extension time used. (**C**) Δ*pppG*. (**D**) Δ*pppE*. (**E**) Δ*pppF*. (**F**) Δ*pppH*. (**G**) Δ*pppI*. (**H**) Δ*pppJ*. (**I**) Δ*pppK*. (**J**) Δ*pppL*. (**K**) Δ*pppM*. An asterisk (*) indicates positive in-frame deletion clones. The blue arrows indicate the size of wild type bands, and the red arrows the size of expected mutant bands.

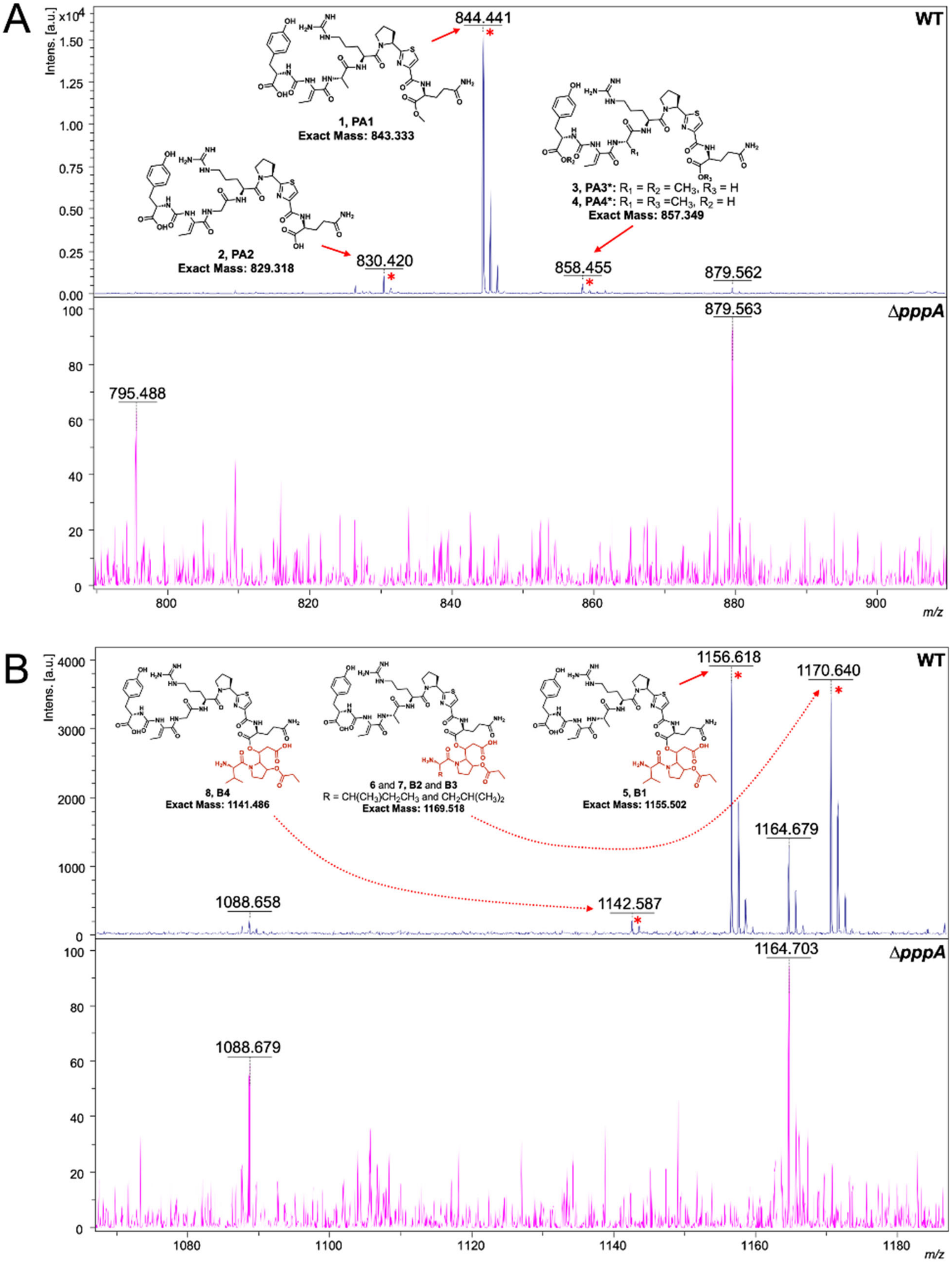
A
WT
ΔpppA
Intens. [a.u.]
x10^4
1, PA1
Exact Mass: 843.333
2, PA2
Exact Mass: 829.318
3, PA3*: R1 = R2 = CH3, R3 = H
4, PA4*: R1 = R3 =CH3, R2 = H
Exact Mass: 857.349
844.441
830.420
858.455
879.562
879.563
795.488
m/z
B
WT
ΔpppA
Intens. [a.u.]
8, B4
Exact Mass: 1141.486
6 and 7, B2 and B3
R = CH(CH3)CH2CH3 and CH2CH(CH3)2
Exact Mass: 1169.518
5, B1
Exact Mass: 1155.502
1156.618
1170.640
1164.679
1142.587
1088.658
1164.703
1088.679
m/z

**Figure S3. MALDI-ToF MS analysis of wild type and ∆*pppA* mutants.** (**A**) The spectrum is zoomed in the range from *m/z* 790 to *m/z* 910. Molecular features of four PA congeners are indicated with red asterisks, that is, *m/z* 830.4 (PA2 $[M+H]^+$), *m/z* 844.4 (PA1 $[M+H]^+$), and *m/z* 858.4 (PA3 or PA4 $[M+H]^+$). (**B**) The spectrum is zoomed in the range from *m/z* 1068 to *m/z* 1183. Molecular features of four PB congeners are indicated with red asterisks, that is, *m/z* 1142.6 (PB4 Val, $[M+H]^+$), *m/z* 1156.6 (PB1 Val, $[M+H]^+$), *m/z* 1170.6 (PB2 or PB3 Ile or Leu, $[M+H]^+$).

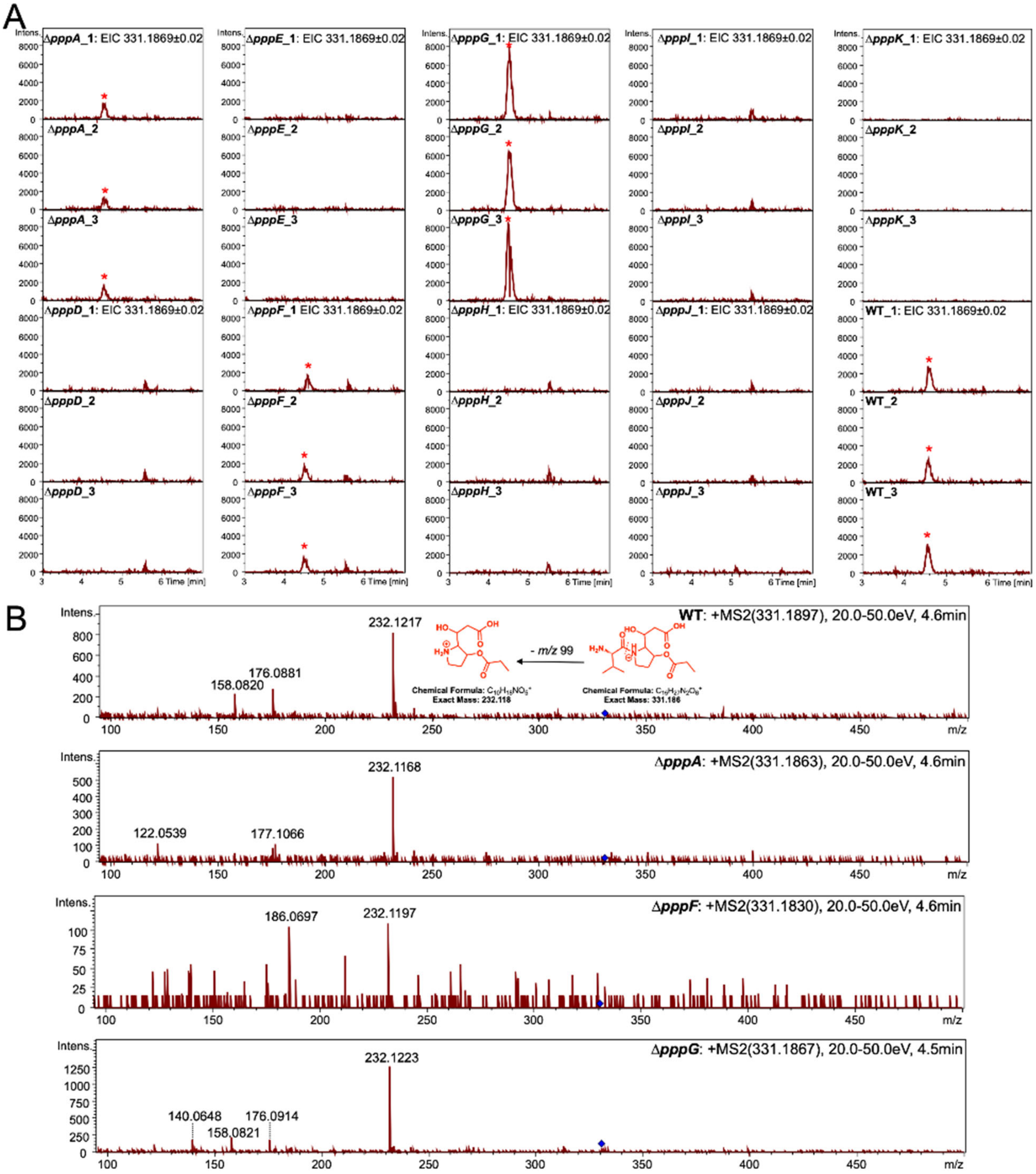

**Figure S4. Comparison of PC1 production and MS/MS fragmentation between wild type and mutants in triplicates.** UPLC-QToF-MS analyses were performed for both WT and mutants in triplicates. (**A**) Extracted Ion Chromatograms (EIC) of singly charged PC1 (Val, $[M+H]^+$). The same mass filter (the expected *m/z* ratio ± *m/z* 0.02) was applied to all samples. PC1 peaks are highlighted with a red asterisk. (**B**) Fragmentation pattern of PC1 produced from WT compared to Δ*pppA/F/G* mutants. The proposed structures of parent and fragment peaks are shown. The upper right numbers from left to right are the *m/z* of the fragmentated compound, the collision energy, and the retention time. The blue diamond pinpoints the parent peak.

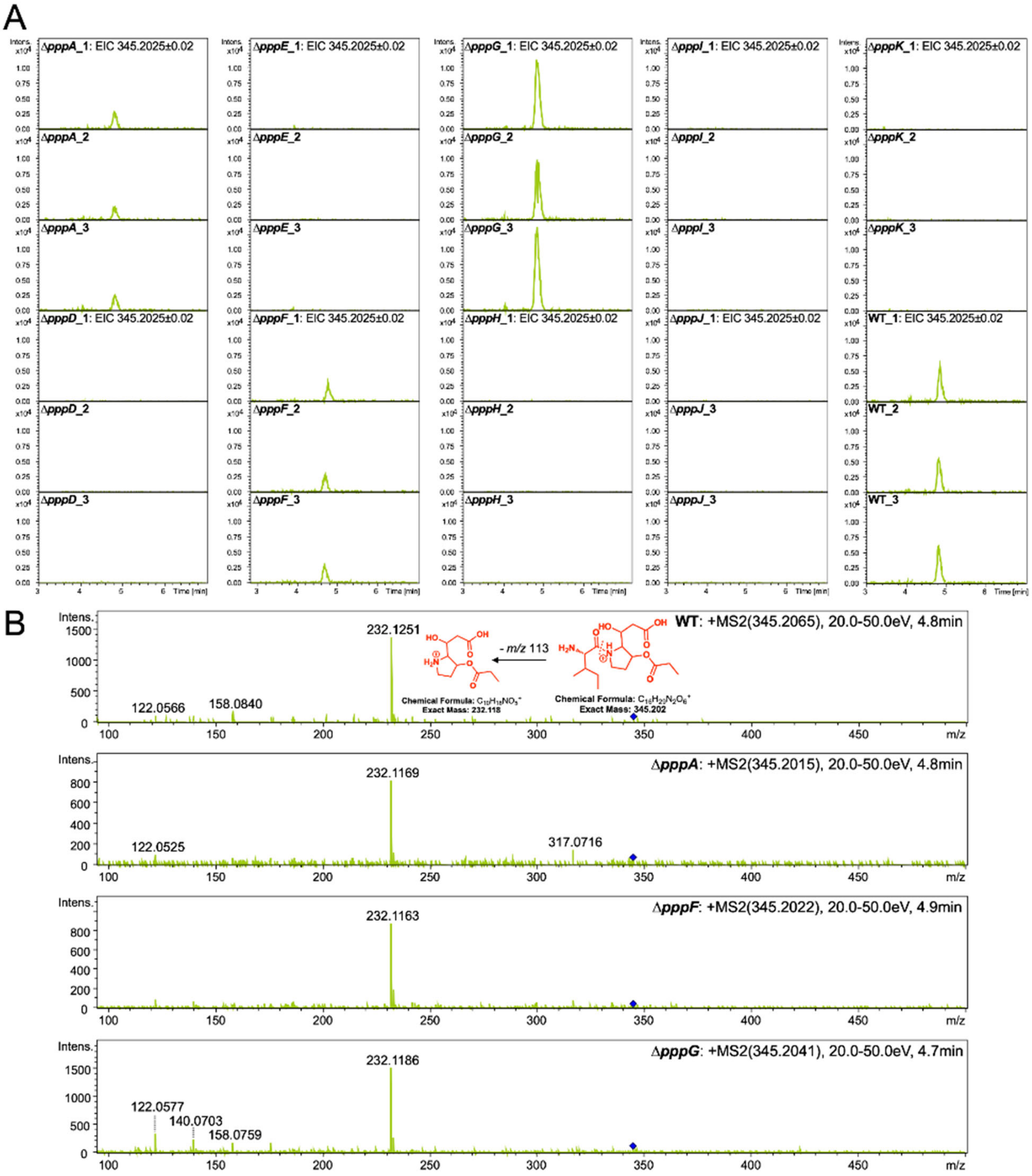

**Figure S5. Comparison of PC2 or PC3 production and MS/MS fragmentation between wild type and mutants in triplicates.** UPLC-QToF-MS analyses were performed for both WT and mutants in triplicates. (**A**) Extracted Ion Chromatograms (EIC) of singly charged PC2 or PC3 (Ile or Leu, $[M+H]^+$). The same mass filter (the expected *m/z* ± 0.02) was applied to all samples. (**B**) The fragmentation pattern of PC2 or PC3 produced from WT compared to Δ*pppA/F/G* mutants. The proposed structures of parent and fragment peaks are shown. The blue diamond pinpoints the parent peak.

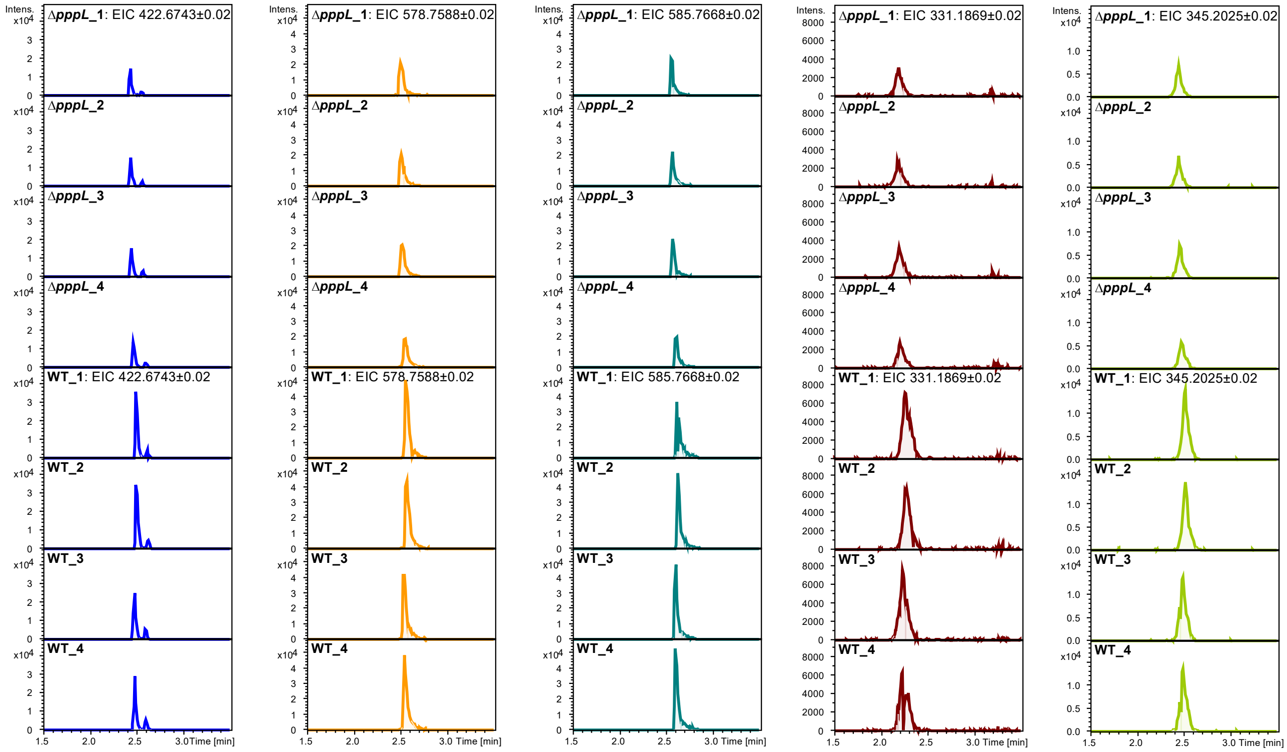

**Figure S6. Comparison of PA, PB, and PC production between wild type and Δ*pppL* mutants.** UPLC-QToF-MS analyses were performed for both WT and Δ*pppL* in quadruplicates. Extracted Ion Chromatograms (EIC) of (from left to right) doubly charged PA1 (*m/z* 442.6743, $[M+2H]^{2+}$), doubly charged PB1 (*m/z* 578.7588, Val, $[M+2H]^{2+}$) and PB2 or PB3 (*m/z* 585.7668, Ile or Leu, $[M+2H]^{2+}$), and singly charged PC1 (*m/z* 331.1869, Val, $[M+H]^+$) and PC2 or PC3 (*m/z*

345.2025, Ile or Leu, [M+H]⁺). The same mass filter (the expected *m/z* ± 0.02) was applied to all samples.

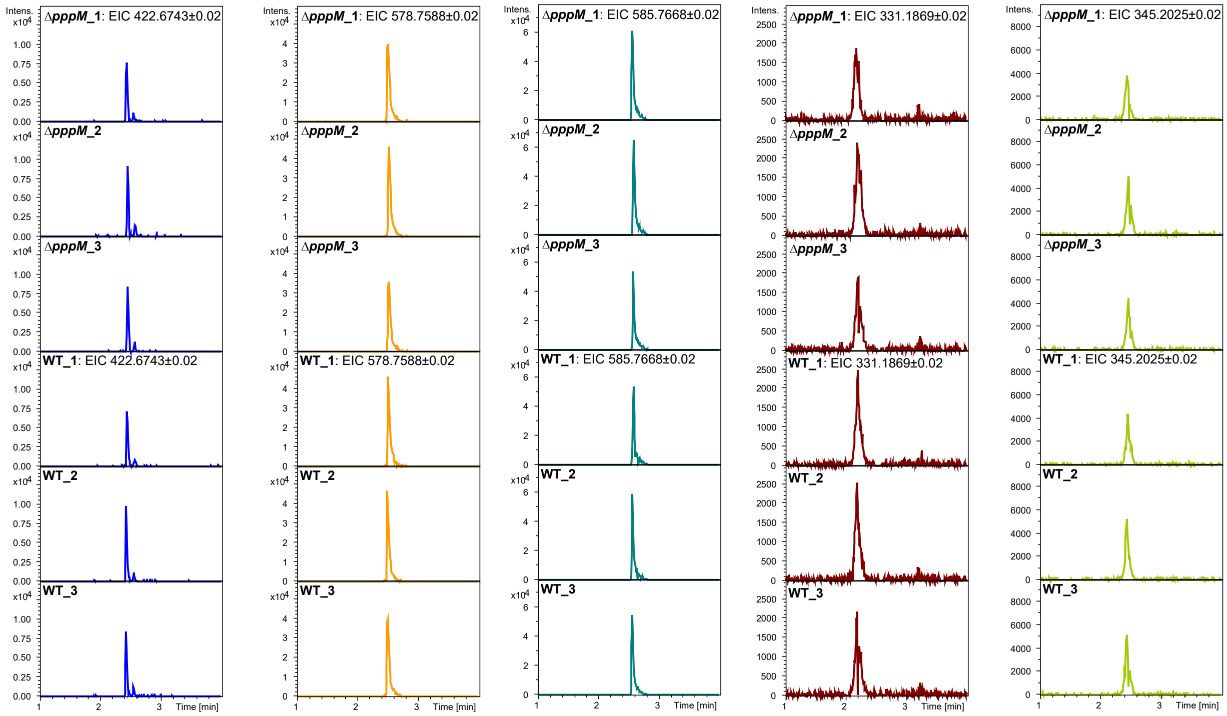

**Figure S7. Comparison of PA, PB, and PC production between wild type and Δ*pppM* mutants.** UPLC-QToF-MS analyses were performed for both WT and Δ*pppM* in triplicates. Extracted Ion Chromatograms (EIC) of (from left to right) doubly charged PA1 (*m/z* 442.6743, $[M+2H]^{2+}$), doubly charged PB1 (*m/z* 578.7588, Val, $[M+2H]^{2+}$) and PB2 or PB3 (*m/z* 585.7668, Ile or Leu, $[M+2H]^{2+}$), and singly charged PC1 (*m/z* 331.1869, Val, $[M+H]^{+}$) and PC2 or PC3 (*m/z* 345.2025, Ile or Leu, $[M+H]^{+}$). The same mass filter (the expected *m/z* ± 0.02) was applied to all samples.

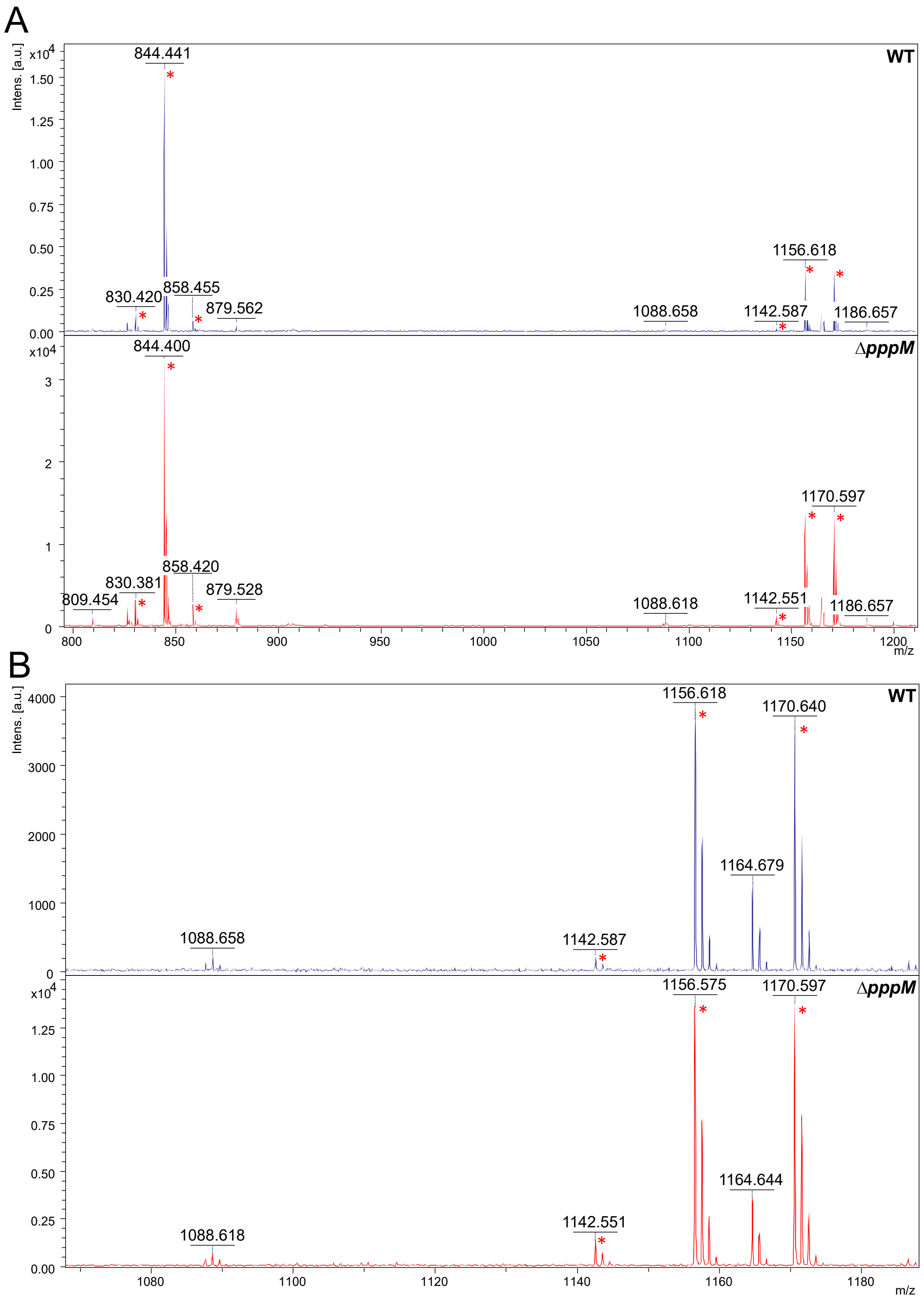
A
Intens. [a.u.]
x10^4
WT
844.441
830.420
858.455
879.562
1088.658
1142.587
1156.618
1186.657
ΔpppM
844.400
809.454
830.381
858.420
879.528
1088.618
1142.551
1170.597
1186.657
m/z
B
WT
1088.658
1142.587
1156.618
1164.679
1170.640
ΔpppM
1088.618
1142.551
1156.575
1164.644
1170.597

**Figure S8. MALDI-ToF MS analyses of wild type and ∆*pppM* mutant.** MALDI-ToF MS analyses were performed for both WT and ∆*pppM* mutant. (**A**) The molecular features for both PA and PB are indicated with red asterisks. *m/z* 830.4, PA2 ($[M+H]^+$); *m/z* 844.4, PA1 ($[M+H]^+$); and *m/z* 858.4, PA3 or PA4 ($[M+H]^+$). *m/z* 1142.6, PB4 (Val, $[M+H]^+$); *m/z* 1156.6, PB1 (Val, $[M+H]^+$); *m/z* 1170.6, PB2 or PB3 (Ile or Leu, $[M+H]^+$). (**B**) The spectrum is zoomed in the range from *m/z* 1068 to *m/z* 1183 to highlight that no modified PB was observed in the ∆*pppM* mutant.

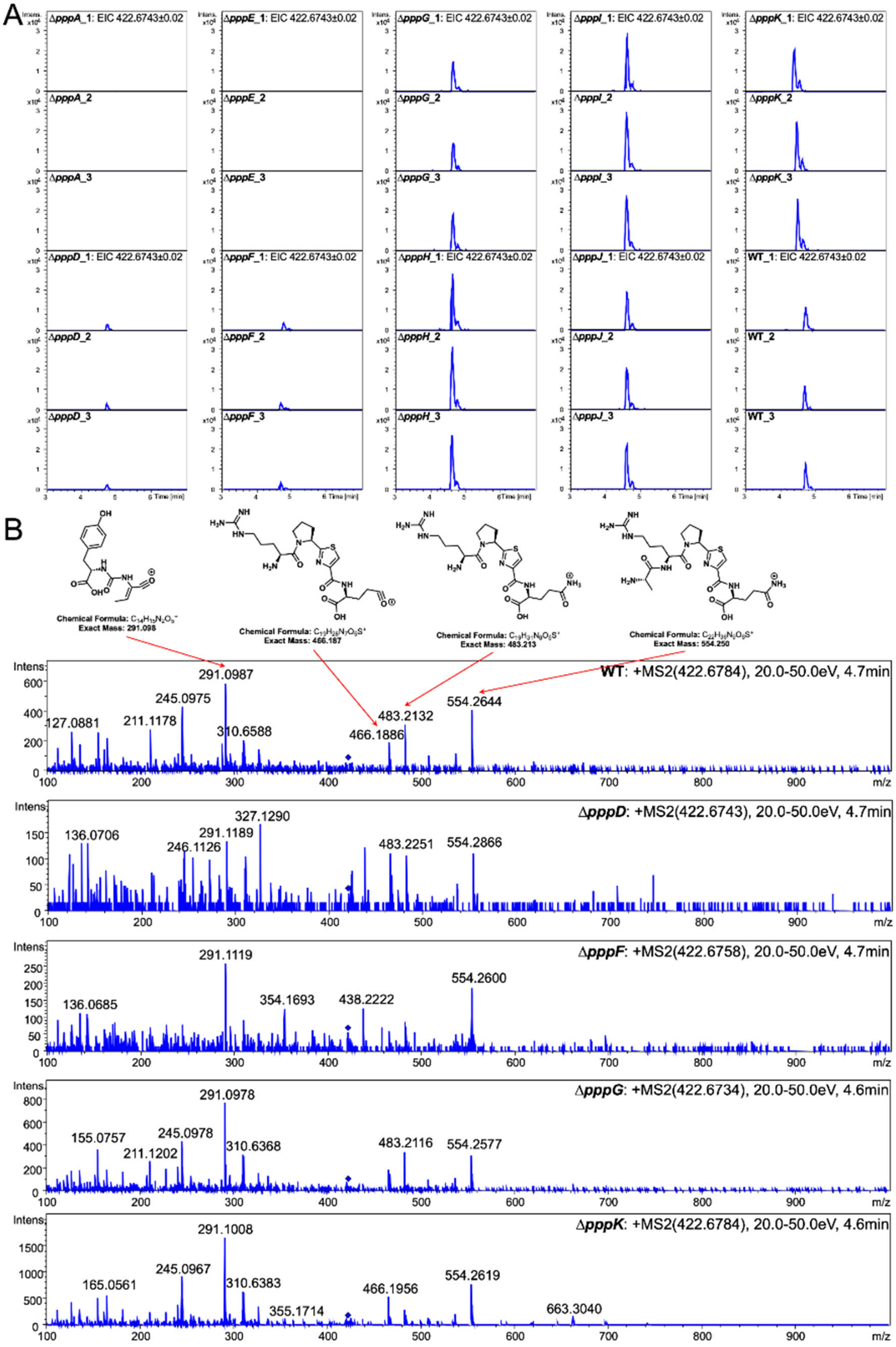

A
ΔpppA_1: EIC 422.6743±0.02
ΔpppE_1: EIC 422.6743±0.02
ΔpppG_1: EIC 422.6743±0.02
ΔpppI_1: EIC 422.6743±0.02
ΔpppK_1: EIC 422.6743±0.02
ΔpppD_1: EIC 422.6743±0.02
ΔpppF_1: EIC 422.6743±0.02
ΔpppH_1: EIC 422.6743±0.02
ΔpppJ_1: EIC 422.6743±0.02
WT_1: EIC 422.6743±0.02
Time [min]
B
Exact Mass: 291.098
Exact Mass: 466.187
Exact Mass: 483.213
Exact Mass: 554.250
WT: +MS2(422.6784), 20.0-50.0eV, 4.7min
ΔpppD: +MS2(422.6743), 20.0-50.0eV, 4.7min
ΔpppF: +MS2(422.6758), 20.0-50.0eV, 4.7min
ΔpppG: +MS2(422.6734), 20.0-50.0eV, 4.6min
ΔpppK: +MS2(422.6784), 20.0-50.0eV, 4.6min
Intens.
m/z

**Figure S9. Comparison of PA1 production and MS/MS fragmentation between wild type and mutants in triplicates.** UPLC-QToF-MS analyses were performed for both WT and mutants in triplicates. (**A**) Extracted Ion Chromatograms (EIC) of doubly charged PA1 ($[M+2H]^{2+}$). The same mass filter (the expected *m/z* ± 0.02) was applied to all samples. (**B**) The fragmentation pattern of PA1 produced from WT is compared to those of Δ*pppD/F/G/K* mutants. Proposed structures of major peaks are shown. The upper right numbers from left to right are the *m/z* of the fragmentated compound, the collision energy, and the retention time. The blue diamond pinpoints the parent peak.

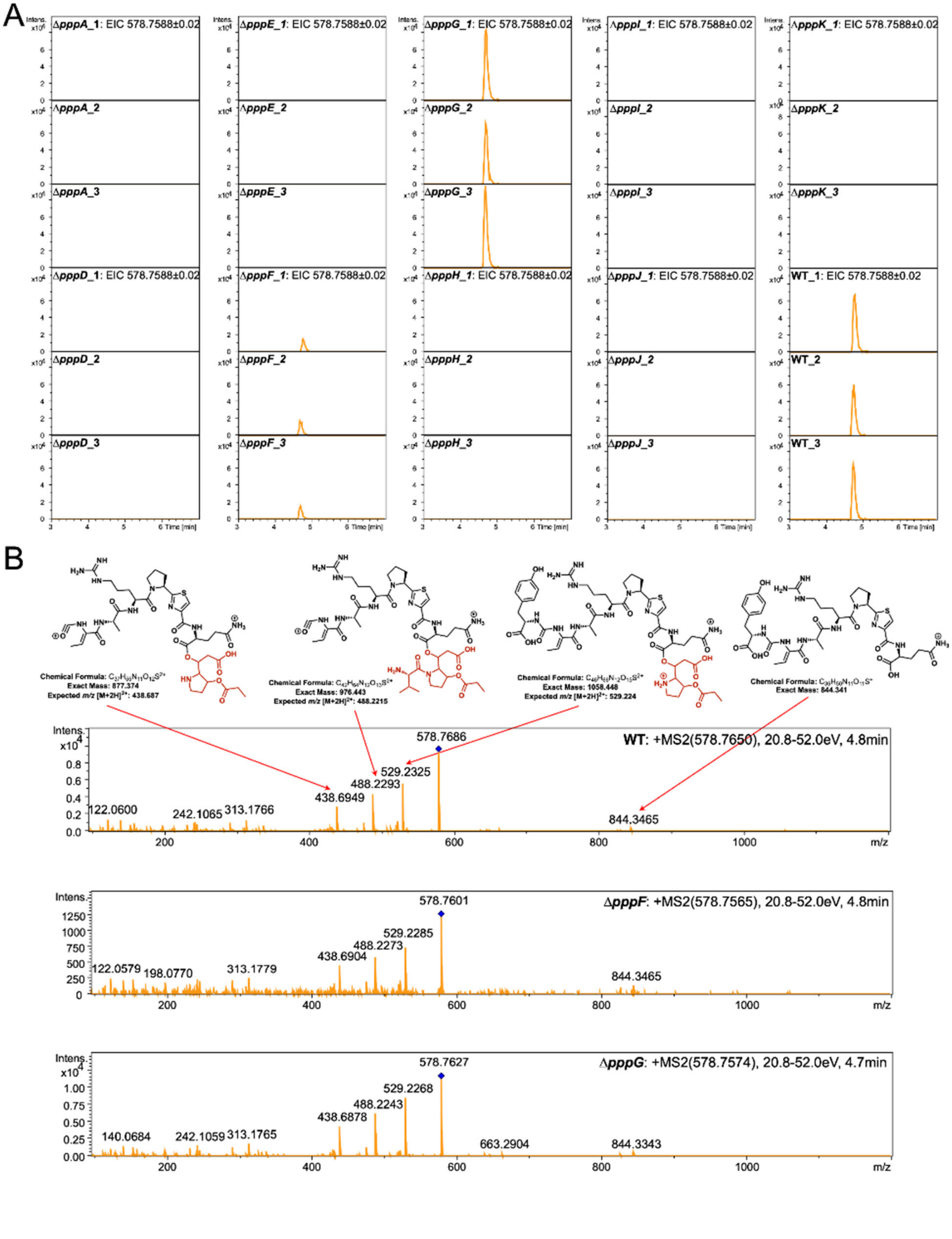
A
ΔpppA_1: EIC 578.7588±0.02
ΔpppE_1: EIC 578.7588±0.02
ΔpppG_1: EIC 578.7588±0.02
ΔpppI_1: EIC 578.7588±0.02
ΔpppK_1: EIC 578.7588±0.02
ΔpppA_2
ΔpppE_2
ΔpppG_2
ΔpppI_2
ΔpppK_2
ΔpppA_3
ΔpppE_3
ΔpppG_3
ΔpppI_3
ΔpppK_3
ΔpppD_1: EIC 578.7588±0.02
ΔpppF_1: EIC 578.7588±0.02
ΔpppH_1: EIC 578.7588±0.02
ΔpppJ_1: EIC 578.7588±0.02
WT_1: EIC 578.7588±0.02
ΔpppD_2
ΔpppF_2
ΔpppH_2
ΔpppJ_2
WT_2
ΔpppD_3
ΔpppF_3
ΔpppH_3
ΔpppJ_3
WT_3
Time [min]
B
Exact Mass: 877.374
Expected m/z [M+2H]2+: 438.687
Exact Mass: 976.443
Expected m/z [M+2H]2+: 488.2215
Exact Mass: 1058.448
Expected m/z [M+2H]2+: 529.224
Exact Mass: 844.341
WT: +MS2(578.7650), 20.8-52.0eV, 4.8min
578.7686
529.2325
488.2293
438.6949
122.0600
242.1065
313.1766
844.3465
ΔpppF: +MS2(578.7565), 20.8-52.0eV, 4.8min
578.7601
529.2285
488.2273
438.6904
122.0579
198.0770
313.1779
844.3465
ΔpppG: +MS2(578.7574), 20.8-52.0eV, 4.7min
578.7627
529.2268
488.2243
438.6878
140.0684
242.1059
313.1765
663.2904
844.3343
Intens.
m/z

**Figure S10. Comparison of PB1 production and MS/MS fragmentation between wild type and mutants in triplicates.** UPLC-QToF-MS analyses were performed for both WT and mutants in triplicates. (**A**) Extracted Ion Chromatograms (EIC) of doubly charged PB1 (Val, $[M+2H]^{2+}$). The same mass filter (the expected *m/z* ± 0.02) was applied to all samples. (**B**) The fragmentation pattern of PB1 produced from WT compared to those of Δ*pppF/G* mutants. Proposed structures of major peaks are shown. The upper right numbers from left to right are the *m/z* of the fragmentated compound, the collision energy, and the retention time. The blue diamond pinpoints the parent peak.

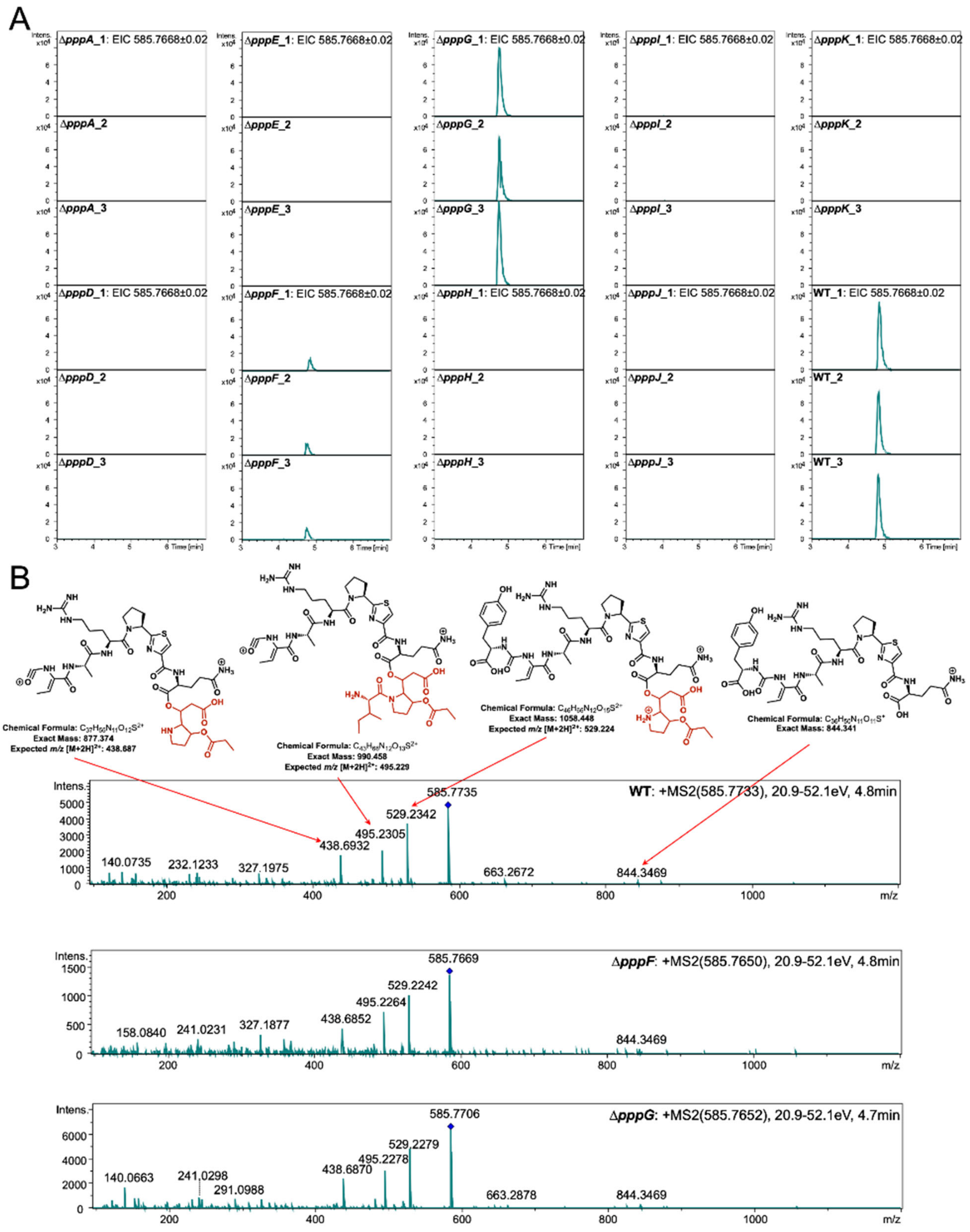
A
ΔpppA_1: EIC 585.7668±0.02
ΔpppE_1: EIC 585.7668±0.02
ΔpppG_1: EIC 585.7668±0.02
ΔpppI_1: EIC 585.7668±0.02
ΔpppK_1: EIC 585.7668±0.02
ΔpppA_2
ΔpppE_2
ΔpppG_2
ΔpppI_2
ΔpppK_2
ΔpppA_3
ΔpppE_3
ΔpppG_3
ΔpppI_3
ΔpppK_3
ΔpppD_1: EIC 585.7668±0.02
ΔpppF_1: EIC 585.7668±0.02
ΔpppH_1: EIC 585.7668±0.02
ΔpppJ_1: EIC 585.7668±0.02
WT_1: EIC 585.7668±0.02
ΔpppD_2
ΔpppF_2
ΔpppH_2
ΔpppJ_2
WT_2
ΔpppD_3
ΔpppF_3
ΔpppH_3
ΔpppJ_3
WT_3
Time [min]
B
Chemical Formula: C37H62N11O12S2+
Exact Mass: 877.374
Expected m/z [M+2H]2+: 438.687
Chemical Formula: C43H68N12O13S2+
Exact Mass: 990.458
Expected m/z [M+2H]2+: 495.229
Chemical Formula: C46H66N12O15S2+
Exact Mass: 1058.448
Expected m/z [M+2H]2+: 529.224
Chemical Formula: C36H52N11O11S+
Exact Mass: 844.341
WT: +MS2(585.7733), 20.9-52.1eV, 4.8min
585.7735
529.2342
495.2305
438.6932
140.0735
232.1233
327.1975
663.2672
844.3469
ΔpppF: +MS2(585.7650), 20.9-52.1eV, 4.8min
585.7669
529.2242
495.2264
438.6852
158.0840
241.0231
327.1877
844.3469
ΔpppG: +MS2(585.7652), 20.9-52.1eV, 4.7min
585.7706
529.2279
495.2278
438.6870
140.0663
241.0298
291.0988
663.2878
844.3469
Intens.
m/z

**Figure S11. Comparison of PB2 or PB3 production and MS/MS fragmentation between wild type and mutants in triplicates.** UPLC-QToF-MS analyses were performed for both WT and mutants in triplicates. (**A**) Extracted Ion Chromatograms (EIC) of doubly charged PB2 or PB3 (Ile or Leu, $[M+2H]^{2+}$). The same mass filter (the expected *m/z* ± 0.02) was applied to all samples. (**B**) The fragmentation pattern of PB2 or PB3 produced from WT compared to Δ*pppF/G* mutants. Proposed structures of major peaks are shown. The structure of PB2 is shown as the representative of peak at *m/z* 495.23. The upper right numbers from left to right are the *m/z* of the fragmentated compound, the collision energy, and the retention time. The blue diamond pinpoints the parent peak.

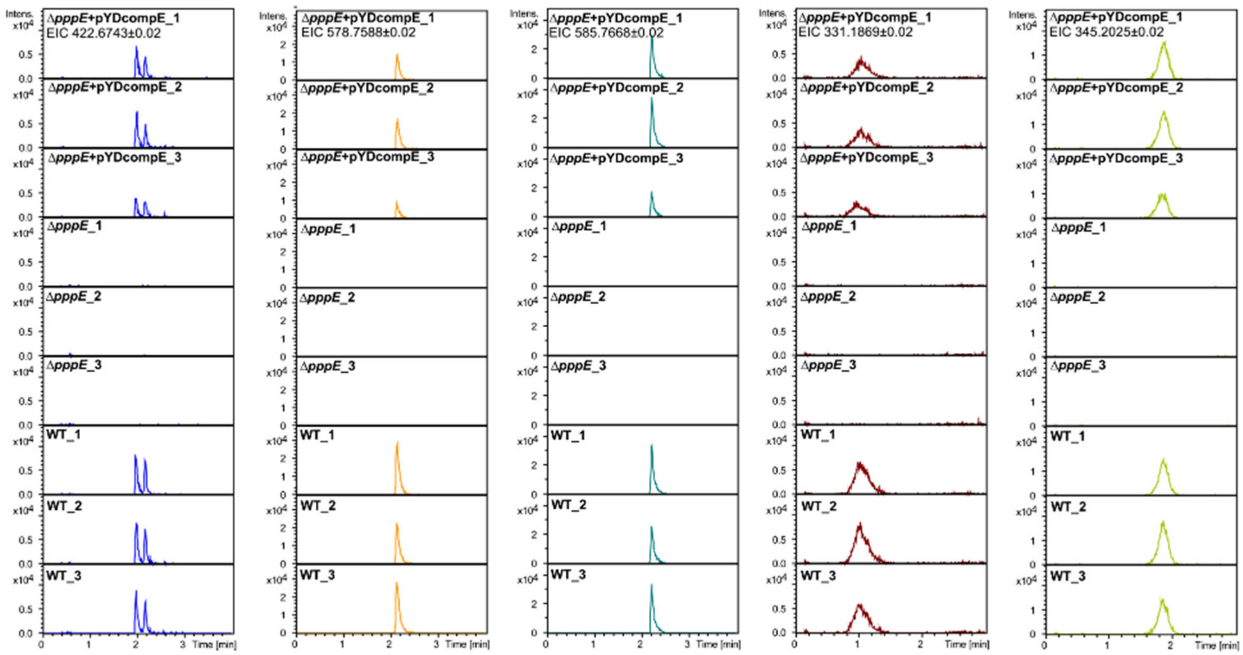

**Figure S12. Comparison of PA, PB, and PC production between Δ*pppE* mutant, wild type and genetically complemented Δ*pppE* mutant.** UPLC-QToF-MS analyses were performed for wild type (bottom), Δ*pppE* mutant (middle), and complemented Δ*pppE* mutant (top) in triplicates. Extracted Ion Chromatograms (EIC) of (from left to right) doubly charged PA1 (*m/z* 442.6743, $[M+2H]^{2+}$), doubly charged PB1 (*m/z* 578.7588, Val, $[M+2H]^{2+}$) and PB2 or PB3 (*m/z* 585.7668, Ile or Leu, $[M+2H]^{2+}$), and singly charged PC1 ($m/z$ 331.1869, Val, $[M+H]^+$) and PC2 or PC3 ($m/z$ 345.2025, Ile or Leu, $[M+H]^+$). The same mass filter (the expected $m/z$ ± 0.02) was applied to all samples.

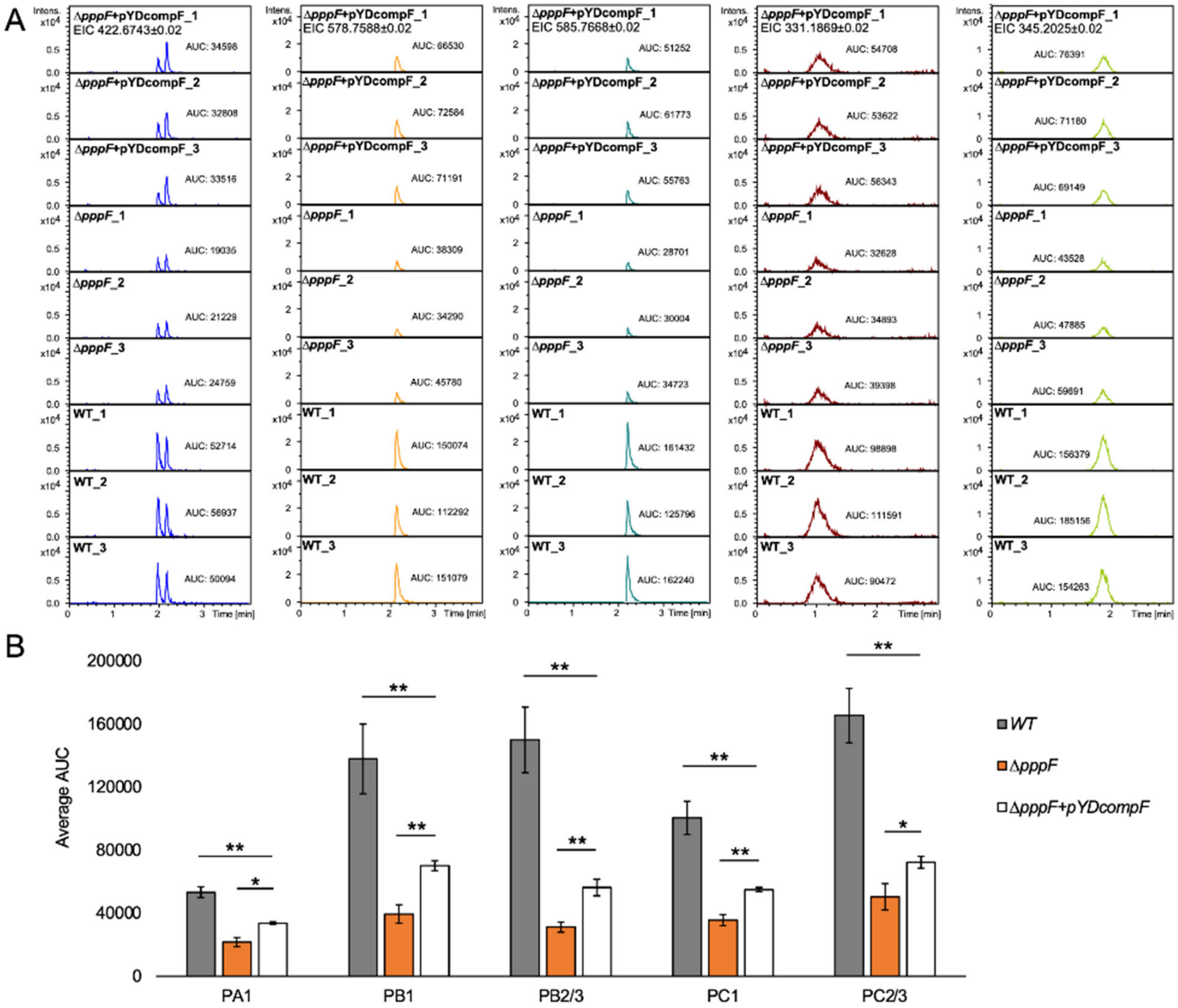

**Figure S13. Comparison of PA, PB, and PC production between Δ*pppF* mutant, wild type and genetically complemented Δ*pppF* mutant.** (**A**) UPLC-QToF-MS analyses were performed for the complemented Δ*pppF* mutant (top), wild type (bottom), and the Δ*pppF* mutant (middle) in triplicates. Extracted Ion Chromatograms (EIC) of (from left to right) doubly charged PA1 ($m/z$ 442.6743, $[M+2H]^{2+}$), doubly charged PB1 ($m/z$ 578.7588, Val, $[M+2H]^{2+}$) and PB2 or PB3 ($m/z$ 585.7668, Ile or Leu, $[M+2H]^{2+}$), and singly charged PC1 ($m/z$ 331.1869, Val, $[M+H]^+$) and PC2 or PC3 ($m/z$ 345.2025, Ile or Leu, $[M+H]^+$). The same mass filter (the expected $m/z$ ± 0.02) was applied to all samples. AUC, area under curve. (**B**) The average AUC of each pseudovibriamide from the wild type, the Δ*pppF* mutant, and the genetically complemented Δ*pppF* mutant was used to assess the relative amount of each pseudovibriamide produced. Two-tail *P*-values from t-Test were used to determine statistical significance; *, *P*-value ≤ 0.05; and **, *P*-value ≤ 0.01.

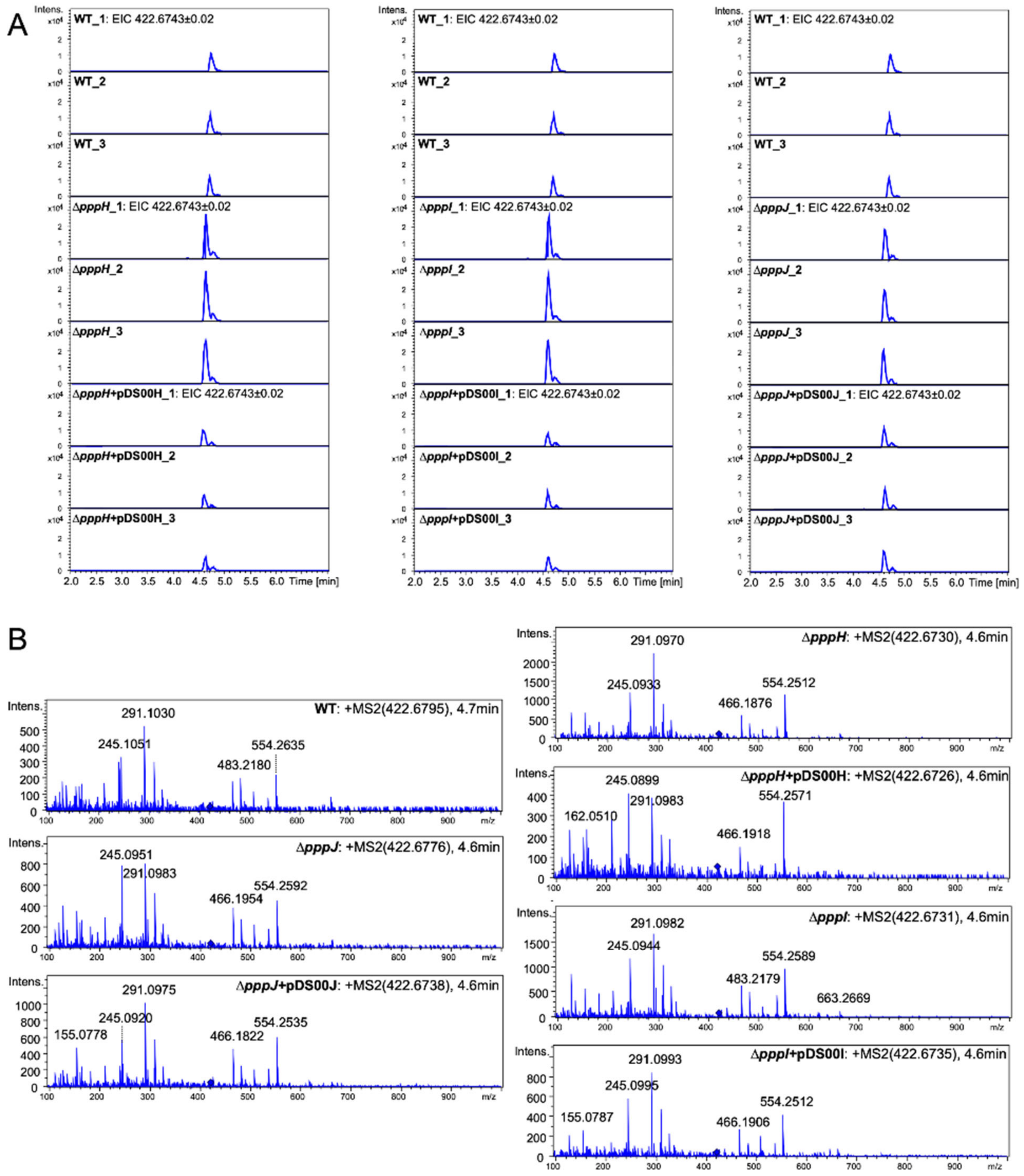

**Figure S14. Comparison of PA1 production and MS/MS fragmentation between wild type, ∆*pppH,* ∆*pppI,* ∆*pppJ* mutants, and genetically complemented strains.** UPLC-QToF-MS analyses were performed in triplicates. (**A**) Extracted Ion Chromatograms (EIC) of doubly charged PA1 ($[M+2H]^{2+}$). The same mass filter (the expected *m/z* ± 0.02) was applied to all samples. (**B**) Fragmentation pattern of PA1 produced from WT compared to ∆*pppH,* ∆*pppI,* and ∆*pppJ* mutants and complemented ∆*pppH,* ∆*pppI,* and ∆*pppJ* mutants. Expected fragmentation peaks of PA1 are *m/z* 554.25, *m/z* 483.21, *m/z* 466.19, and *m/z* 291.10 (**Fig. S9**). The upper right numbers from left to right are the *m/z* ratio of the fragmentated compound, and the retention time. The blue diamond pinpoints the parent peak.

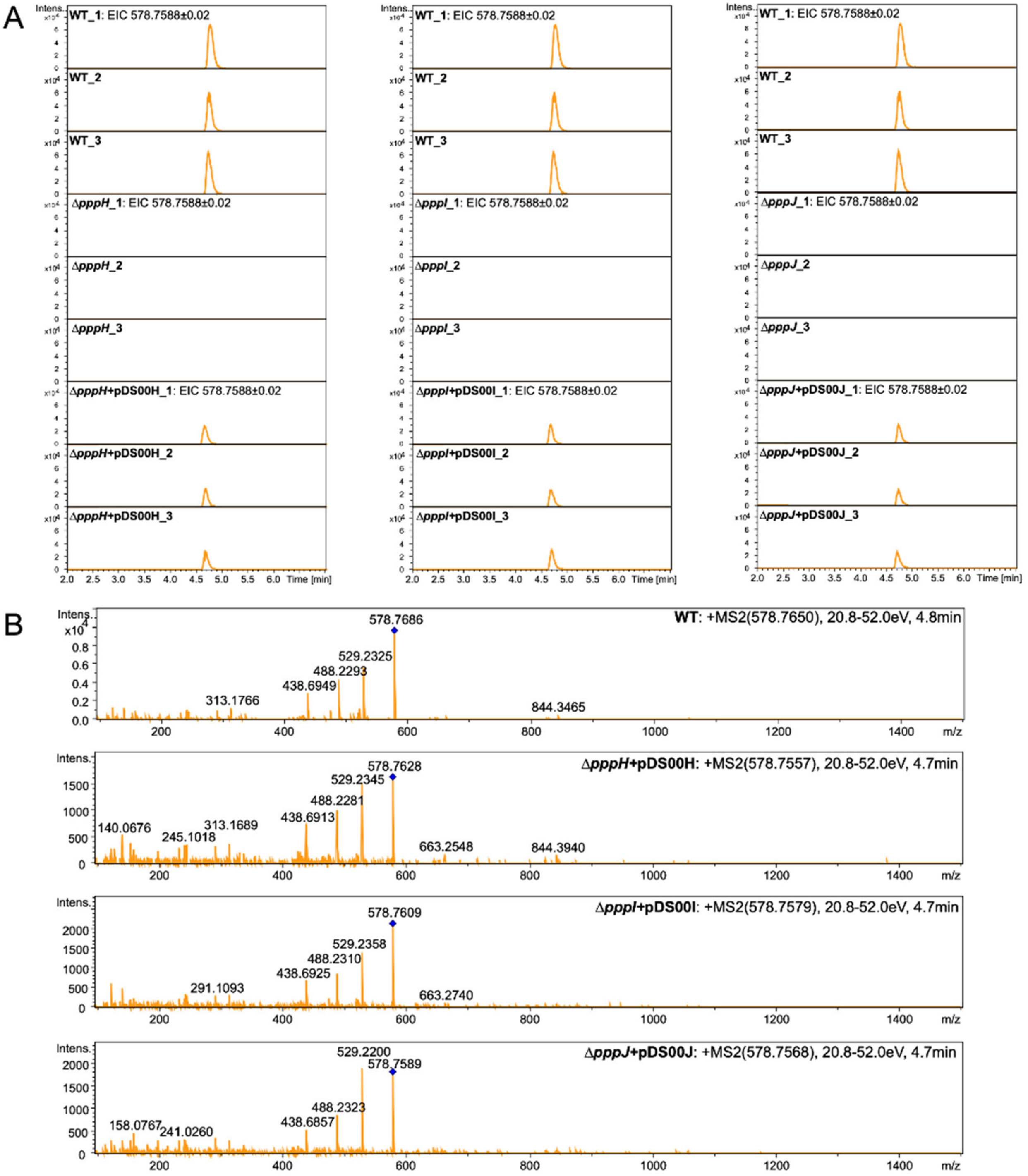

**Figure S15. Comparison of PB1 production and MS/MS fragmentation between wild type, Δ*pppH*, Δ*pppI*, Δ*pppJ* mutants, and genetically complemented strains.** UPLC-QToF-MS analyses were performed in triplicates. (**A**) Extracted Ion Chromatograms (EIC) of doubly charged PB1 (Val, $[M+2H]^{2+}$). The same mass filter (the expected *m/z* ± 0.02) was applied to all samples.

(**B**) The fragmentation pattern of PB1 produced from WT compared to those of complemented Δ*pppH,* Δ*pppI,* and Δ*pppJ* mutants. Expected fragmentation peaks of PB1 are *m/z* 844.34, *m/z* 529.22, *m/z* 488.22, and *m/z* 438.69 (**Fig. S10**). The upper right numbers from left to right are the *m/z* of the fragmentated compound, the collision energy, and the retention time. The blue diamond pinpoints the parent peak.

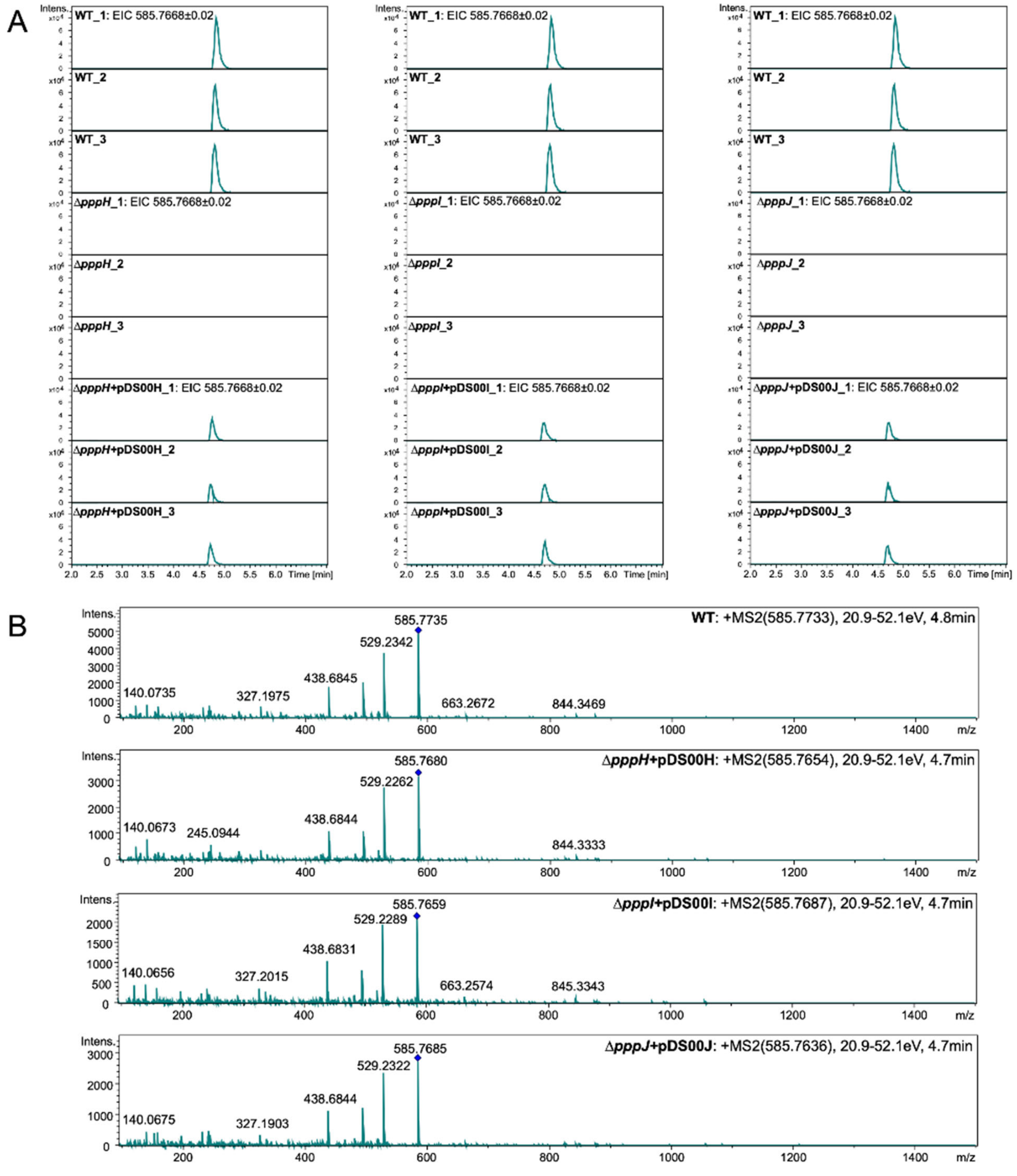

**Figure S16. Comparison of PB2 or PB3 production and MS/MS fragmentation between wild type, Δ*pppH,* Δ*pppI,* Δ*pppJ* mutants, and genetically complemented strains.** UPLC-QToF-MS analyses were performed in triplicates. (**A**) Extracted Ion Chromatograms (EIC) of doubly charged PB2 or PB3 (Ile or Leu, $[M+2H]^{2+}$). The same mass filter (the expected *m/z* ± 0.02) was applied to all samples. (**B**) Fragmentation pattern of PB2 or PB3 produced from WT compared to those of complemented Δ*pppH,* Δ*pppI,* and Δ*pppJ* mutants. Expected fragmentation peaks of PB2 or PB3 are *m/z* 844.34, *m/z* 529.22, *m/z* 495.23, and *m/z* 438.69 (**Fig. S11**).The upper right numbers from left to right are the *m/z* of the fragmentated compound, the collision energy, and the retention time. The blue diamond pinpoints the parent peak.

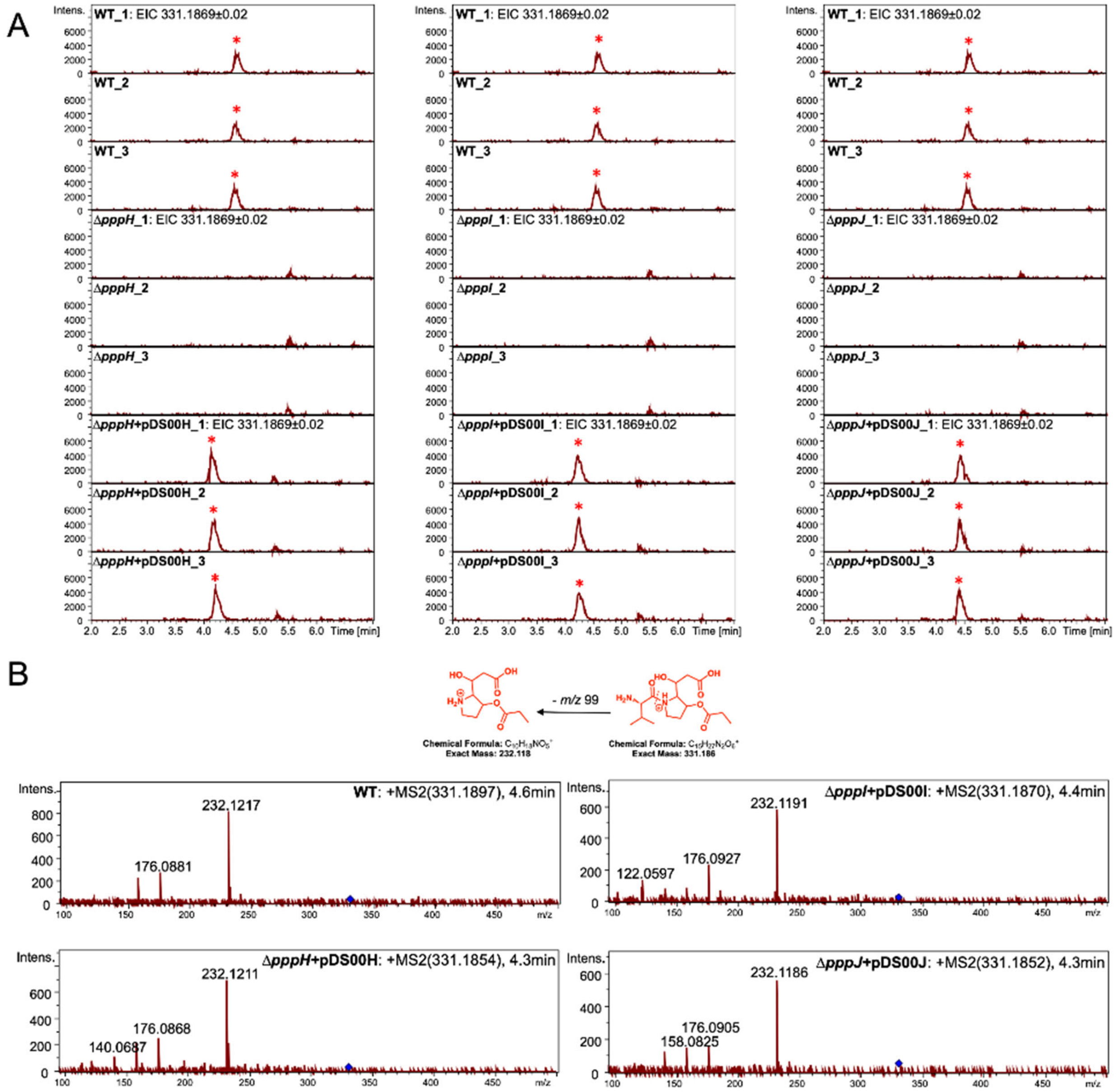

**Figure S17. Comparison of PC1 production and MS/MS fragmentation between wild type, ∆*pppH,* ∆*pppI,* ∆*pppJ* mutants, and genetically complemented strains.** UPLC-QToF-MS analyses were performed in triplicates. (**A**) Extracted Ion Chromatograms (EIC) of singly charged PC1 (Val, $[M+H]^+$). The same mass filter (the expected *m/z* ± 0.02) was applied to all samples. PC1 peaks are highlighted with a red asterisk. (**B**) Fragmentation pattern of PC1 produced from WT compared to those of ∆*pppH,* ∆*pppI,* and ∆*pppJ* mutants and complemented ∆*pppH,* ∆*pppI,* and ∆*pppJ* mutants*.* The proposed structures of parent and fragment peaks are shown. The upper right numbers from left to right are the *m/z* of the fragmentated compound and the retention time. The blue diamond pinpoints the parent peak.

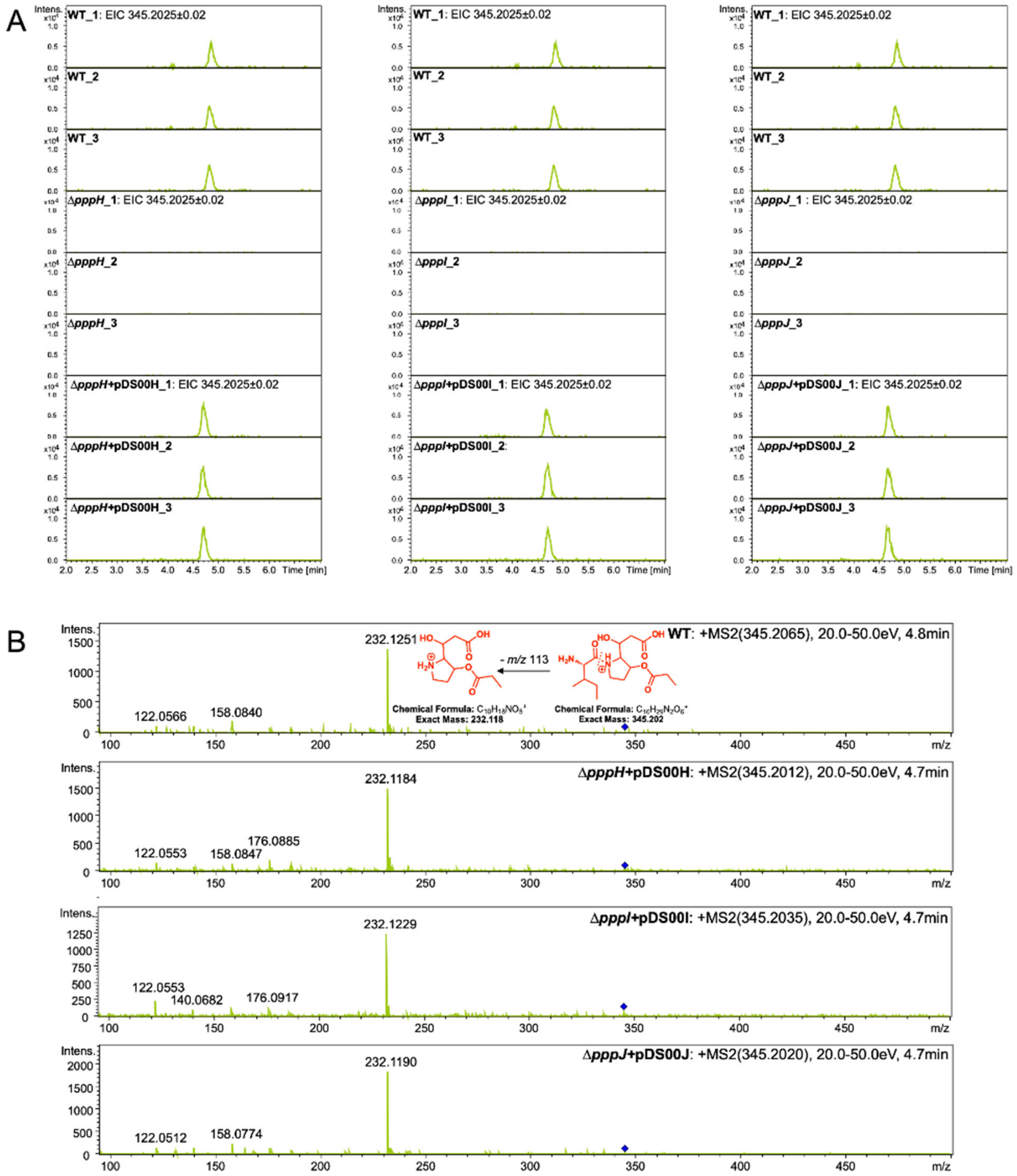

**Figure S18. Comparison of PC2 or PC3 production and MS/MS fragmentation between wild type, Δ*pppH,* Δ*pppI,* Δ*pppJ* mutants, and genetically complemented strains.** UPLC-QToF-MS analyses were performed in triplicates. (**A**) Extracted Ion Chromatograms (EIC) of singly charged PC2 or PC3 (Leu or Ile, $[M+H]^+$). The same mass filter (the expected *m/z* ± 0.02) was applied to all samples. (**B**) Fragmentation pattern of PC2 or PC3 produced from WT compared to those of complemented Δ*pppH,* Δ*pppI,* and Δ*pppJ* mutants. The proposed structures of parent and fragment peaks are shown. The upper right numbers from left to right are the *m/z* of the fragmentated compound and the retention time. The blue diamond pinpoints the parent peak.

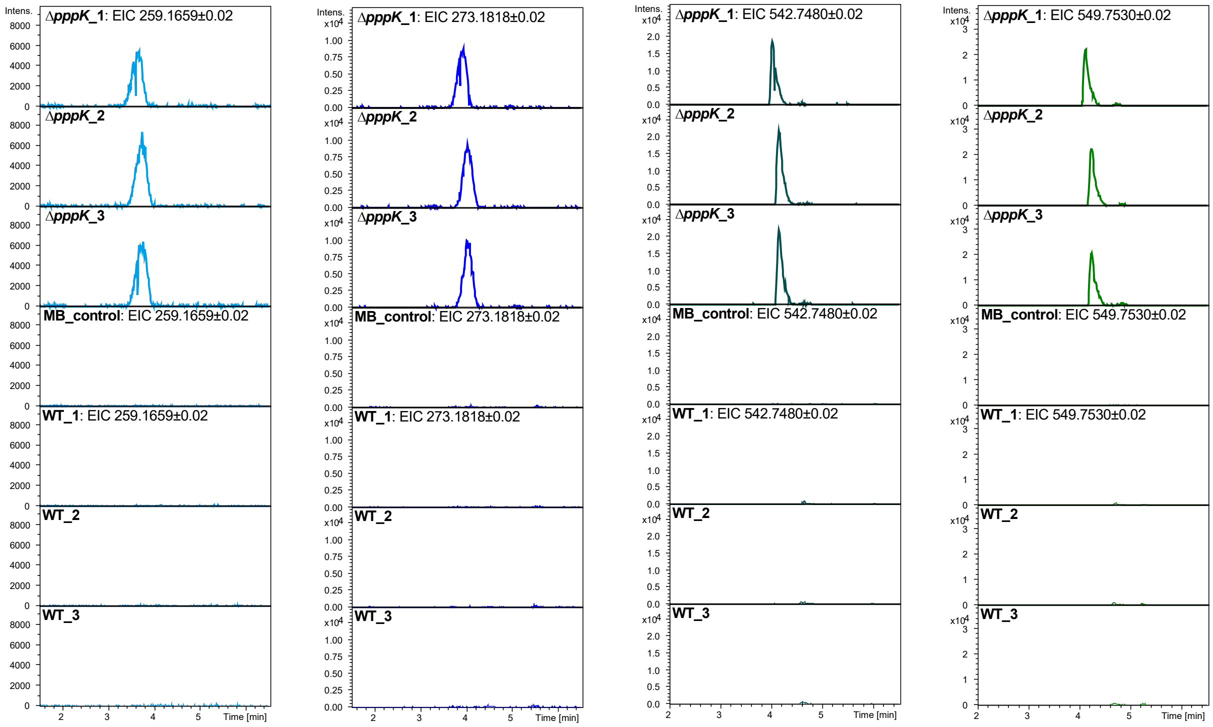

**Figure S19. Comparison of depropionylated PBs and PCs production between wild type and Δ*pppK*.** UPLC-QToF-MS analyses were performed in triplicates. Extracted Ion Chromatograms (EIC) of (from left to right) depropionylated PCs (Val, $[M+H]^+$ and Ile or Leu, $[M+H]^+$), and depropionylated PBs (Val, $[M+2H]^{2+}$ and Leu or Ile, $[M+2H]^{2+}$). The same mass filter (the expected *m/z* ± 0.02) was applied to all samples.

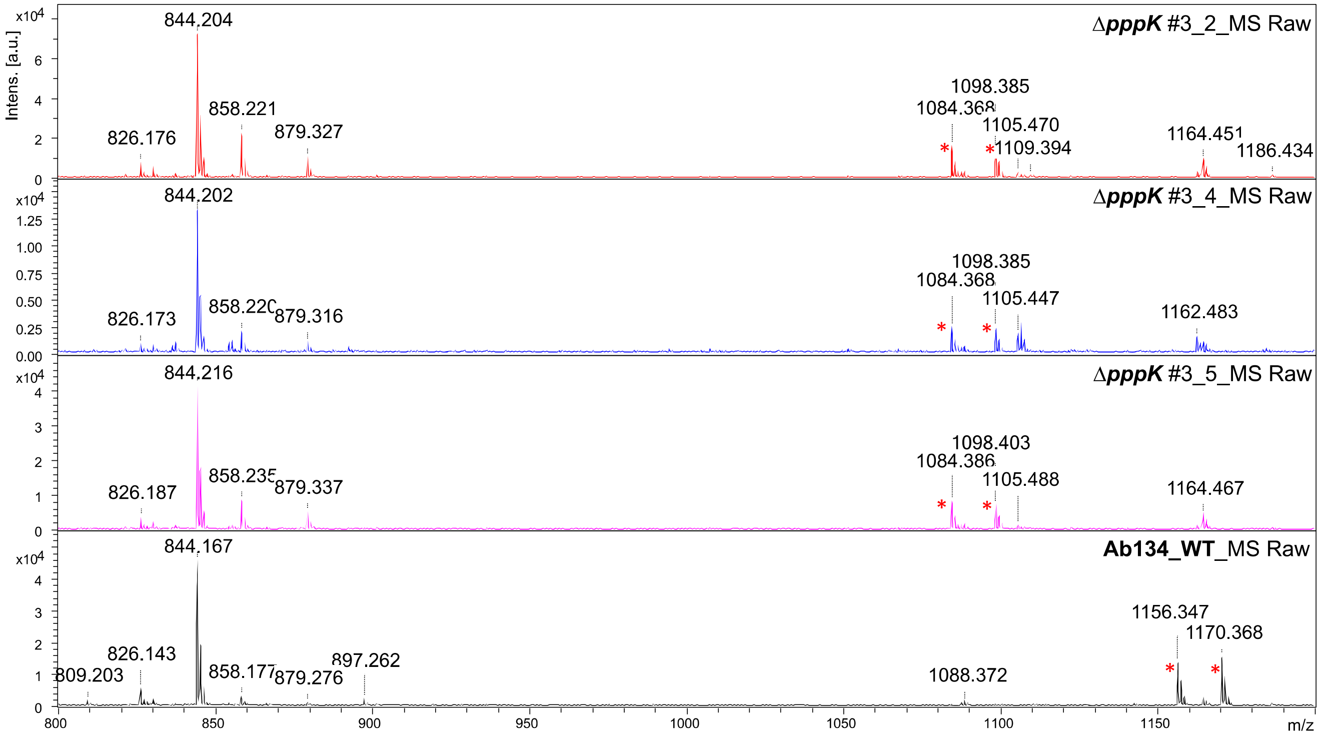

**Figure S20. MALDI-ToF MS analyses of wild type and biological triplicates of Δ*pppK* mutants.** Mass spectra of WT and Δ*pppK* biological triplicates (#3_2, #3_4, and #3_5). The molecular features representing PBs are indicated with red asterisks. The peak at *m/z* 844.2 represents PA1 ($[M+H]^+$); the peak at *m/z* 1156.3, PB1 (Val, $[M+H]^+$); the peak at *m/z* 1170.4, PB2 or PB3 (Leu or Ile, $[M+H]^+$). The peak at *m/z* 1084.4 represents depropionylated PB1 (Val, $[M+H]^+$); and the peak at *m/z* 1098.4, depropionylated PB2 or PB3 (Leu or Ile, $[M+H]^+$).

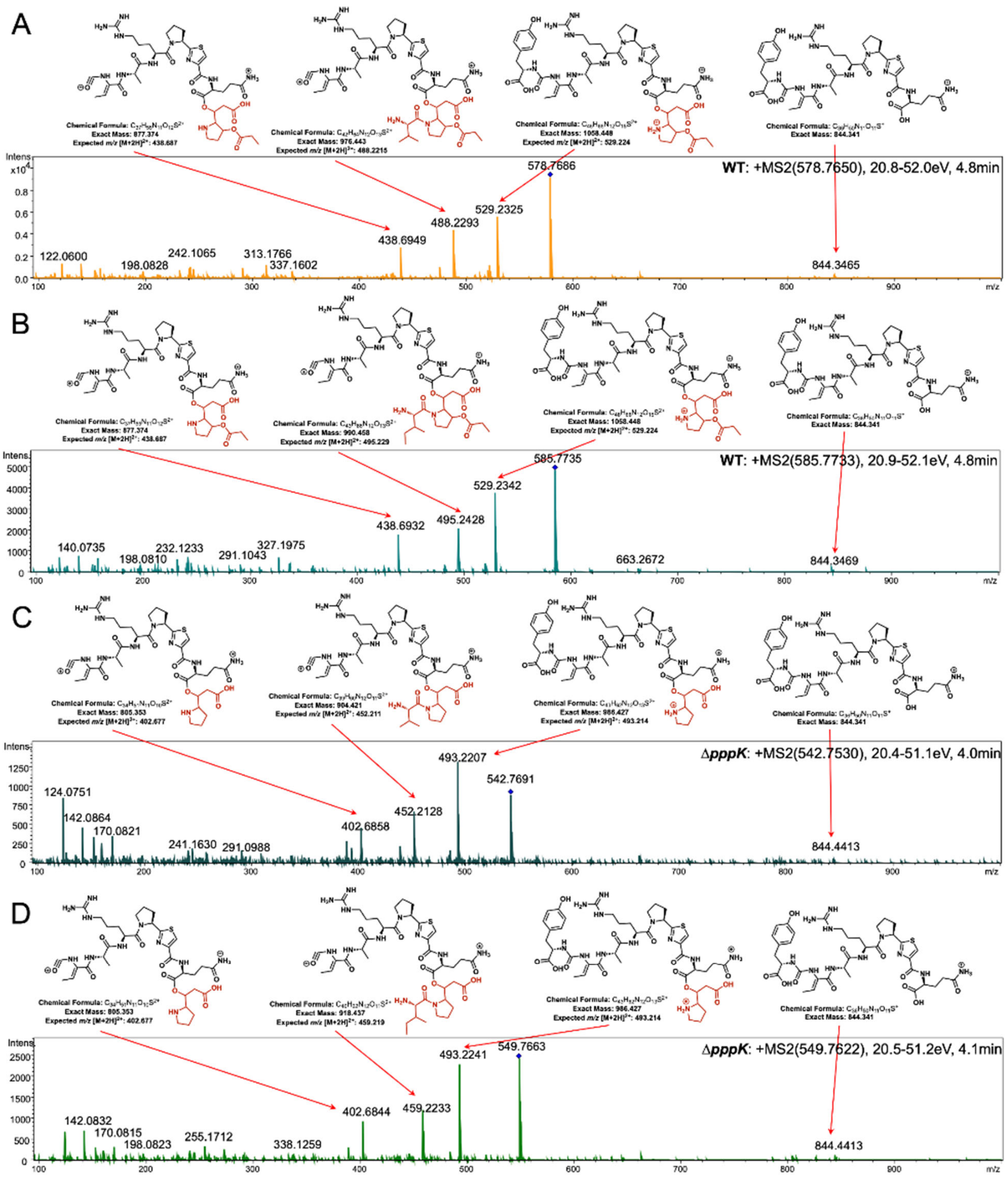

816

817 **Figure S21. Comparison of MS/MS fragmentation between PBs from wild type and**

818 **depropionylated PBs from *ΔpppK* mutant.** UPLC-QToF-MS/MS analyses were performed for

819 comparing the fragmentation pattern of (**A**) PB1 (Val, $[M+2H]^{2+}$) and (**B**) PB2 or PB3 (Ile or Leu,

[M+2H]$^{2+}$) produced from WT and (**C**) depropionylated PB1 (dPB1, Val, [M+2H]$^{++}$) and (**D**) depropionylated PB2 or PB3 (dPB2, Leu, [M+2H]$^{2+}$ or dPB3, Ile, [M+2H]$^{2+}$) from Δ*pppK*. Proposed structures of major peaks are shown. Structures of PB2 and dPB2 are shown as representatives in (**B**) at *m/z* 495.23 and in (**D**) at *m/z* 459.22. The upper right numbers from left to right are the *m/z* of the fragmentated compound, the collision energy, and the retention time. The blue diamond pinpoints the parent peak.

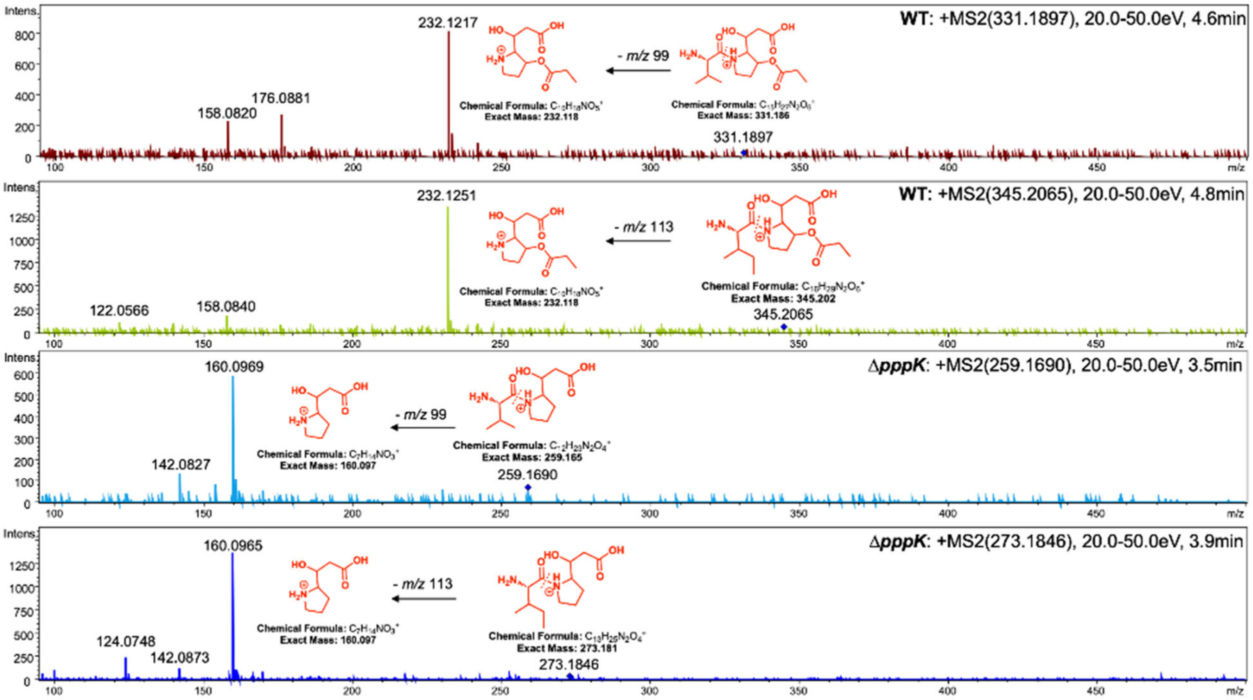

**Figure S22. Comparison of MS/MS fragmentation between PCs and depropionylated PCs.** UPLC-QToF-MS/MS analyses were performed for comparing the fragmentation pattern (from top to bottom) of PC1 (Val, [M+H]$^{+}$) and PC2 or PC3 (Ile or Leu, [M+H]$^{+}$) produced from WT and depropionylated PC1 (dPC1, Val, [M+H]$^{+}$) and depropionylated PC2 or PC3 (dPC2, Leu, [M+H]$^{+}$ , or dPC3, Leu, [M+H]$^{+}$) from Δ*pppK.* Predicted singly charged structures are displayed beside peaks. The *m/z* ratio difference between the molecular peak and the base peak corresponds to the molecular weight of valine (PC1 and dPC1) or leucine/isoleucine (PC2 or PC3, and dPC2 or dPC3). The upper right numbers from left to right are the *m/z* of the fragmentated compound, the collision energy, and the retention time. The blue diamond pinpoints the parent peak.

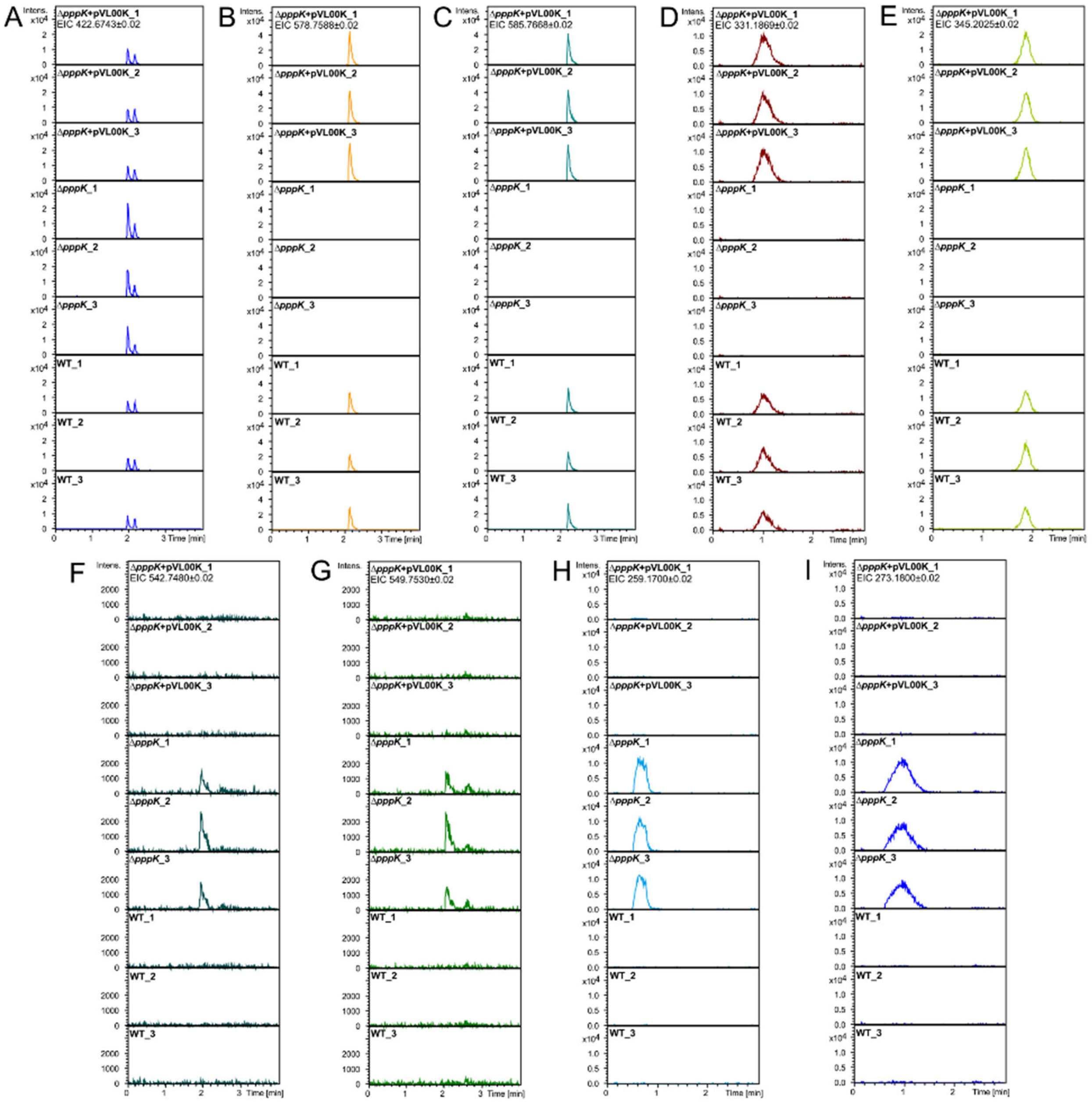

**Figure S23. Comparison of PA, PB, and PC production between Δ*pppK* mutant, wild type and genetically complemented ΔpppK mutant.** UPLC-QToF-MS analyses were performed for wild type, Δ*pppK* mutant, and complemented Δ*pppK* mutant in triplicates. Extracted Ion Chromatograms (EIC) of (**A**) doubly charged PA1 (*m/z* 442.6743, $[M+2H]^{2+}$), (**B**) doubly charged

PB1 ($m/z$ 578.7588, Val, $[M+2H]^{2+}$), (**C**) PB2 or PB3 ($m/z$ 585.7668, Ile or Leu, $[M+2H]^{2+}$), (**D**) singly charged PC1 ($m/z$ 331.1869, Val, $[M+H]^{+}$), (**E**) PC2 or PC3 ($m/z$ 345.2025, Ile or Leu, $[M+H]^{+}$), (**F**) doubly charged dPB1 ($m/z$ 542.7480, Val, $[M+2H]^{2+}$), (**G**) doubly charged dPB2 or dPB3 ($m/z$ 549.7530, Leu or Ile, $[M+2H]^{2+}$), (**H**) dPC1 ($m/z$ 259.1659, Val, $[M+H]^{+}$), and (**I**) dPC2 or dPC3 ($m/z$ 273.1818, Leu or Ile, $[M+H]^{+}$). The same mass filter (the expected $m/z$ ± 0.02) was applied to all samples.

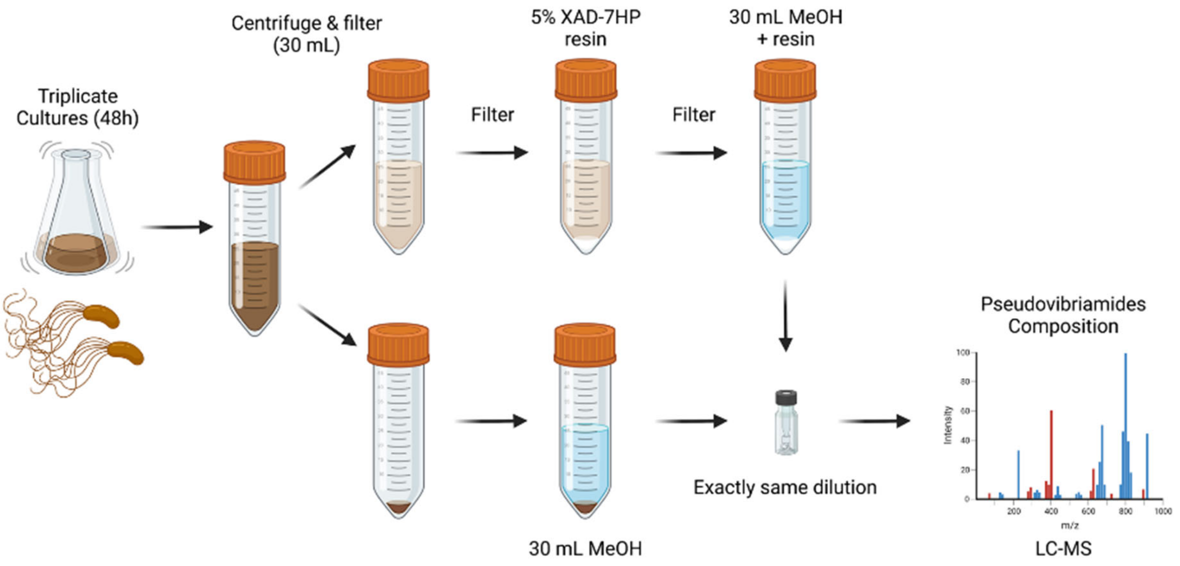

**Figure S24. The workflow of pseudovibriamides extraction from cell pellet and supernatant.** For the wild type, the Δ*pppG* mutant, the genetically complemented Δ*pppG* mutant, and the Δ*pppL* mutant.

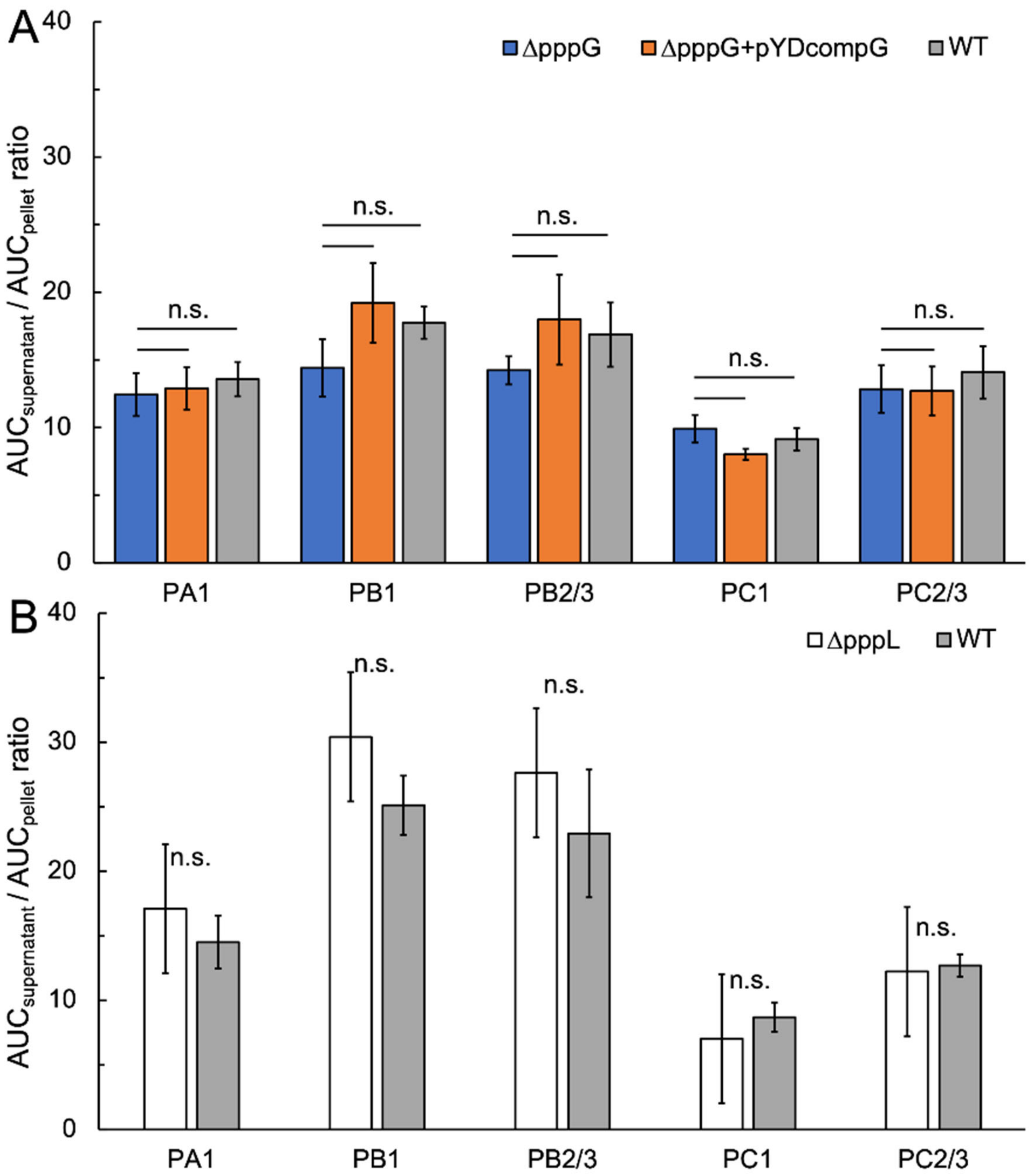

**Figure S25. Comparison of pseudovibriamide export ratio between wild type and transporter mutants.** Each bar represents the average export ratio of each pseudovibriamide as indicated. (**A**) The average export ratio calculated from triplicate supernatant/pellet extracts was used to assess the relative amount of each pseudovibriamide exported by Δ*pppG,* complemented Δ*pppG* and wild type. Two-tail *P*-values were used to determine statistical significance; n.s., not significant (*P*-value > 0.05). Error bars indicate standard deviation. (**B**) The average export ratio calculated from quadruplicate supernatant/pellet extracts were used to assess the relative amount of each pseudovibriamide exported in Δ*pppL* and wild type. The same statistical analysis as above was applied.

.

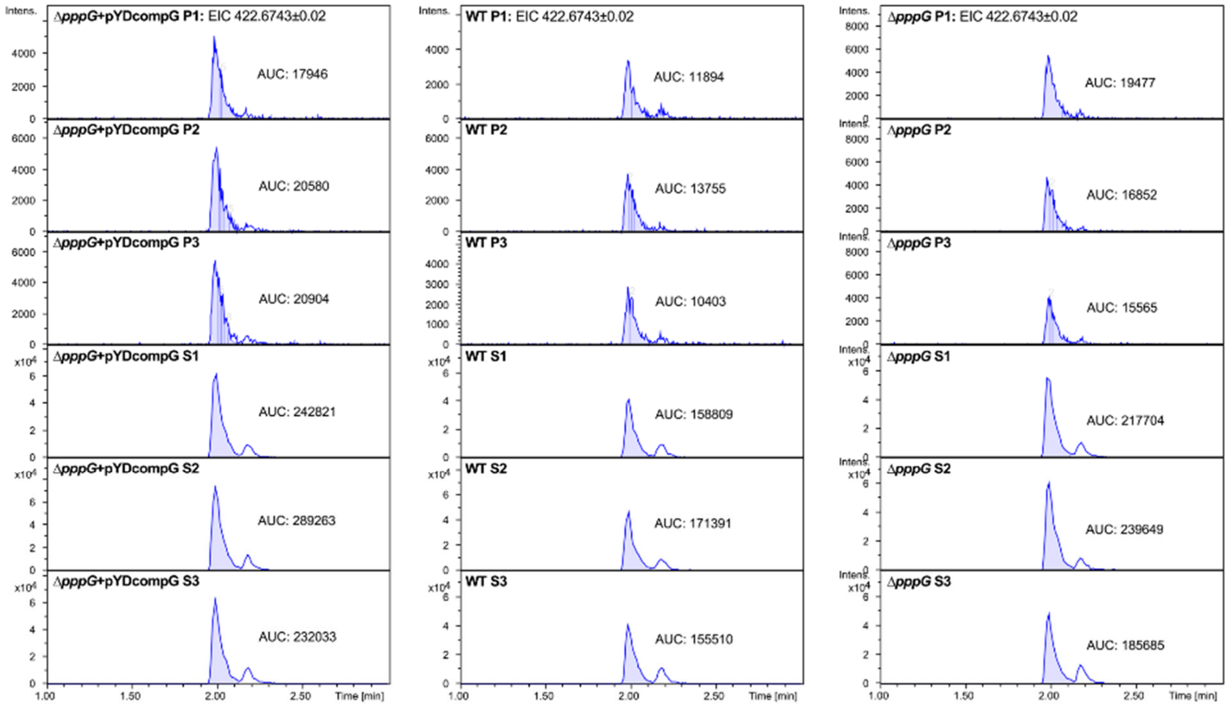

**Figure S26. EIC of PA1 from supernatant and pellet extracts of genetically complemented Δ*pppG* strain, wild type, and Δ*pppG* mutant.** UPLC-QToF-MS analyses were performed for supernatant and pellet extracts in triplicates. EIC of doubly charged PA1 ($[M+2H]^{2+}$). P, pellet extract; S, supernatant extract; AUC, area under curve. AUC was obtained using “Chromatogram Compound” function, represented by the shaded area.

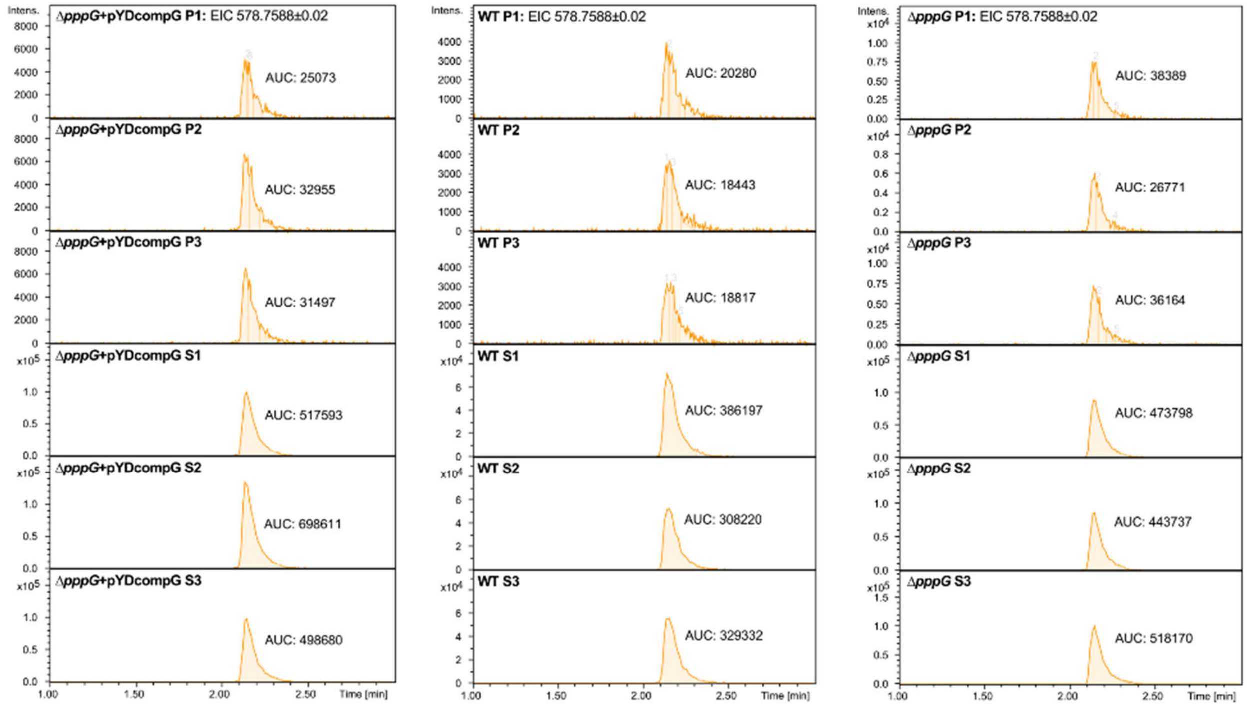

**Figure S27. EIC of PB1 from supernatant and pellet extracts of genetically complemented Δ*pppG* strain, wild type, and Δ*pppG* mutant.** UPLC-QToF-MS analyses were performed for supernatant and pellet extracts in triplicates. EIC of doubly charged PB1 (Val, $[M+2H]^{2+}$). P, pellet extract; S, supernatant extract; AUC, area under curve. AUC was obtained using "Chromatogram Compound" function, represented by the shaded area.

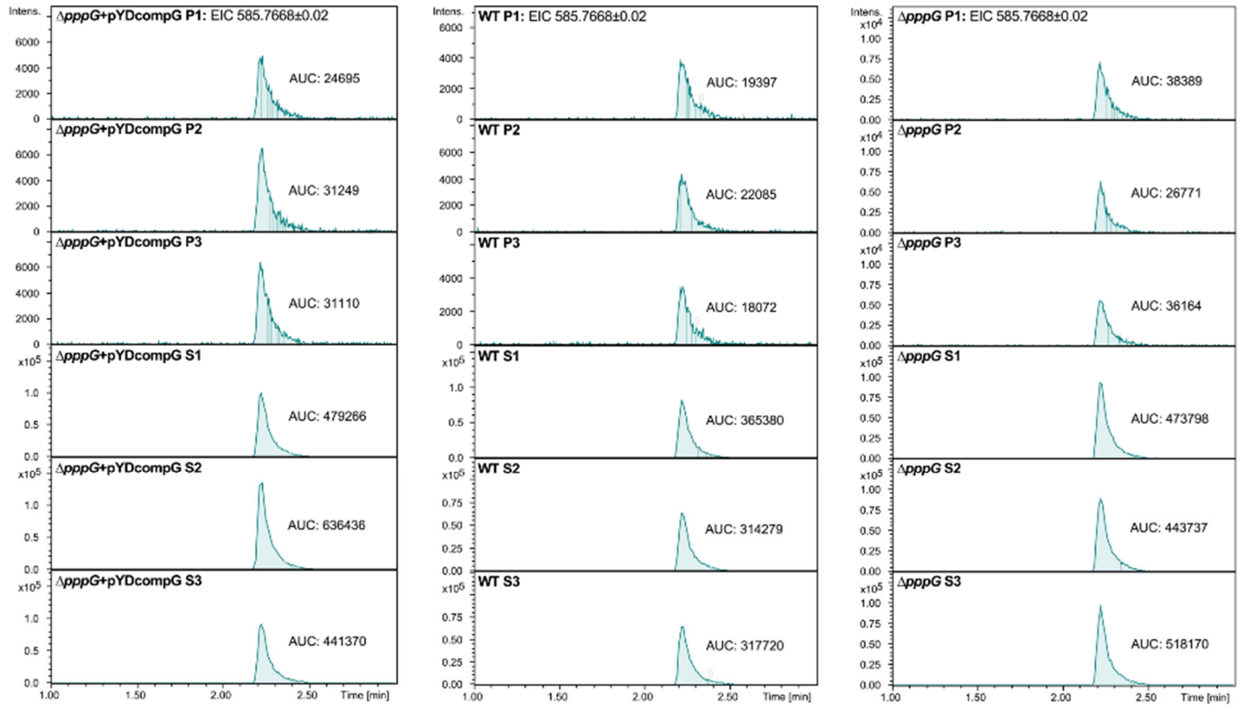

**Figure S28. EIC of PB2 or PB3 from supernatant and pellet extracts of genetically complemented Δ*pppG* strain, wild type, and Δ*pppG* mutant.** UPLC-QToF-MS analyses were performed for supernatant and pellet extracts in triplicates. EIC of doubly charged PB2 or PB3 (Ile or Leu, $[M+2H]^{2+}$). P, pellet extract; S, supernatant extract; AUC, area under curve. AUC was obtained using "Chromatogram Compound" function, represented by the shaded area.

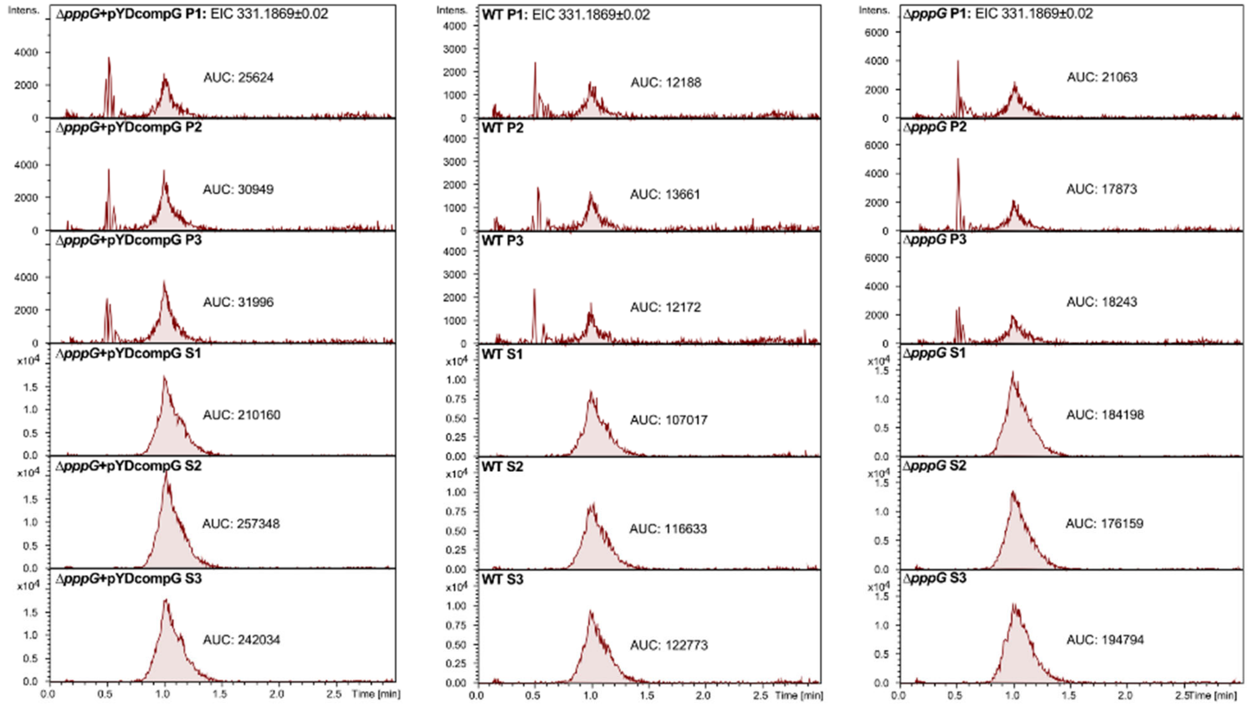

**Figure S29. EIC of PC1 from supernatant and pellet extracts of genetically complemented ΔpppG strain, wild type, and Δ*pppG* mutant.** UPLC-QToF-MS analyses were performed for supernatant and pellet extracts in triplicates. EIC of singly charged PC1 (Val, $[M+H]^+$). P, pellet extract; S, supernatant extract; AUC, area under curve. AUC was obtained using "Chromatogram Compound" function, represented by the shaded area.

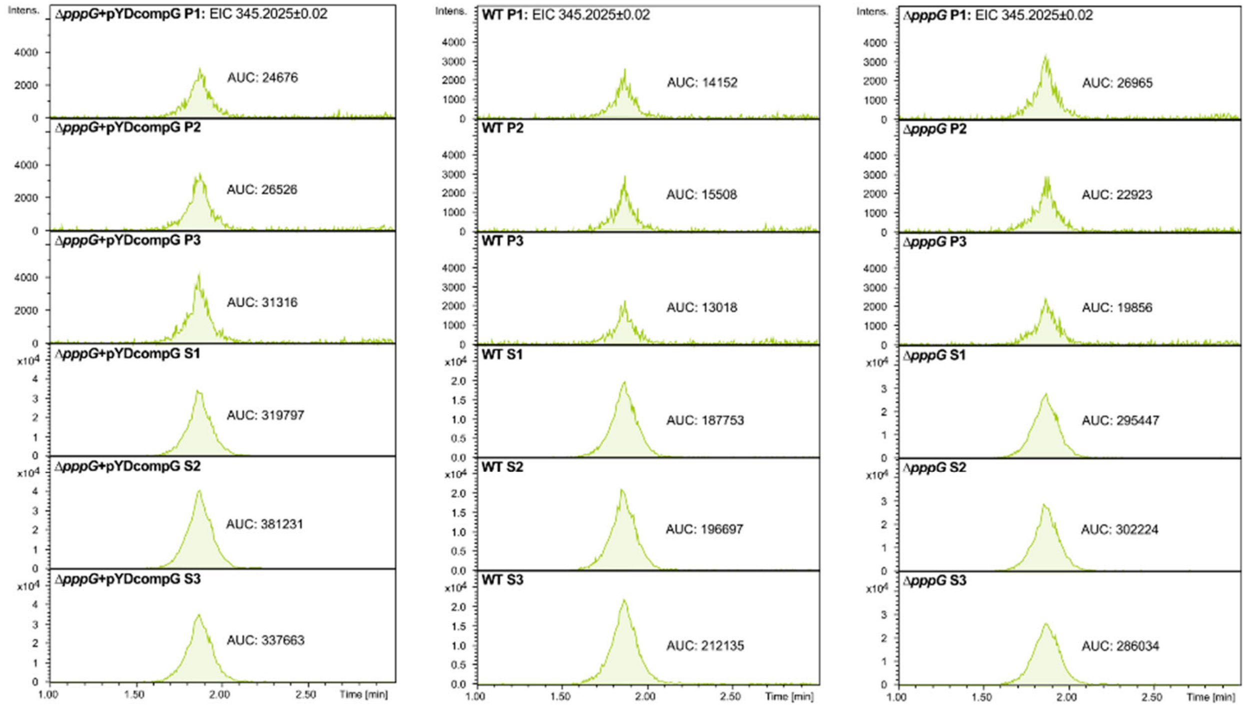

**Figure S30. EIC of PC2 or PC3 from supernatant and pellet extracts of genetically complemented Δ*pppG* strain, wild type, and Δ*pppG* mutant.** UPLC-QToF-MS analyses were performed for supernatant and pellet extracts in triplicates. EIC of singly charged PC2 or PC3 (Leu or Ile, $[M+H]^+$). P, pellet extract; S, supernatant extract; AUC, area under curve. AUC was obtained using “Chromatogram Compound” function, represented by the shaded area.

| Pseudovi -briamide A1 | AUC |
|---|---|
| LP1 | 1920.3 |
| LP2 | 1985.7 |
| LP3 | 1779.2 |
| LP4 | 2143.1 |
| WP1 | 5005.2 |
| WP2 | 4498.2 |
| WP3 | 4339 |
| WP4 | 4948.9 |

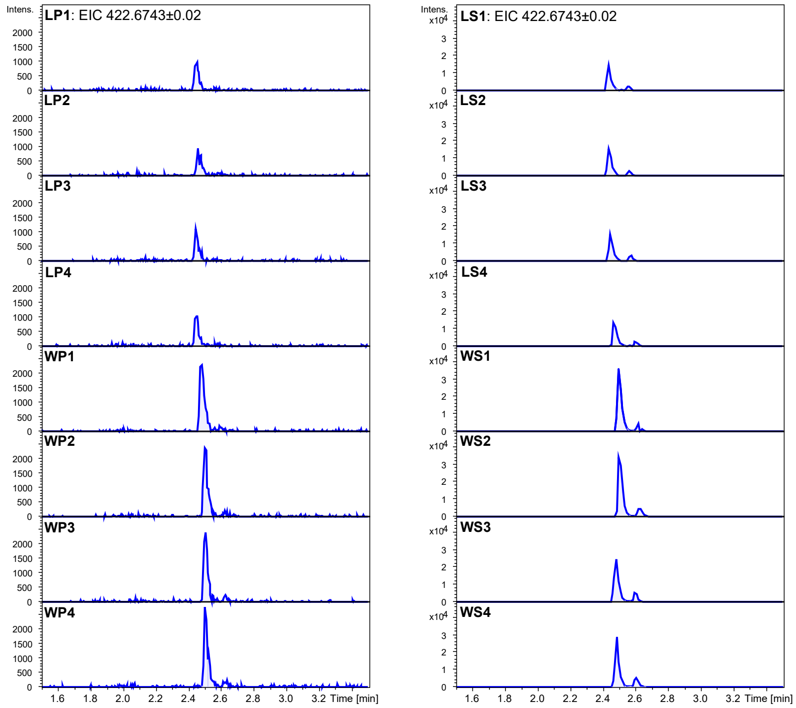

| Pseudovi -briamide A1 | AUC |
|---|---|
| LS1 | 31561.4 |
| LS2 | 33522.5 |
| LS3 | 35599.7 |
| LS4 | 32180.7 |
| WS1 | 73655.7 |
| WS2 | 77891 |
| WS3 | 58383.3 |
| WS4 | 62277 |

**Figure S31. EIC of PA1 from supernatant and pellet extracts of wild type and Δ*pppL* mutant.** UPLC-QToF-MS analyses were performed for supernatant and pellet extracts from both wild type and Δ*pppL* mutant in quadruplicates. EIC of doubly charged PA1 ($[M+2H]^{2+}$). LP, Δ*pppL* pellet extract; LS, Δ*pppL* supernatant extract; WP, wild type pellet extract; WS, wild type supernatant extract. AUC was obtained using "Chromatogram Compound" function.

| Pseudovi-briamide B1 | AUC |
|---|---|
| LP1 | 2555.6 |
| LP2 | 2211.7 |
| LP3 | 2664.5 |
| LP4 | 2831.6 |
| WP1 | 6237.4 |
| WP2 | 6177.4 |
| WP3 | 5099.8 |
| WP4 | 6021.3 |

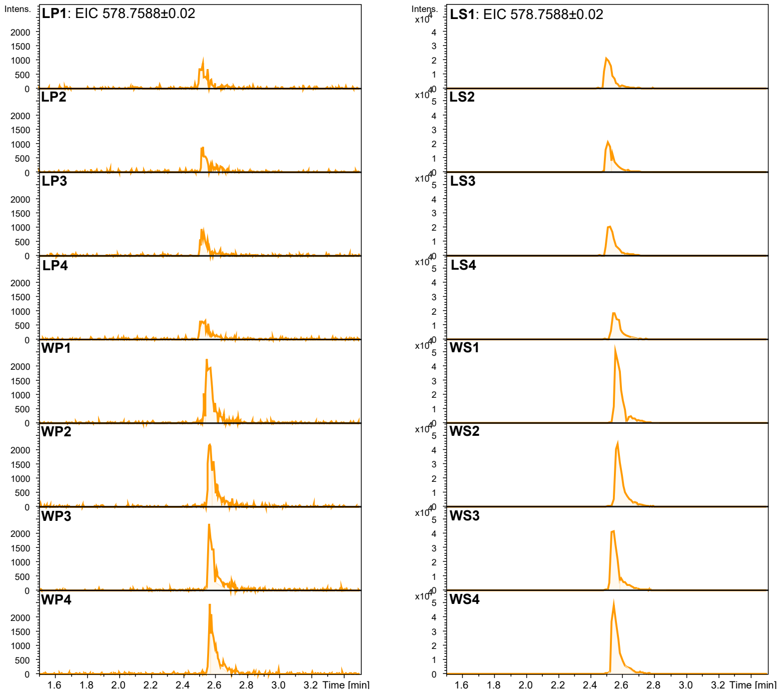

| Pseudovi-briamide B1 | AUC |
|---|---|
| LS1 | 79528.6 |
| LS2 | 77462.3 |
| LS3 | 80499.1 |
| LS4 | 71563.6 |
| WS1 | 158663.2 |
| WS2 | 139277.8 |
| WS3 | 143083.4 |
| WS4 | 146893.1 |

**Figure S32. EIC of PB1 from supernatant and pellet extracts of wild type and Δ*pppL* mutant.** UPLC-QToF-MS analyses were performed for supernatant and pellet extracts from both wild type and Δ*pppL* mutant in quadruplicates. EIC of doubly charged PB1 (Val, $[M+2H]^{2+}$). LP, Δ*pppL* pellet extract; LS, Δ*pppL* supernatant extract; WP, wild type pellet extract; WS, wild type supernatant extract. AUC was obtained using “Chromatogram Compound” function.

| Pseudovi-briamide B2/3 | AUC |
|---|---|
| LP1 | 1857.7 |
| LP2 | 1963.3 |
| LP3 | 2403.8 |
| LP4 | 2322.5 |
| WP1 | 6183.1 |
| WP2 | 4797 |
| WP3 | 5442 |
| WP4 | 5187.8 |

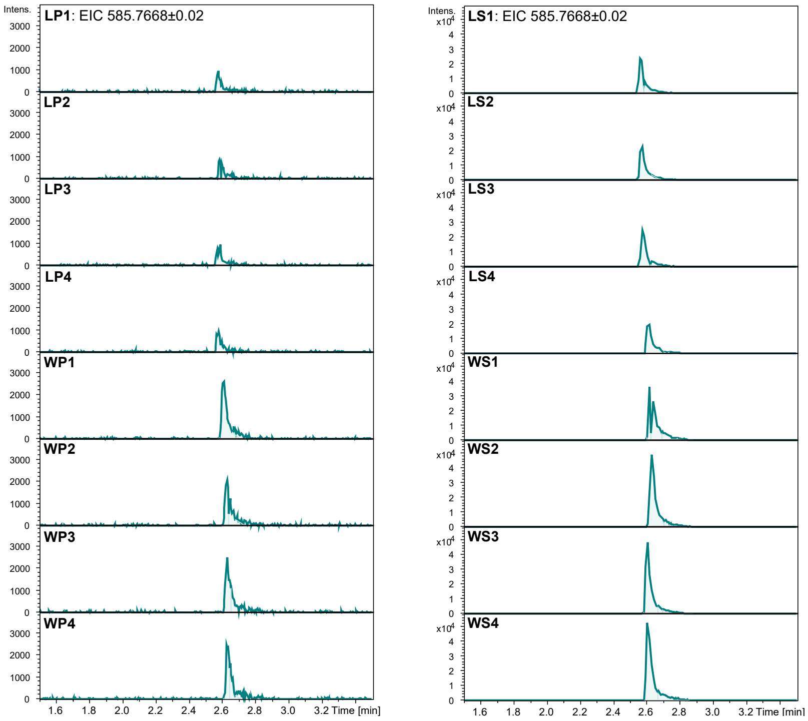

| Pseudovi-briamide B2/3 | AUC |
|---|---|
| LS1 | 62551.2 |
| LS2 | 58696.8 |
| LS3 | 51417.6 |
| LS4 | 59312.8 |
| WS1 | 98416.9 |
| WS2 | 125048.9 |
| WS3 | 125158.5 |
| WS4 | 138666.5 |

**Figure S33. EIC of PB2 or PB3 from supernatant and pellet extracts of wild type and Δ*pppL* mutant.** UPLC-QToF-MS analyses were performed for supernatant and pellet extracts from both wild type and Δ*pppL* mutant in quadruplicates. EIC of doubly charged PB2 or PB3 (Ile or Leu, $[M+2H]^{2+}$). LP, Δ*pppL* pellet extract; LS, Δ*pppL* supernatant extract; WP, wild type pellet extract; WS, wild type supernatant extract. AUC was obtained using "Chromatogram Compound" function.

| Pseudovi-briamide C1 | AUC |
|---|---|
| LP1 | 2281.6 |
| LP2 | 2078.2 |
| LP3 | 2014.3 |
| LP4 | 2573.2 |
| WP1 | 5249.8 |
| WP2 | 3634 |
| WP3 | 4917.8 |
| WP4 | 4239.3 |

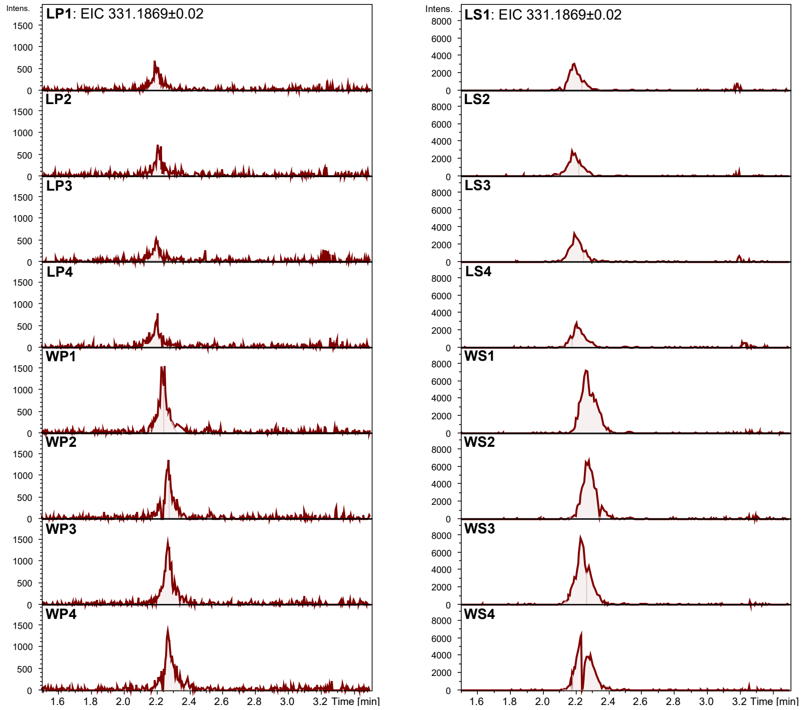

| Pseudovi-briamide C1 | AUC |
|---|---|
| LS1 | 15527.7 |
| LS2 | 15178 |
| LS3 | 16596.9 |
| LS4 | 14818.7 |
| WS1 | 42620.1 |
| WS2 | 37514.8 |
| WS3 | 41612.8 |
| WS4 | 33424.3 |

**Figure S34. EIC of PC1 from supernatant and pellet extracts of wild type and Δ*pppL* mutant.** UPLC-QToF-MS analyses were performed for supernatant and pellet extracts from both wild type and Δ*pppL* mutant in quadruplicates. EIC of singly charged PC1 (Val, $[M+H]^+$). LP, Δ*pppL* pellet extract; LS, Δ*pppL* supernatant extract; WP, wild type pellet extract; WS, wild type supernatant extract. AUC was obtained using "Chromatogram Compound" function.

| Pseudovi-briamide C2/3 | AUC |
|---|---|
| LP1 | 2395.5 |
| LP2 | 2628.2 |
| LP3 | 2479.9 |
| LP4 | 2248.9 |
| WP1 | 5291.9 |
| WP2 | 5443.9 |
| WP3 | 4698.2 |
| WP4 | 5326.5 |

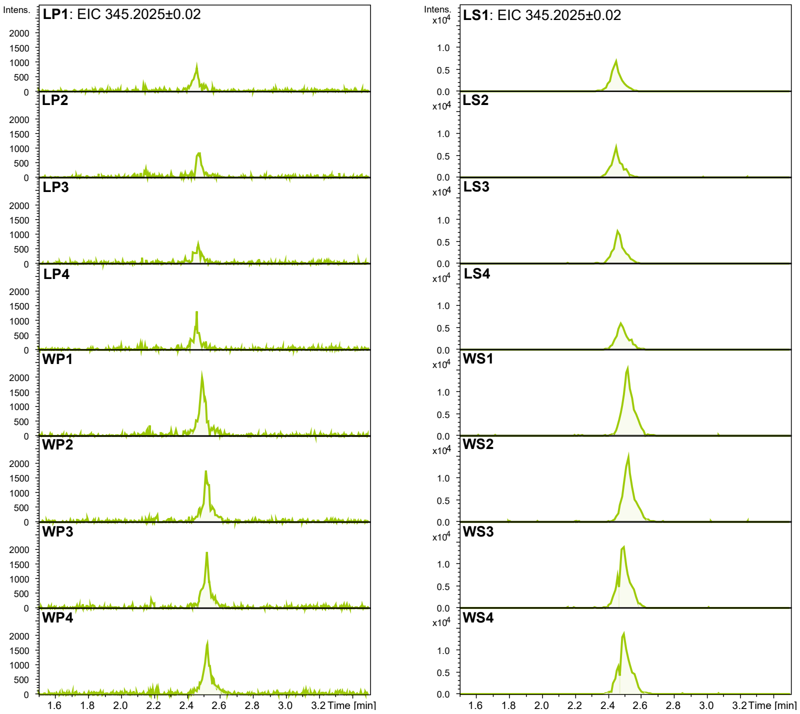

| Pseudovi-briamide C2/3 | AUC |
|---|---|
| LS1 | 29571.7 |
| LS2 | 27973.3 |
| LS3 | 32053.7 |
| LS4 | 29252.5 |
| WS1 | 68494.1 |
| WS2 | 65128.7 |
| WS3 | 64977.1 |
| WS4 | 64362.9 |

**Figure S35. EIC of PC2 or PC3 from supernatant and pellet extracts of wild type and Δ*pppL* mutant.** UPLC-QToF-MS analyses were performed for supernatant and pellet extracts from both wild type and Δ*pppL* mutant in quadruplicates. EIC of singly charged PC2 or PC3 (Leu or Ile, $[M+H]^+$). LP, Δ*pppL* pellet extract; LS, Δ*pppL* supernatant extract; WP, wild type pellet extract; WS, wild type supernatant extract. AUC was obtained using "Chromatogram Compound" function.

A

| t-Test: Two-Sample Assuming Equal Variances | | |
|---|---|---|
| PA1 | | |
| | *ΔpppG* | *ΔpppG+pYDcompG* |
| Mean | 12.44268 | 12.89502965 |
| Variance | 2.5135 | 2.486250028 |
| Observations | 3 | 3 |
| Pooled Variance | 2.499875 | |
| Hypothesized Mean Differen | 0 | |
| df | 4 | |
| t Stat | -0.350399 | |
| P(T<=t) one-tail | 0.371857 | |
| t Critical one-tail | 2.131847 | |
| P(T<=t) two-tail | 0.743713 | |
| t Critical two-tail | 2.776445 | |

B

| t-Test: Two-Sample Assuming Equal Variances | | |
|---|---|---|
| PB1 | | |
| | *ΔpppG* | *ΔpppG+pYDcompG* |
| Mean | 14.41519 | 19.22499342 |
| Variance | 4.485253 | 8.707890248 |
| Observations | 3 | 3 |
| Pooled Variance | 6.596572 | |
| Hypothesized Mean Differenc | 0 | |
| df | 4 | |
| t Stat | -2.29358 | |
| P(T<=t) one-tail | 0.041764 | |
| t Critical one-tail | 2.131847 | |
| P(T<=t) two-tail | 0.083527 | |
| t Critical two-tail | 2.776445 | |

C

| t-Test: Two-Sample Assuming Unequal Variances | | |
|---|---|---|
| PB2/3 | | |
| | *ΔpppG* | *ΔpppG+pYDcompG* |
| Mean | 14.24008 | 17.98705339 |
| Variance | 1.080221 | 11.05809013 |
| Observations | 3 | 3 |
| Hypothesized Mean Differenc | 0 | |
| df | 2 | |
| t Stat | -1.862782 | |
| P(T<=t) one-tail | 0.101764 | |
| t Critical one-tail | 2.919986 | |
| P(T<=t) two-tail | 0.203529 | |
| t Critical two-tail | 4.302653 | |

D

| t-Test: Two-Sample Assuming Unequal Variances | | |
|---|---|---|
| PC1 | | |
| | *ΔpppG* | *ΔpppG+pYDcompG* |
| Mean | 9.759722 | 8.02712732 |
| Variance | 0.941016 | 0.163733112 |
| Observations | 3 | 3 |
| Hypothesized Mean Difference | 0 | |
| df | 3 | |
| t Stat | 2.85513 | |
| P(T<=t) one-tail | 0.032418 | |
| t Critical one-tail | 2.353363 | |
| P(T<=t) two-tail | 0.064835 | |
| t Critical two-tail | 3.182446 | |

E

| t-Test: Two-Sample Assuming Equal Variances | | |
|---|---|---|
| PC2/3 | | |
| | *ΔpppG* | *ΔpppG+pYDcompG* |
| Mean | 12.84888 | 12.70466587 |
| Variance | 3.057219 | 3.269551996 |
| Observations | 3 | 3 |
| Pooled Variance | 3.163385 | |
| Hypothesized Mean Differenc | 0 | |
| df | 4 | |
| t Stat | 0.099304 | |
| P(T<=t) one-tail | 0.462837 | |
| t Critical one-tail | 2.131847 | |
| P(T<=t) two-tail | 0.925675 | |
| t Critical two-tail | 2.776445 | |

**Figure S36. A summary of statistical analyses performed for pseudovibriamides export ratio comparison between Δ*pppG* mutant and complemented Δ*pppG* mutant.** (**A**) pseudovibriamide A1; (**B**) pseudovibriamide B1; (**C**) pseudovibriamide B2/3; (**D**) pseudovibriamide C1; (**E**) pseudovibriamide C2/3. P-value of Two-sample two-tail t-Test is used. If the ratio of the larger variance to the smaller variance is less than 4 then assuming the variances are approximately equal or vice versa.

A

| t-Test: Two-Sample Assuming Equal Variances | | |
|---|---|---|
| PA1 | | |
| | Δ*pppG* | WT |
| Mean | 12.44268 | 13.586988 |
| Variance | 2.5135 | 1.5885838 |
| Observations | 3 | 3 |
| Pooled Variance | 2.051042 | |
| Hypothesized Mean Differenc | 0 | |
| df | 4 | |
| t Stat | -0.978593 | |
| P(T<=t) one-tail | 0.191595 | |
| t Critical one-tail | 2.131847 | |
| P(T<=t) two-tail | 0.38319 | |
| t Critical two-tail | 2.776445 | |

B

| t-Test: Two-Sample Assuming Equal Variances | | |
|---|---|---|
| PB1 | | |
| | Δ*pppG* | WT |
| Mean | 14.41519 | 17.75247 |
| Variance | 4.485253 | 1.406649 |
| Observations | 3 | 3 |
| Pooled Variance | 2.945951 | |
| Hypothesized Mean Differenc | 0 | |
| df | 4 | |
| t Stat | -2.381363 | |
| P(T<=t) one-tail | 0.037938 | |
| t Critical one-tail | 2.131847 | |
| P(T<=t) two-tail | 0.075877 | |
| t Critical two-tail | 2.776445 | |

C

| t-Test: Two-Sample Assuming Unequal Variances | | |
|---|---|---|
| PB2/3 | | |
| | Δ*pppG* | WT |
| Mean | 14.24008 | 16.88279 |
| Variance | 1.080221 | 5.670846 |
| Observations | 3 | 3 |
| Hypothesized Mean Differenc | 0 | |
| df | 3 | |
| t Stat | -1.761666 | |
| P(T<=t) one-tail | 0.08817 | |
| t Critical one-tail | 2.353363 | |
| P(T<=t) two-tail | 0.17634 | |
| t Critical two-tail | 3.182446 | |

D

| t-Test: Two-Sample Assuming Equal Variances | | |
|---|---|---|
| PC1 | | |
| | Δ*pppG* | WT |
| Mean | 9.759722 | 9.135051 |
| Variance | 0.941016 | 0.693882 |
| Observations | 3 | 3 |
| Pooled Variance | 0.817449 | |
| Hypothesized Mean Differenc | 0 | |
| df | 4 | |
| t Stat | 0.846188 | |
| P(T<=t) one-tail | 0.22255 | |
| t Critical one-tail | 2.131847 | |
| P(T<=t) two-tail | 0.4451 | |
| t Critical two-tail | 2.776445 | |

E

| t-Test: Two-Sample Assuming Equal Variances | | |
|---|---|---|
| PC2/3 | | |
| | Δ*pppG* | WT |
| Mean | 12.84888 | 14.08208 |
| Variance | 3.057219 | 3.760672 |
| Observations | 3 | 3 |
| Pooled Variance | 3.408945 | |
| Hypothesized Mean Difference | 0 | |
| df | 4 | |
| t Stat | -0.818033 | |
| P(T<=t) one-tail | 0.229634 | |
| t Critical one-tail | 2.131847 | |
| P(T<=t) two-tail | 0.459268 | |
| t Critical two-tail | 2.776445 | |

**Figure S37. A summary of statistical analyses performed for pseudovibriamides export ratio comparison between Δ*pppG* mutant and wild type.** (**A**) pseudovibriamide A1; (**B**) pseudovibriamide B1; (**C**) pseudovibriamide B2/3; (**D**) pseudovibriamide C1; (**E**) pseudovibriamide C2/3. P-value of Two-sample two-tail t-Test is used. If the ratio of the larger variance to the smaller variance is less than 4 then assuming the variances are approximately equal or vice versa.

**A** t-Test: Two-Sample Assuming Equal Variances

$AUC_{Sup}$ / $AUC_{pellet}$ Ratio

| | ***ΔpppL PA1*** | ***WT PA1*** |
|---|---|---|
| Mean | 17.0855999 | 14.5178241 |
| Variance | 4.43084945 | 4.2458049 |
| Observations | 4 | 4 |
| Pooled Variance | 4.33832717 | |
| Hypothesized Mean Difference | 0 | |
| df | 6 | |
| t Stat | 1.7434558 | |
| P(T<=t) one-tail | 0.06593936 | |
| t Critical one-tail | 1.94318028 | |
| P(T<=t) two-tail | 0.13187871 | |
| t Critical two-tail | 2.44691185 | |

**B** t-Test: Two-Sample Assuming **Unequal** Variances

$AUC_{Sup}$ / $AUC_{pellet}$ Ratio

| | ***ΔpppL PB1*** | ***WT PB1*** |
|---|---|---|
| Mean | 30.4070327 | 25.1090378 |
| Variance | 16.072323 | 5.2909549 |
| Observations | 4 | 4 |
| Hypothesized Mean Difference | 0 | |
| df | 5 | |
| t Stat | 2.29249072 | |
| P(T<=t) one-tail | 0.03521492 | |
| t Critical one-tail | 2.01504837 | |
| P(T<=t) two-tail | 0.07042985 | |
| t Critical two-tail | 2.57058184 | |

**C** t-Test: Two-Sample Assuming Equal Variances

$AUC_{Sup}$ / $AUC_{pellet}$ Ratio

| | ***ΔpppL PB2/3*** | ***WT PB2/3*** |
|---|---|---|
| Mean | 27.62419968 | 22.9282923 |
| Variance | 28.31588535 | 24.4896975 |
| Observations | 4 | 4 |
| Pooled Variance | 26.40279145 | |
| Hypothesized Mean Difference | 0 | |
| df | 6 | |
| t Stat | 1.2924377 | |
| P(T<=t) one-tail | 0.12187237 | |
| t Critical one-tail | 1.94318028 | |
| P(T<=t) two-tail | 0.24374474 | |
| t Critical two-tail | 2.44691185 | |

**D** t-Test: Two-Sample Assuming Equal Variances

$AUC_{Sup}$ / $AUC_{pellet}$ Ratio

| | ***ΔpppL PC1*** | ***WT PC1*** |
|---|---|---|
| Mean | 7.0268631 | 8.69692961 |
| Variance | 1.067950189 | 1.23176821 |
| Observations | 4 | 4 |
| Pooled Variance | 1.149859198 | |
| Hypothesized Mean Difference | 0 | |
| df | 6 | |
| t Stat | -2.20255333 | |
| P(T<=t) one-tail | 0.03492803 | |
| t Critical one-tail | 1.94318028 | |
| P(T<=t) two-tail | 0.06985605 | |
| t Critical two-tail | 2.44691185 | |

**E** t-Test: Two-Sample Assuming Equal Variances

$AUC_{Sup}$ / $AUC_{pellet}$ Ratio

| | ***ΔpppL PC2/3*** | ***WT PC2/3*** |
|---|---|---|
| Mean | 12.2302697 | 12.7051177 |
| Variance | 1.2060375 | 0.7528973 |
| Observations | 4 | 4 |
| Pooled Variance | 0.9794674 | |
| Hypothesized Mean Difference | 0 | |
| df | 6 | |
| t Stat | -0.67853871 | |
| P(T<=t) one-tail | 0.26136265 | |
| t Critical one-tail | 1.94318028 | |
| P(T<=t) two-tail | 0.5227253 | |
| t Critical two-tail | 2.44691185 | |

**Figure S38. A summary of statistical analyses performed for pseudovibriamides export ratio comparison between Δ*pppL* mutant and wild type.** (**A**) pseudovibriamide A1; (**B**) pseudovibriamide B1; (**C**) pseudovibriamide B2/3; (**D**) pseudovibriamide C1; (**E**) pseudovibriamide C2/3. P-value of Two-sample two-tail t-Test is used. If the ratio of the larger variance to the smaller variance is less than 4 then assuming the variances are approximately equal or vice versa.

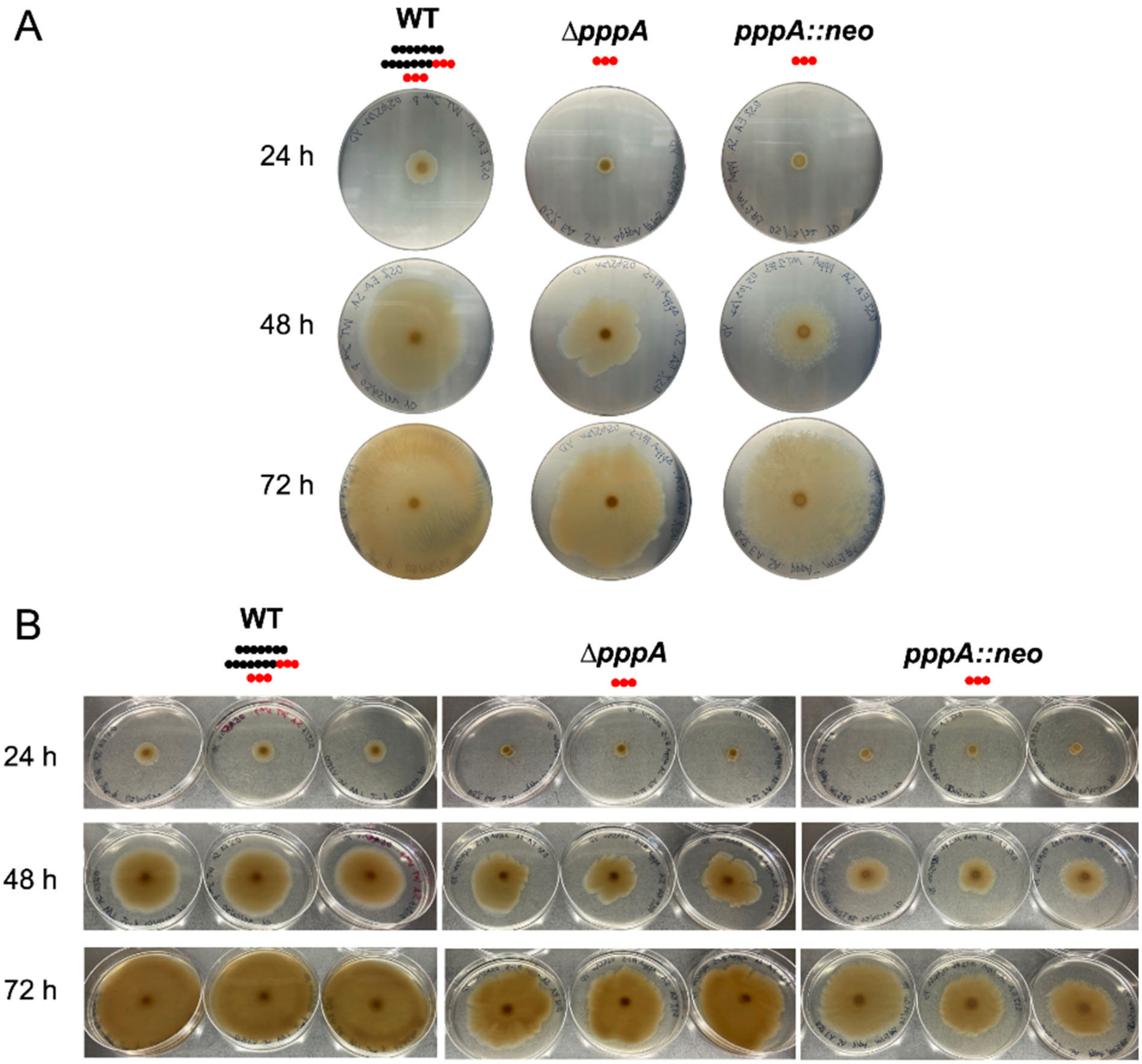

**Figure S39. Swarming assay results of Δ*pppA* and *pppA::neo* mutants compared to the wild type.** The swarming assay was performed as described under Methods. Pictures shown were taken at 24, 48, and 72 hours after inoculation. (**A**) Representative results are shown. Pseudovibriamides are represented by beads: PA, seven black beads; PB, seven black beads and three red beads; PC, three red beads. (**B**) Triplicates of each strain are shown.

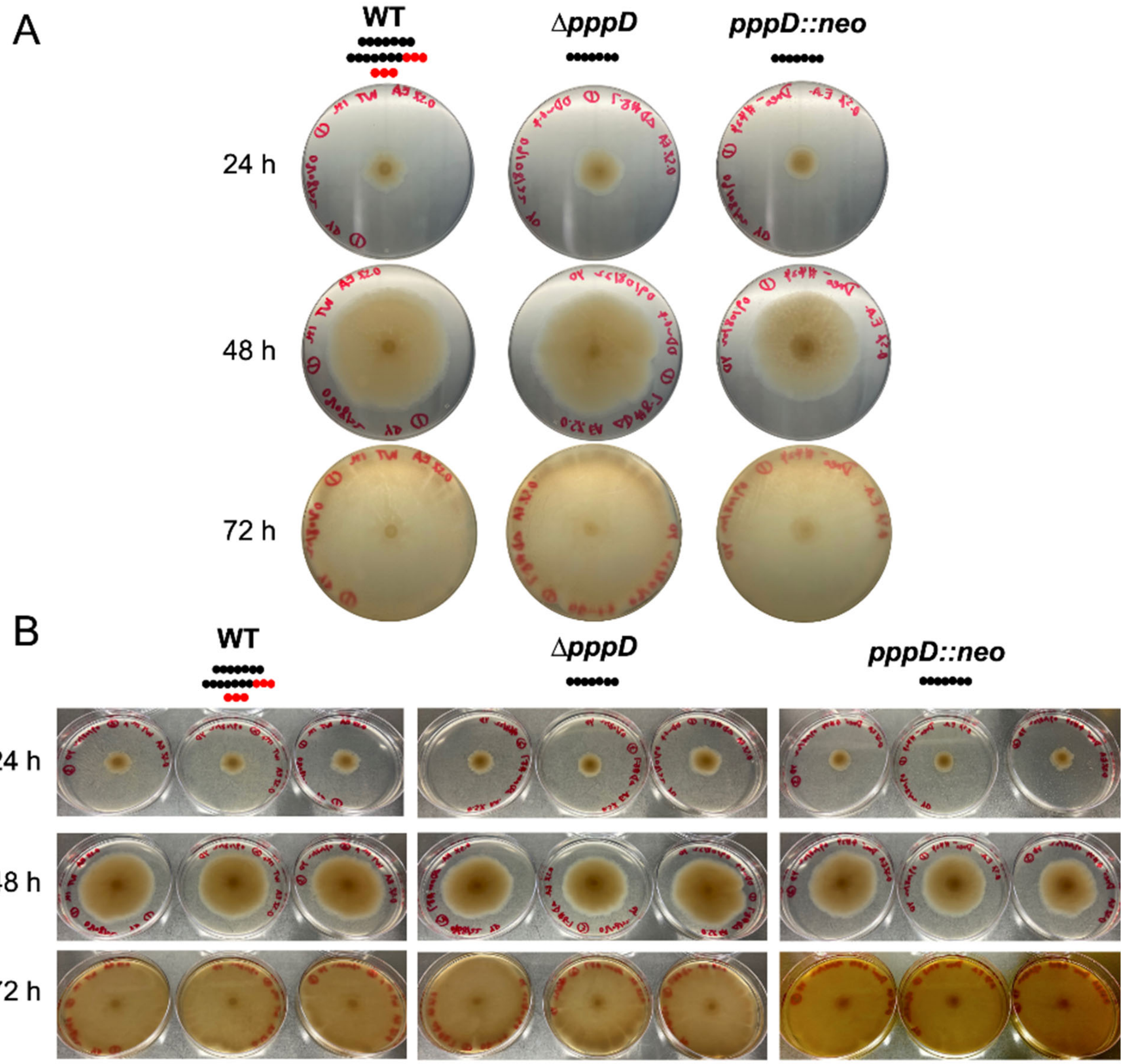

**Figure S40. Swarming assay results of Δ*pppD* and *pppD::neo* mutants compared to the wild type.** The swarming assay was performed as described under Methods. Pictures shown were taken at 24, 48, and 72 hours after inoculation. (**A**) Representative results are shown. Pseudovibriamides are represented by beads: PA, seven black beads; PB, seven black beads and three red beads; PC, three red beads. (**B**) Triplicates of each strain are shown.

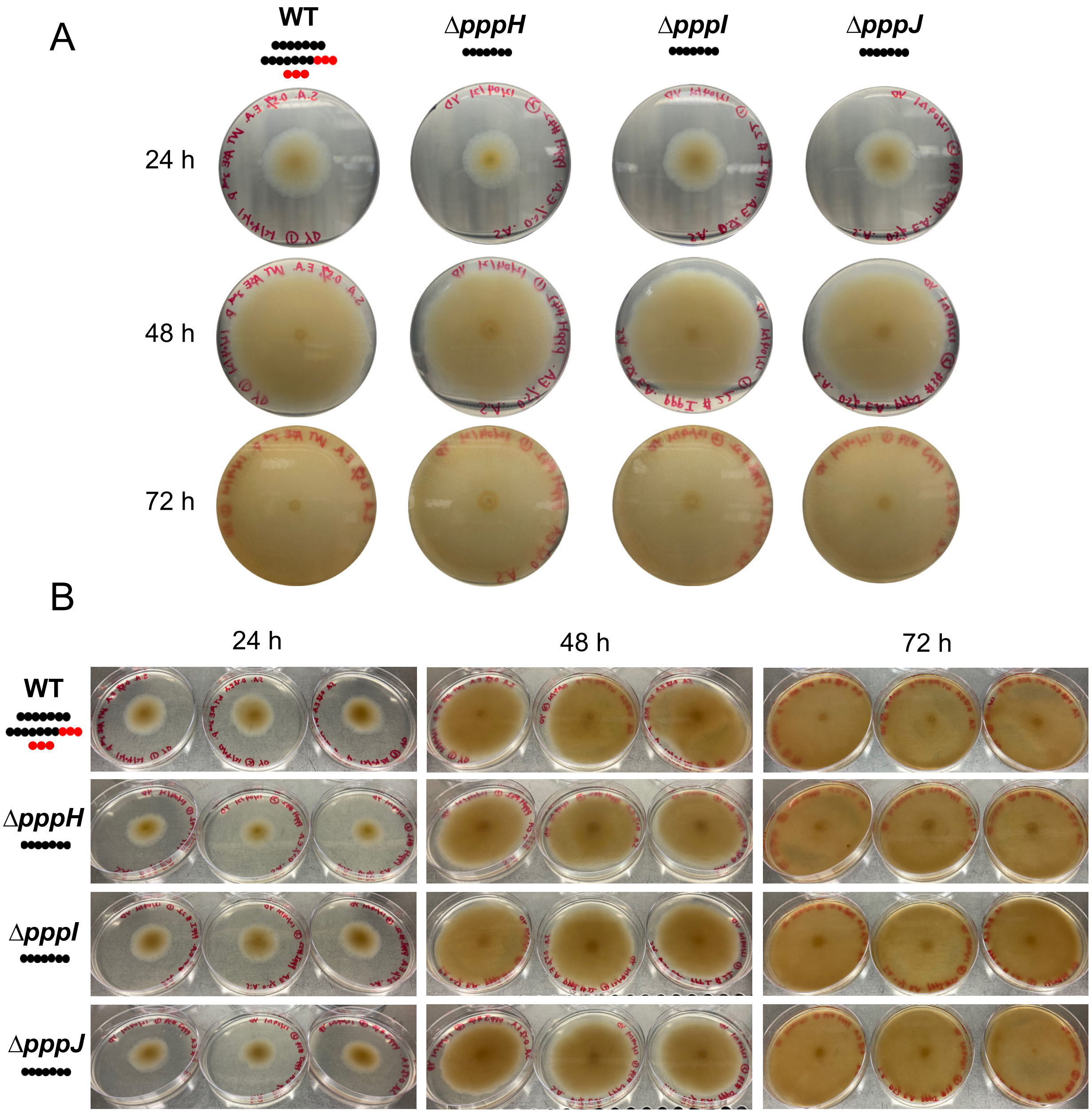

**Figure S41. Swarming assay results of Δ*pppH,* Δ*pppI,* and Δ*pppJ* mutants compared to the wild type.** The swarming assay was performed as described under Methods. Pictures shown were taken at 24, 48, and 72 hours after inoculation. (**A**) Representative results are shown. Pseudovibriamides are represented by beads: PA, seven black beads; PB, seven black beads and three red beads; PC, three red beads. (**B**) Triplicates of each strain are shown.

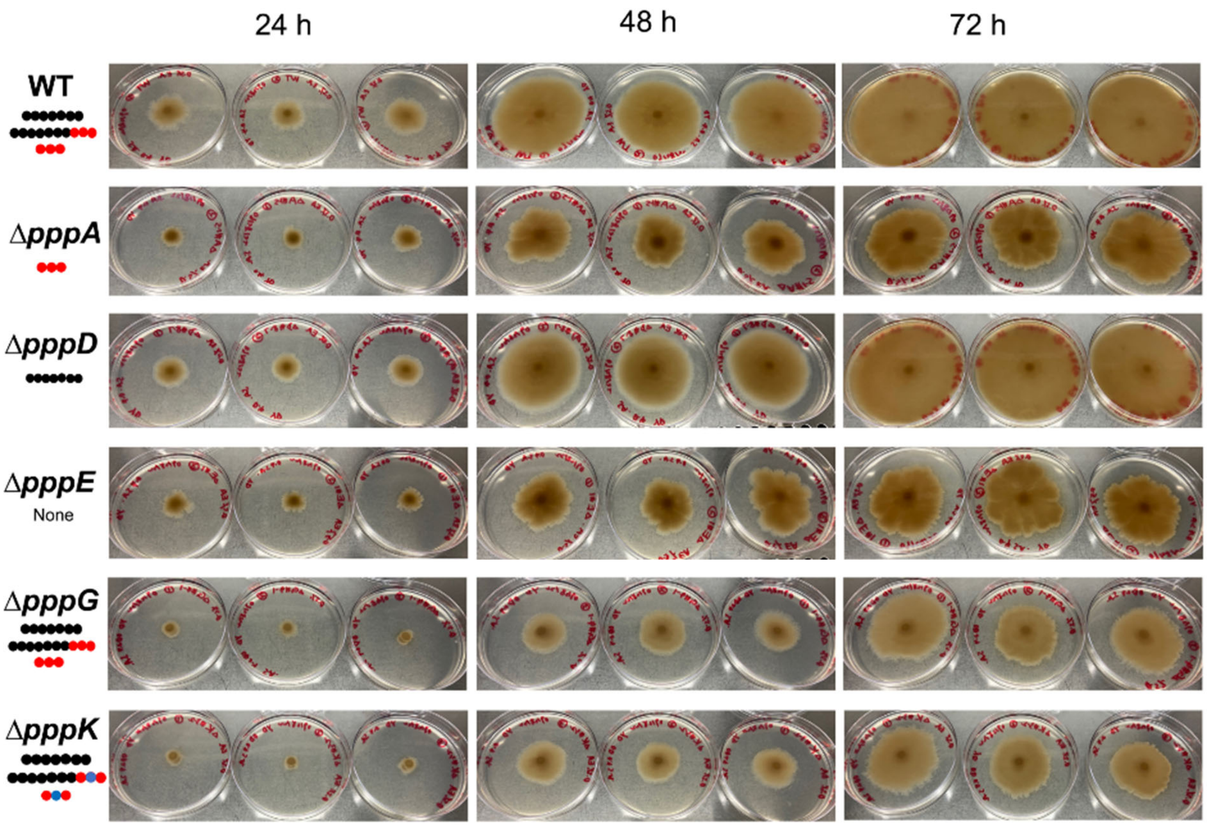

**Figure S42. Triplicate swarming assay results of Δ*pppA,* Δ*pppD,* Δ*pppE,* Δ*pppG,* and Δ*pppK* mutants compared to the wild type.** The swarming assay was performed as described under Methods. Pictures shown were taken at 24, 48, and 72 hours after inoculation. Pseudovibriamides are represented by beads: PA, seven black beads; PB, seven black beads and three red beads; PC, three red beads; dPB, seven black beads and two red beads with one blue bead in the middle; and dPC, two red beads with one blue bead in the middle. Triplicates of each strain are shown.

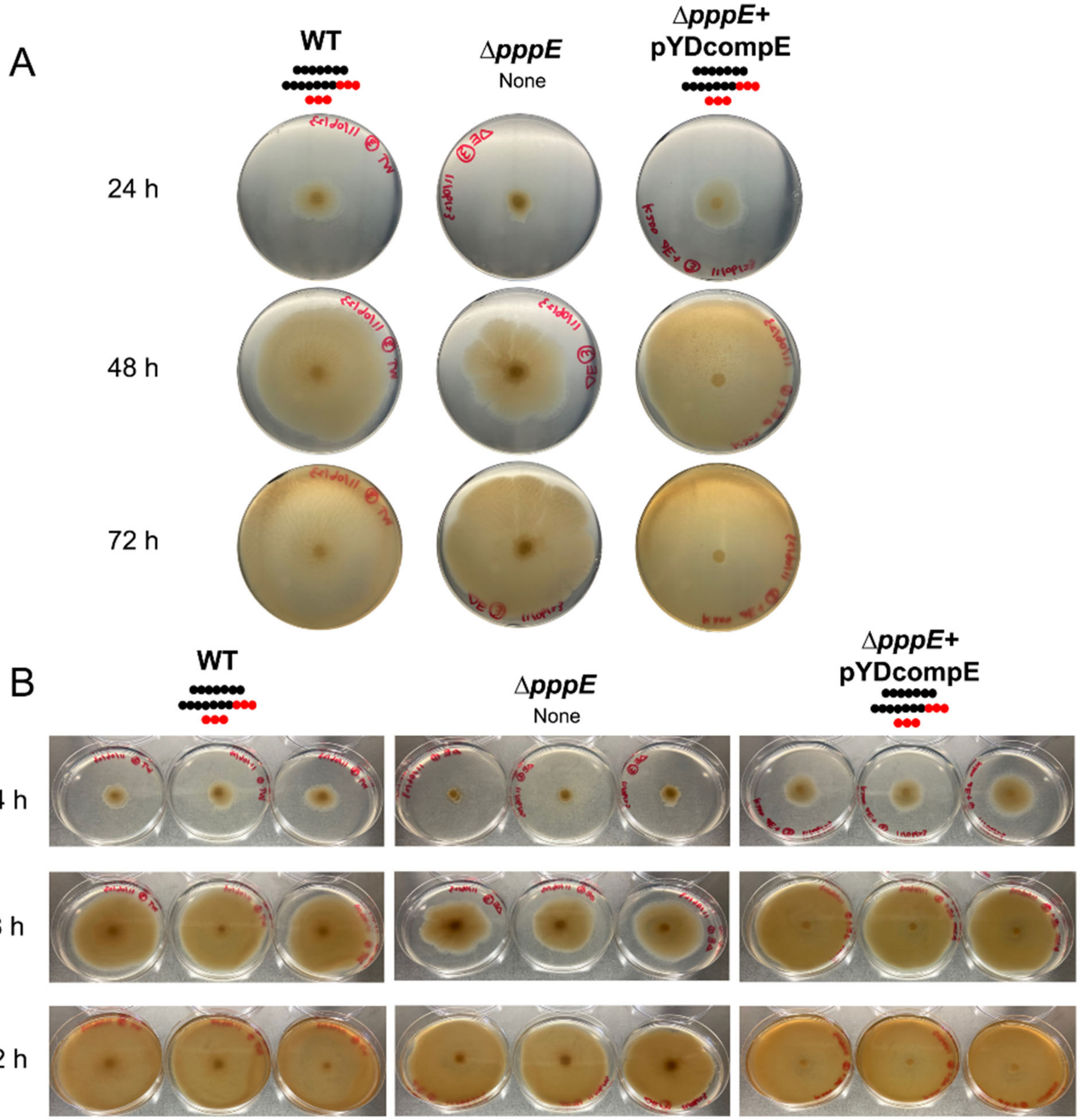

**Figure S43. Swarming assay results of Δ*pppE* mutant compared to the wild type and the genetically complemented strain.** The swarming assay was performed as described under Methods. Pictures shown were taken at 24, 48, and 72 hours after inoculation. (**A**) Representative results are shown. Pseudovibriamides are represented by beads: PA, seven black beads; PB, seven black beads and three red beads; PC, three red beads. Δ*pppE*+pYDcompE, complemented Δ*pppE* mutant using plasmid pYDcompE. (**B**) Triplicates of each strain are shown.

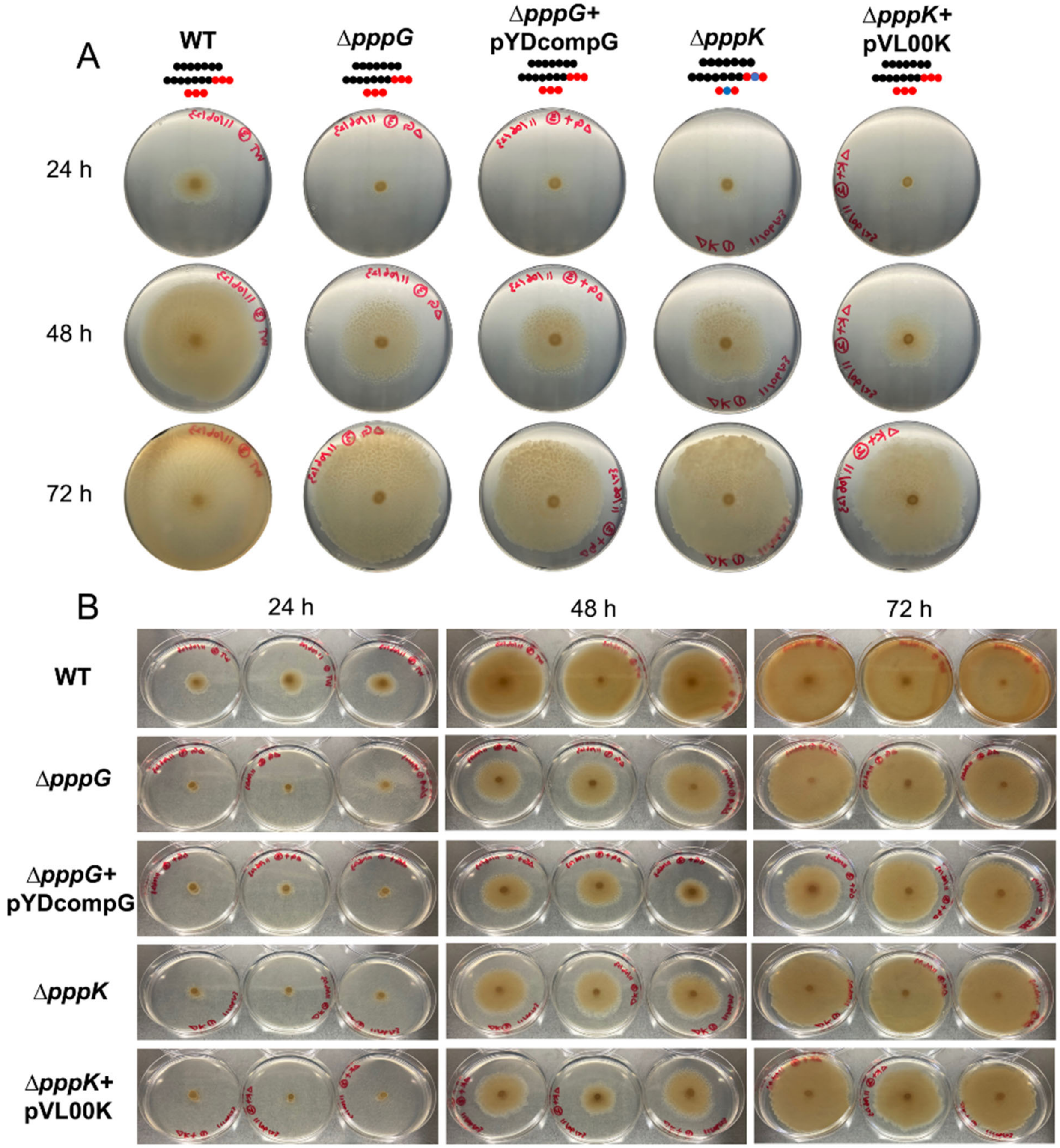

**Figure S44. Swarming assay results of Δ*pppG* and Δ*pppK* mutants compared to the wild type and genetically complemented strains.** The swarming assay was performed as described under Methods. Pictures shown were taken at 24, 48, and 72 hours after inoculation. (**A**) Representative results are shown. Pseudovibriamides are represented by beads: PA, seven black beads; PB, seven black beads and three red beads; PC, three red beads; dPB, seven black beads and two red beads with one blue bead in the middle; dPC, two red beads with one blue bead in the middle; Δ*pppK*+pVL00K, trans-complemented Δ*pppK* mutant using plasmid pVL00K; Δ*pppG*+pYDcompG, trans-complemented Δ*pppG* mutant using plasmid pYDcompG. (**B**) Triplicates of each strain are shown.

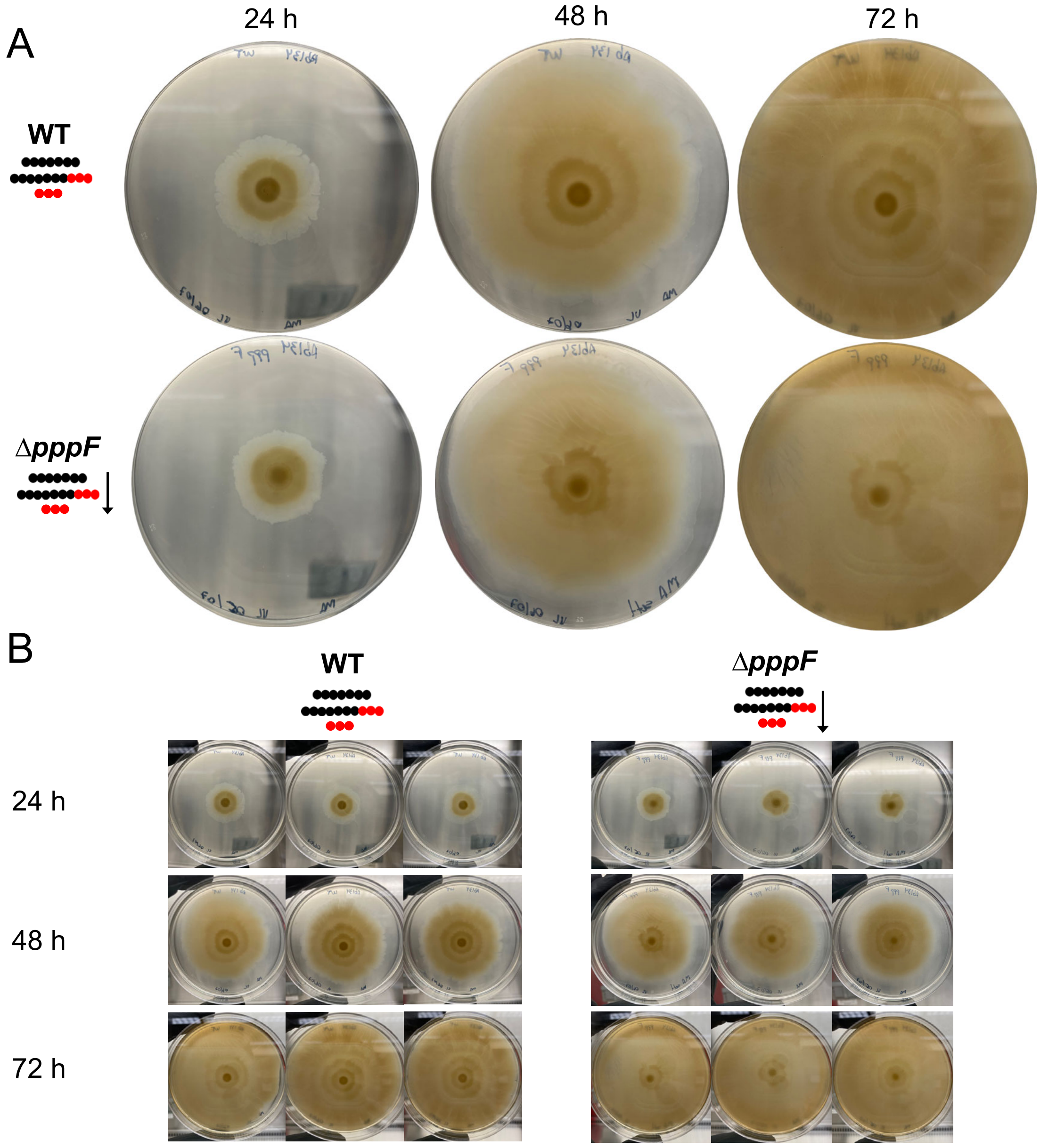

**Figure S45. Swarming assay results of Δ*pppF* mutant compared to the wild type.** The swarming assay was performed as described under Methods. Pictures shown were taken at 24, 48, and 72 hours after inoculation. (**A**) Representative results are shown. Pseudovibriamides are represented by beads: PA, seven black beads; PB, seven black beads and three red beads; PC, three red beads. (**B**) Triplicates of each strain are shown. The downward arrow means less pseudovibriamide produced.

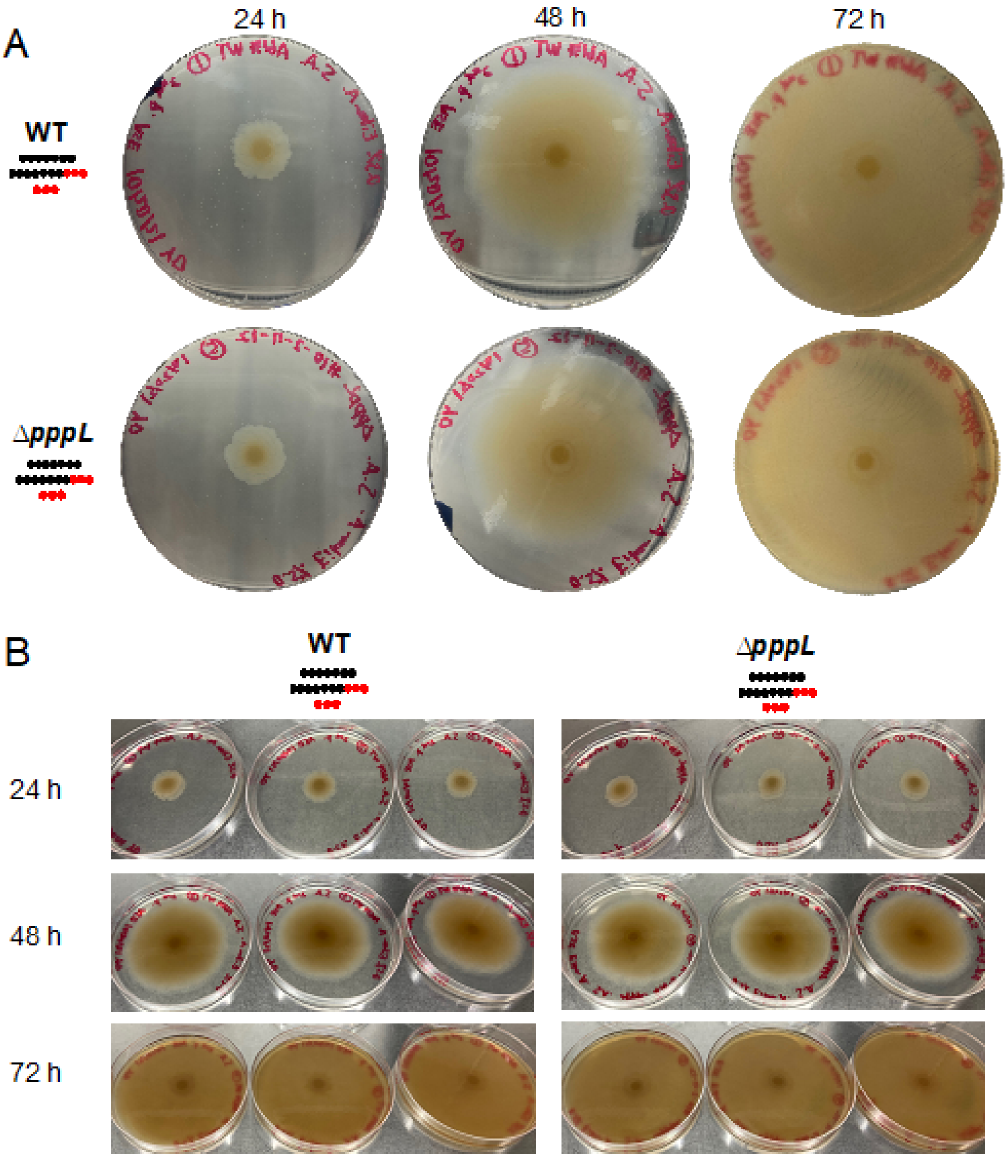

**Figure S46. Swarming assay results of Δ*pppL* mutant compared to the wild type.** The swarming assay was performed as described under Methods. Pictures shown were taken at 24, 48, and 72 hours after inoculation. (**A**) Representative results are shown. Pseudovibriamides are represented by beads: PA, seven black beads; PB, seven black beads and three red beads; PC, three red beads. (**B**) Triplicates of each strain are shown.

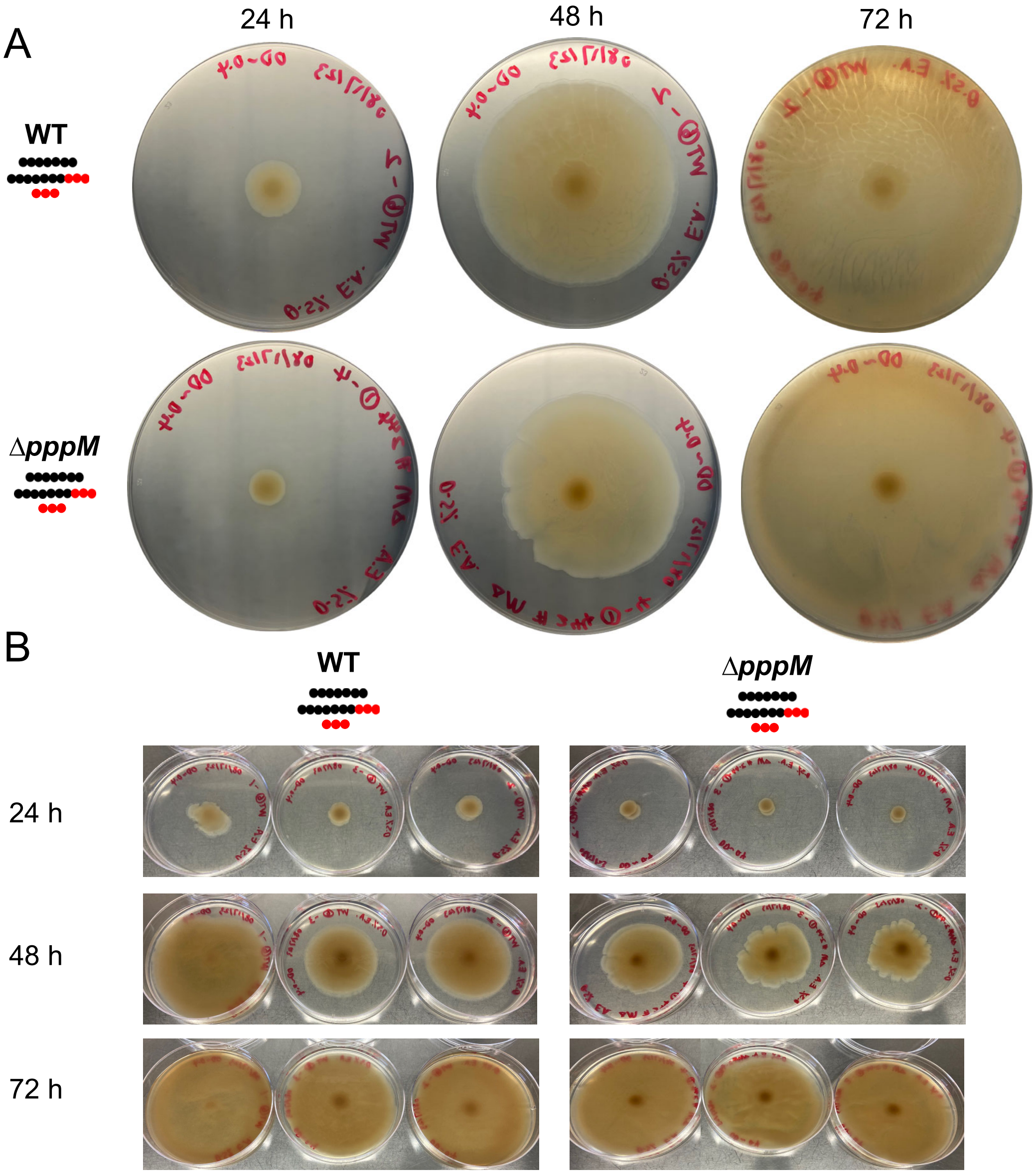

**Figure S47. Swarming assay results of Δ*pppM* mutant compared to the wild type.** The swarming assay was performed as described under Methods. Pictures shown were taken at 24,

48, and 72 hours after inoculation. (**A**) Representative results are shown. Pseudovibriamides are represented by beads: PA, seven black beads; PB, seven black beads and three red beads; PC, three red beads. (**B**) Triplicates of each strain are shown.

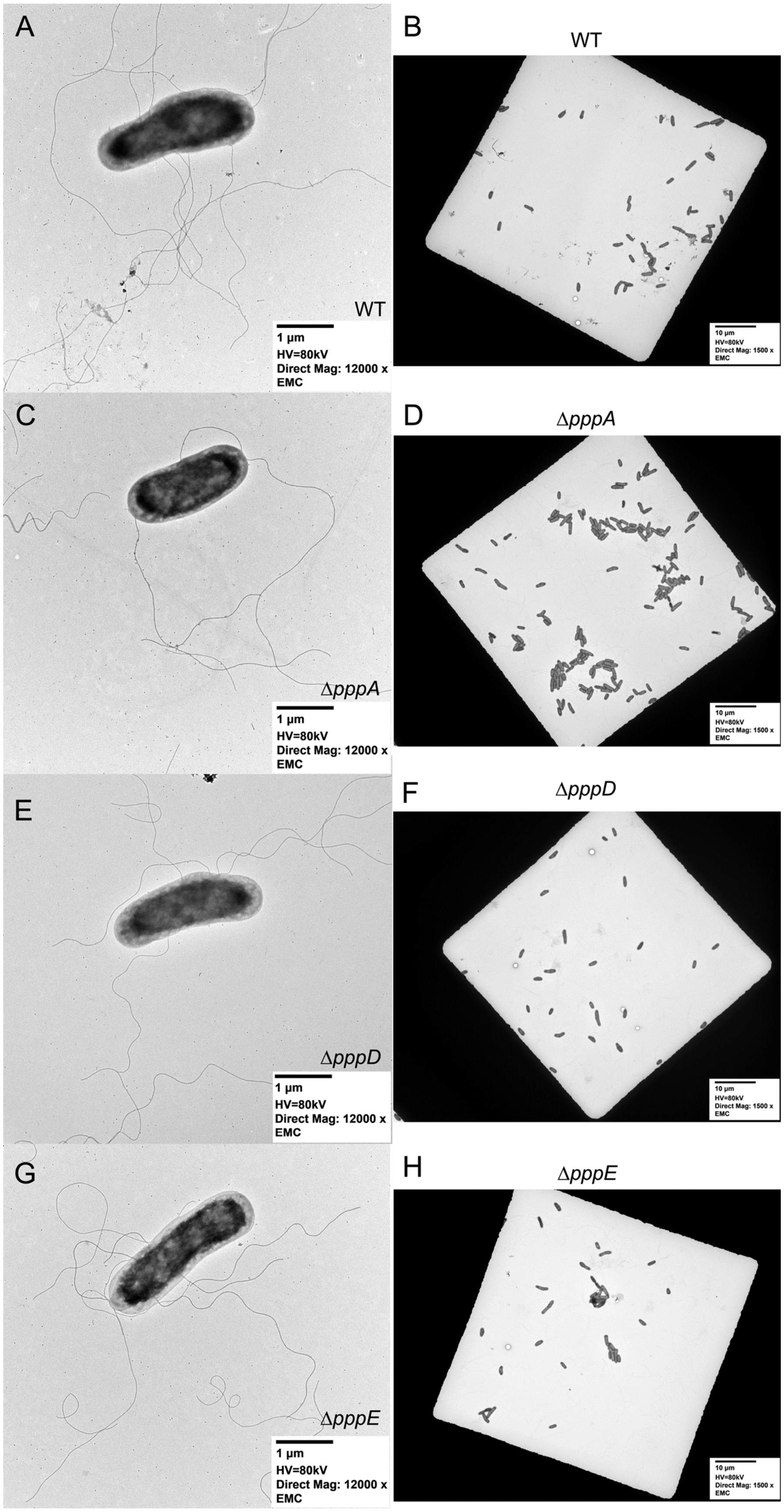
A
WT
1 µm
HV=80kV
Direct Mag: 12000 x
EMC
B
WT
10 µm
HV=80kV
Direct Mag: 1500 x
EMC
C
ΔpppA
1 µm
HV=80kV
Direct Mag: 12000 x
EMC
D
ΔpppA
10 µm
HV=80kV
Direct Mag: 1500 x
EMC
E
ΔpppD
1 µm
HV=80kV
Direct Mag: 12000 x
EMC
F
ΔpppD
10 µm
HV=80kV
Direct Mag: 1500 x
EMC
G
ΔpppE
1 µm
HV=80kV
Direct Mag: 12000 x
EMC
H
ΔpppE
10 µm
HV=80kV
Direct Mag: 1500 x
EMC

**Figure S48. Transmission Electron Microscopy of wild type and mutants.** Images of single cells and individual mesh windows, respectively of (**A**) and (**B**) wild type; (**C**) and (**D**) $\Delta$*pppA* mutant; (**E**) and (**F**) $\Delta$*pppD* mutant; (**G**) and (**H**) $\Delta$*pppE* mutant. Each TEM grid has 300 mesh windows in total. Negative staining with 1% phosphotungstic acid was used. HV, high voltage; EMC, Electron Microscopy Core.

1027

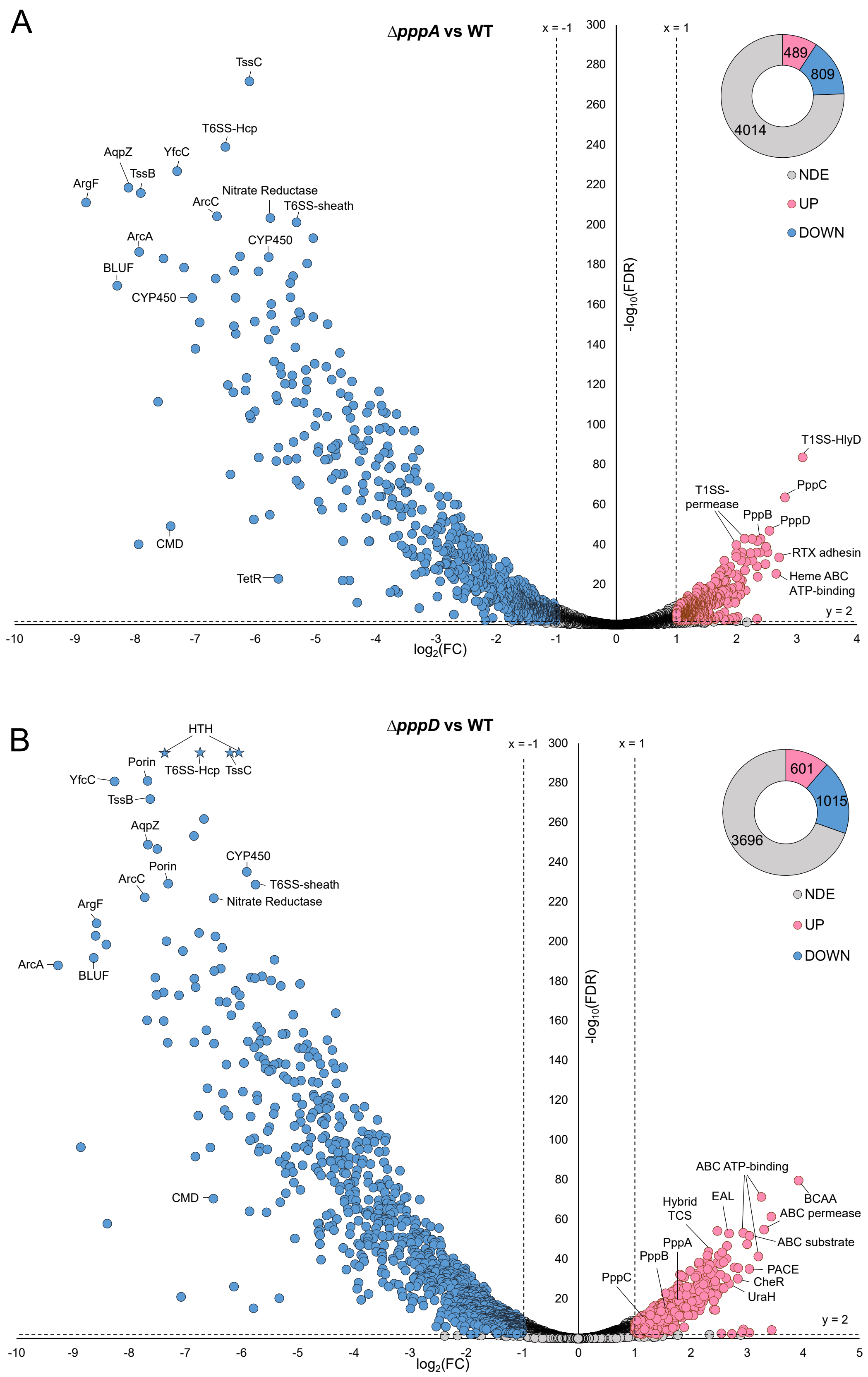
A
ΔpppA vs WT
x = -1
x = 1
y = 2
TssC
T6SS-Hcp
AqpZ
YfcC
TssB
ArgF
ArcC
Nitrate Reductase
T6SS-sheath
ArcA
CYP450
BLUF
CYP450
CMD
TetR
T1SS-HlyD
PppC
T1SS-permease
PppB
PppD
RTX adhesin
Heme ABC ATP-binding
489
809
4014
NDE
UP
DOWN
-log10(FDR)
log2(FC)
B
ΔpppD vs WT
x = -1
x = 1
y = 2
HTH
Porin
T6SS-Hcp
TssC
YfcC
TssB
AqpZ
Porin
CYP450
ArcC
T6SS-sheath
Nitrate Reductase
ArgF
ArcA
BLUF
CMD
ABC ATP-binding
BCAA
ABC permease
Hybrid TCS
EAL
ABC substrate
PppA
PppB
PACE
CheR
UraH
PppC
601
1015
3696
NDE
UP
DOWN
-log10(FDR)
log2(FC)

**Figure S49. Volcano plots of differentially expressed genes identified between Δ*pppA* and Δ*ppp*D mutants and the wild type using DESeq2.** (**A**) Differentially expressed genes of Δ*pppA* mutant compared to wild type. (**B**) Differentially expressed genes of Δ*pppD* mutant compared to wild type. FDR, false discovery rate or q-value; FC, fold-change; NDE, non-differential expression; UP, upregulated; DOWN, downregulated; Blue dots or blue donut portion, downregulated genes; Pink dots or pink donut portion, upregulated genes; and grey dots or grey donut portion, non-differentially expressed genes. The same filter (dotted lines) was applied to all differential expression analyses that is FC ≤ -2 (x = -1) or FC ≥ 2 (x = 1), and FDR ≤ 1% (y = 2). The total number of genes from each portion is listed in the donut chart. Products of outstanding downregulated or upregulated genes are labeled. HTH, helix-turn-helix transcriptional regulator; TssB, type VI secretion system contractile sheath small subunit; TssC, type VI secretion system contractile sheath large subunit; T6SS-sheath, type VI secretion system contractile sheath large subunit; T6SS-Hcp, type VI secretion system tube protein Hcp; YfcC, Arginine/ornithine antiporter; AqpZ, aquaporin Z; ArcA, arginine deiminase; ArcC, carbamate kinase; ArgF, ornithine carbamoyltransferase; BLUF, blue light using flavin domain; CYP450, cytochrome P450; T1SS-HlyD, HlyD family type I secretion periplasmic adaptor subunit; RTX adhesin, repeats-in-toxin adhesion protein; T1SS permease, type I secretion system permease/ATPase; CMD, carboxymuconolactone decarboxylase; TetR, TetR/AcrR family transcriptional regulator; BCAA ABC permease, branched-chain amino acid ATP-binding cassette transporter permease; ABC ATP-binding, ATP-binding cassette transporter ATP-binding protein; ABC substrate, ATP-binding cassette transporter substrate binding protein; Heme ABC ATP-binding, heme ATP-binding cassette transporter ATP-binding protein; EAL, EAL domain containing protein; Hybrid TCS, hybrid sensor histidine kinase/response regulator; PACE, the proteobacterial antimicrobial compound efflux transporter; CheR, protein-glutamate O-methyltransferase; and UraH, hydroxyisourate hydrolase. Some outstanding dots are unlabeled as they are hypothetical proteins. Stars represent DE genes with too large values of -log10(FDR), which were rounded to 1000.

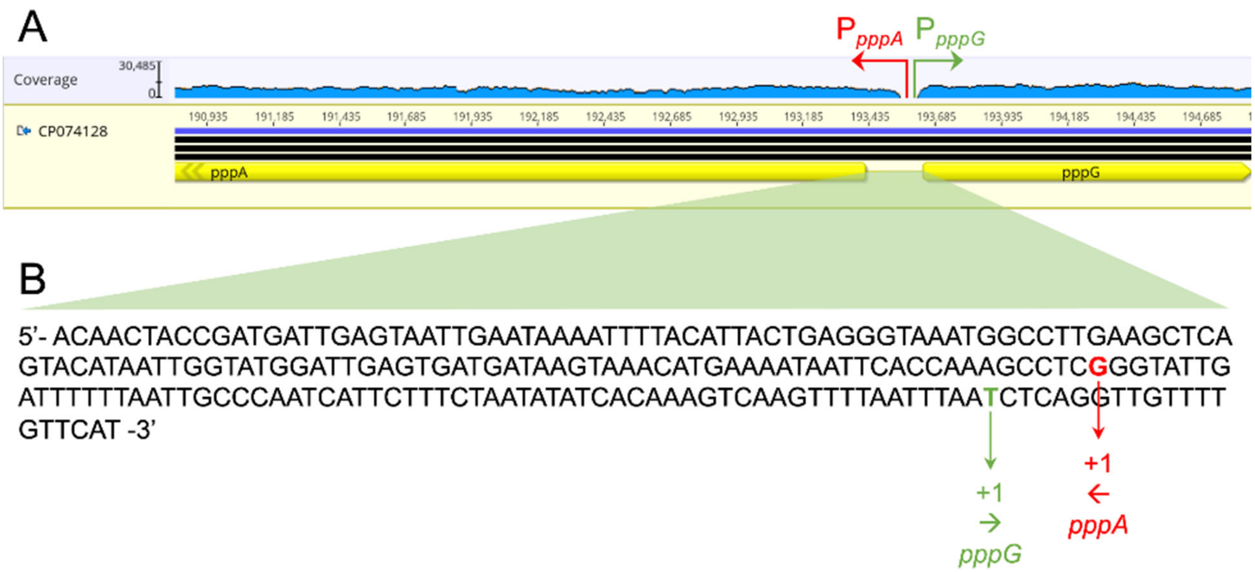

**Figure S50. Identification of $P_{pppA}$ based on RNA-Seq data.** (**A**) RNA-Seq reads mapped to the reference genome. The gap in coverage indicates the presence of divergent promoters for *pppA* and *pppG*. (**B**) Sequence of promoter region highlighting the predicted +1 of transcription of *pppG* and +1 of transcription of *pppA* according to RNA-seq data.

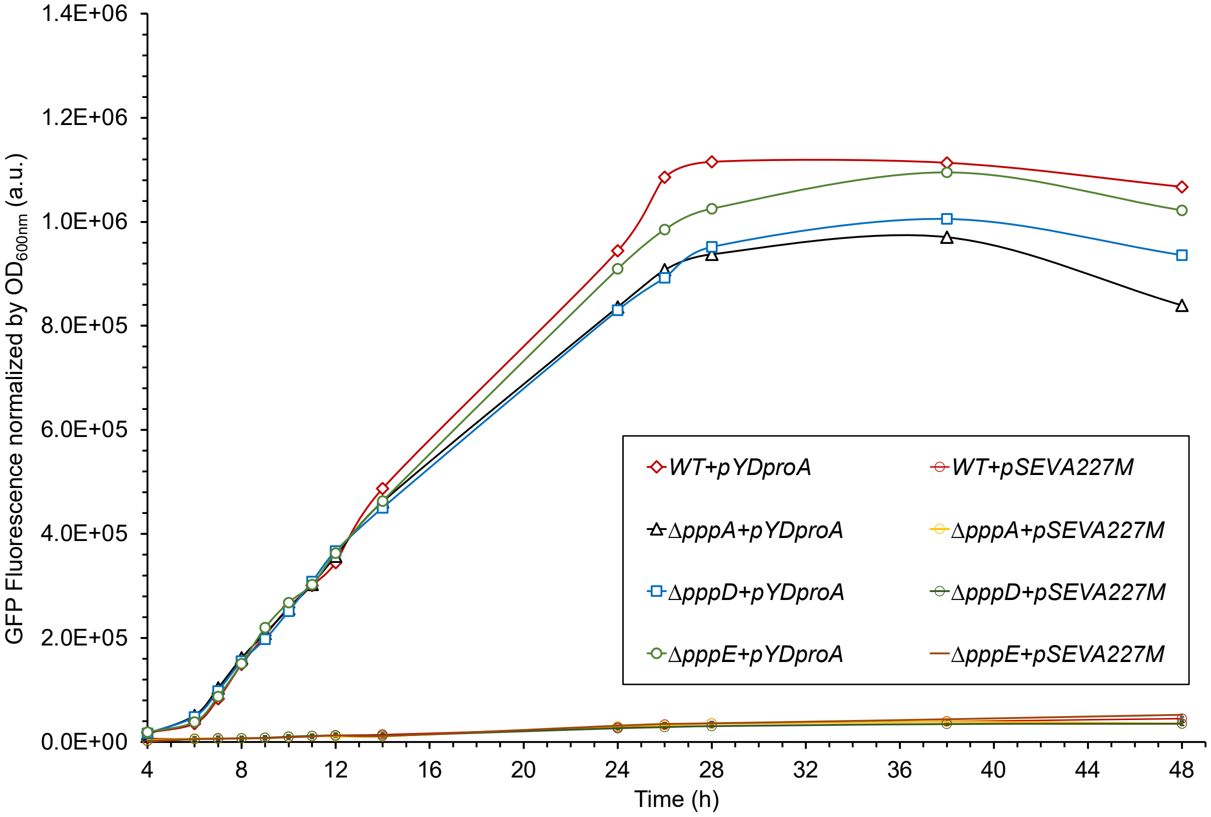

**Figure S51. $P_{pppA}$ autoregulation assay.** Readings of GFP fluorescence level were normalized by $OD_{600nm}$ and plotted against incubation time of 48 hours. pYDproA, GFP under $P_{pppA}$; pSEVA227M, promoterless GFP. *N*=12. Error bars indicate standard deviation.

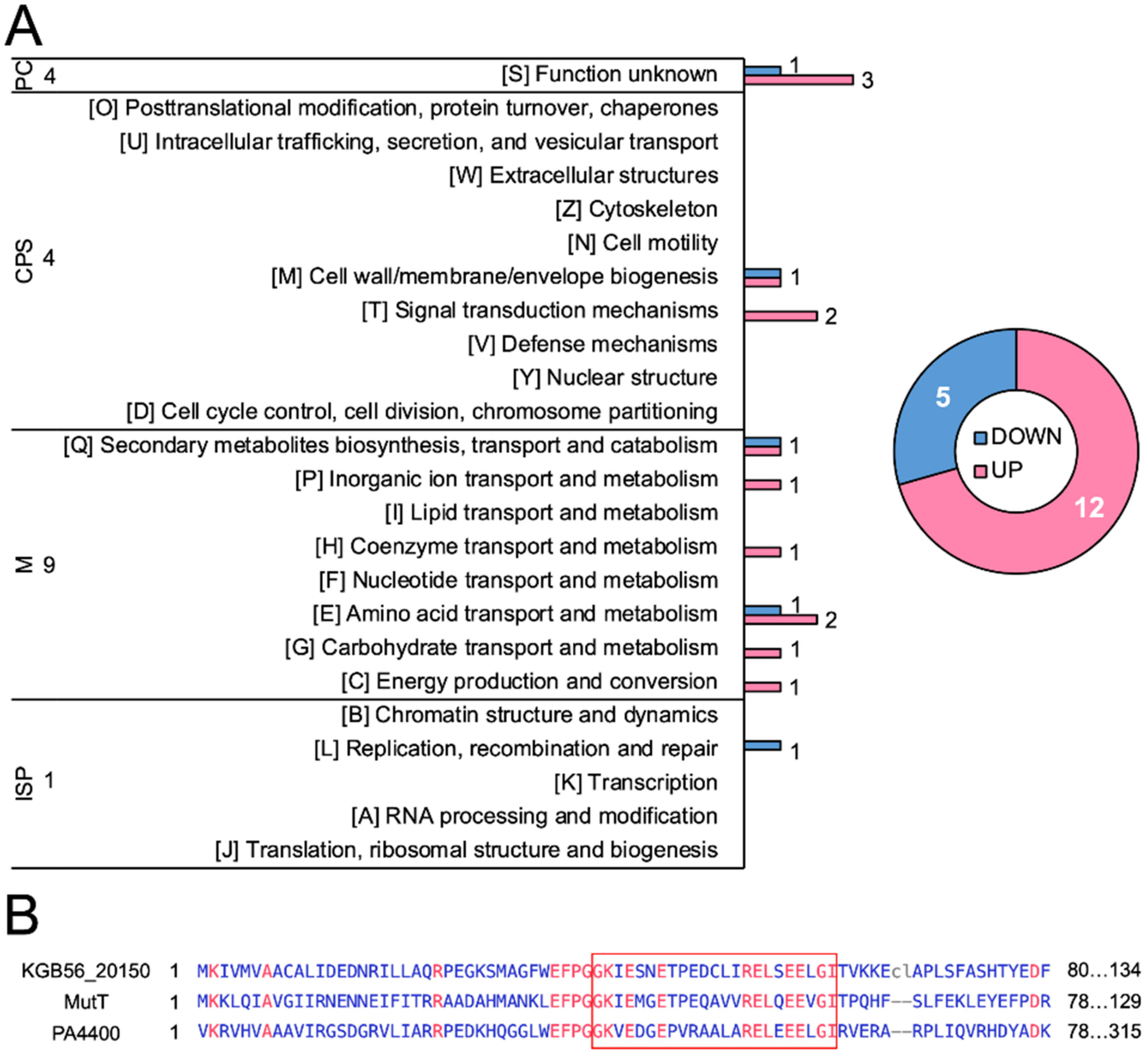

**Figure S52. COG classification of DE genes in ∆*pppA/E* mutants filtered using NDE genes of ∆*pppD* mutant compared to the wild type.** (**A**) The distribution of 24 upregulated genes and 5 downregulated genes in each COG category. The donut graph shows the total number of upregulated or downregulated genes. Pink, upregulated; Blue, downregulated. PC, poorly characterized; CPS, cellular processes and signaling; M, metabolism; and ISP, information storage and processing. The total number of genes in each category is listed. (**B**) COBALT was used to align amino acid sequences using Ab134 MutT as the anchor sequence. *P. brasiliensis*

Ab134 KGB56_20150 (QUS55589.1); *E. coli* strain K12 MutT (CAD6022180.1); and *P. aeruginosa* PAO1 PA4400 (NP_253090.1). Red color denotes conserved residues. Blue color denotes not conserved residues. The red box highlights the conserved Nudix motif, G-X5-E-X7-R-E-U-X-E-E-X-G-U, where X could be any amino acid, and U should be a hydrophobic amino acid.

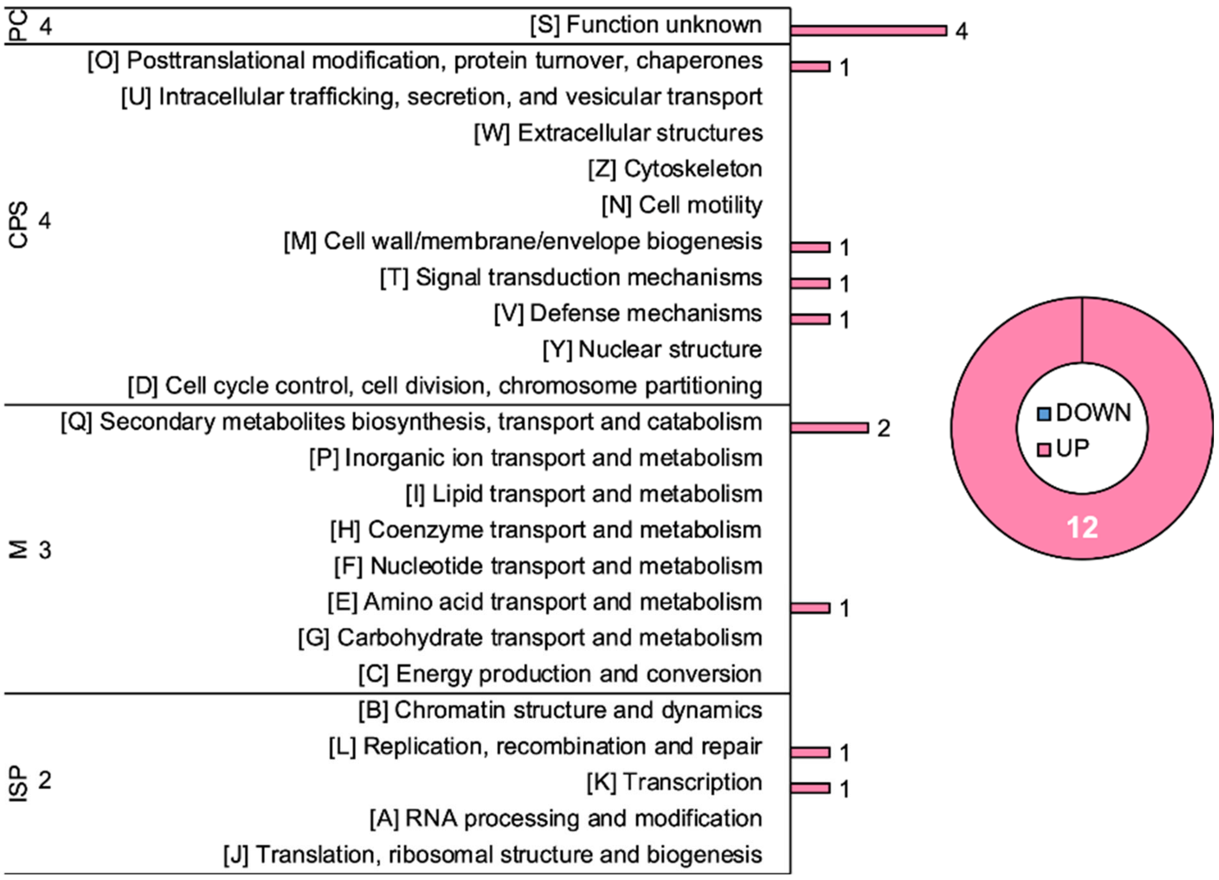

**Figure S53. COG classification of DE genes reversely regulated in Δ*pppA/E* mutants and the Δ*pppD* mutant compared to the wild type.** Donut graph shows the total number of upregulated or downregulated genes. Pink color, upregulated; Blue color, downregulated; PC, poorly characterized; CPS, cellular processes and signaling; M, metabolism; and ISP, information storage and processing. The total number of genes in each category is listed.

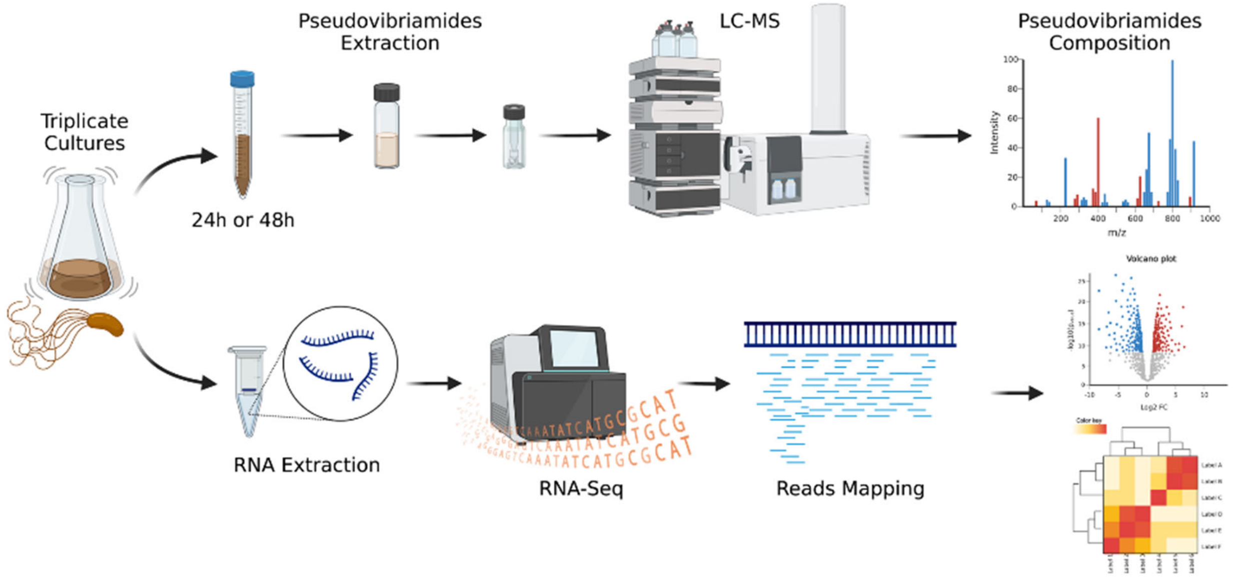

**Figure S54. Workflow from bacterial culture to transcriptomic analysis.**
